# Myeloid cell influx into the colonic epithelium is associated with disease severity and non-response to anti-Tumor Necrosis Factor Therapy in patients with Ulcerative Colitis

**DOI:** 10.1101/2023.06.02.542863

**Authors:** Divya Jha, Zainab Al-Taie, Azra Krek, Shadi Toghi Eshghi, Aurelie Fantou, Thomas Laurent, Michael Tankelevich, Xuan Cao, Hadar Meringer, Alexandra E Livanos, Minami Tokuyama, Francesca Cossarini, Arnaud Bourreille, Regis Josien, Ruixue Hou, Pablo Canales-Herrerias, Ryan C. Ungaro, Maia Kayal, James Marion, Alexandros D Polydorides, Huaibin M. Ko, Darwin D’souza, Raphael Merand, Seunghee Kim-Schulze, Jason A. Hackney, Allen Nguyen, Jacqueline M. McBride, Guo-Cheng Yuan, Jean Frederic Colombel, Jerome C. Martin, Carmen Argmann, Mayte Suárez-Fariñas, Francesca Petralia, Saurabh Mehandru

## Abstract

Ulcerative colitis (UC) is an idiopathic chronic inflammatory disease of the colon with sharply rising global prevalence. Dysfunctional epithelial compartment (EC) dynamics are implicated in UC pathogenesis although EC-specific studies are sparse. Applying orthogonal high-dimensional EC profiling to a Primary Cohort (PC; n=222), we detail major epithelial and immune cell perturbations in active UC. Prominently, reduced frequencies of mature *BEST4*^+^*OTOP2*^+^ absorptive and *BEST2*^+^*WFDC2*^+^ secretory epithelial enterocytes were associated with the replacement of homeostatic, resident *TRDC^+^KLRD1^+^HOPX^+^* γδ^+^ T cells with *RORA^+^CCL20^+^S100A4^+^* T_H17_ cells and the influx of inflammatory myeloid cells. The EC transcriptome (exemplified by *S100A8, HIF1A, TREM1, CXCR1*) correlated with clinical, endoscopic, and histological severity of UC in an independent validation cohort (n=649). Furthermore, therapeutic relevance of the observed cellular and transcriptomic changes was investigated in 3 additional published UC cohorts (n=23, 48 and 204 respectively) to reveal that non-response to anti-Tumor Necrosis Factor (anti-TNF) therapy was associated with EC related myeloid cell perturbations. Altogether, these data provide high resolution mapping of the EC to facilitate therapeutic decision-making and personalization of therapy in patients with UC.

## Main

Ulcerative colitis (UC) involves epithelial barrier dysfunction in combination with immune dysregulation that leads to chronic inflammation of the large intestines^1^. In addition to reinforcing the physical barrier, the intestinal epithelium regulates nutrient uptake and facilitates tissue homeostasis^2,3^. Murine studies have delineated bi-directional epithelial-intraepithelial T cell interactions^4^, as being critical for response to infection^5^, nutrient sensing^6^, and regulation of gut inflammation^7^. Additionally, epithelium-proximate myeloid cells, especially macrophages regulate epithelial stem cells^8–10^, epithelial viability^11^ and host-microbiota interactions^12^ to reinforce the epithelial-barrier function. Despite their critical role in tissue homeostasis, human intestinal epithelium-associated myeloid cells are poorly defined.

The therapeutic landscape of UC has significantly expanded with the advent of therapies targeting tumor necrosis factor (TNF)^13^, integrin α4β7^14^, IL-12/23 pathways^15^, JAK-STAT pathways^16, 17^ and sphingosine 1 phosphate (S1P) receptors^18^, with additional therapeutic options poised to become available in the near future^19^. However, approximately 30% of patients are non-responsive to initial therapy and an additional 50% become secondary non-responders^20^. To break this ‘therapeutic ceiling’, a ‘precision medicine-like’, drug-mechanism and biomarker-based approach is suggested. However, this strategy is hampered by the lack of mechanistic data to guide therapeutic decisions^21^. High dimensional approaches such as single cell RNA sequencing (scRNA-seq) have enabled profiling of intestinal immune cells at unprecedented resolution^22–28^. However, prior studies have focused on whole intestinal tissues or on the intestinal lamina propria (LP) and to-date, dedicated epithelial compartment-focused studies that relate epithelial-immune perturbations to therapeutic outcomes in patients with UC are lacking.

Here, using bulk RNA sequencing, scRNA-seq, Spatial Transcriptomics (ST), and complemented by relevant protein-level assessments, we have focused on epithelium-specific immune and non-immune alterations in patients with UC. Further, we have defined epithelium-specific pathotypes that associate with non-response to TNF inhibitors (TNFi). Taken together, our study comprehensively evaluates the role of EC in active UC and provide a rational basis for therapeutic choices in patients who are non-responsive to TNF-inhibition.

## Results

### Intraepithelial immune compartment is extensively remodeled in active UC

Bulk RNA sequencing was performed on ethylenediaminetetraacetic acid (EDTA)-dissociated epithelial fraction (Epi_bulk_) or EDTA-dissociated, percoll-enriched intraepithelial immune cells (IIC, IIC_bulk_) using biopsies derived from histologically uninflamed and inflamed colon of subjects with UC or from Normal Volunteers (NV; Fig. 1a, Extended Data Fig. 1, Methods, Supplementary Table 1). Inflamed UC samples (red) clustered distinctly from uninflamed UC (green) and NV (blue) samples for both Epi_bulk_ and IIC_bulk_ (Fig. 1b). We identified differentially expressed (DEGs) between inflamed, non-inflamed and NV within EPI and IIC (DEGs; false discovery rate (FDR) <0.05; fold change (FCH) >2.0) (Fig. 1b, Supplementary Table 2). In contrast, no significant DEGs were detected in either Epi_bulk_ or IIC_bulk_ when comparing uninflamed UC and NV samples. When comparing the Epi_bulk_ and IIC_bulk_ inflammation signatures (hereby refer to as IIC_inf_ and Epi_inf_ signatures respectively), significant overlap was noted, with 989 DEGs (562 shared, up- and 427 down-regulated genes, Fisher’s exact test p<2.2×10^-16^ for both; Fig. 1d, Supplementary Table 2a,b). Next, to characterize the cell types associated with the transcriptome of Epi_inf_ (Epi_bulk_-derived DEGs, inflamed vs. uninflamed) and IIC_inf_ (IIC_bulk_-derived DEGs, inflamed vs. uninflamed), we performed an over-representation analysis (ORA) using a curated scRNA-seq derived atlas for cell types in the human colonic tissues^23^. Up-regulated Epi_inf_ and IIC_inf_ genes were significantly (adjP<0.01) enriched for monocytes, macrophage, dendritic cells, T cells and plasma cells suggesting an infiltration of immune cells in the inflamed epithelia while down-regulated Epi_inf_ and IIC_inf_ DEGs were enriched for epithelial cells (Fig. 1e). Consistent with the cell type enrichments, ORA of KEGG^29^ pathways revealed that the up-regulated Epi_inf_ and IIC_inf_ DEGs were involved in several inflammation-associated pathways notable for IL-17 signaling, T_H17_ cell differentiation, cytokine-cytokine receptor interactions, antigen presentation and immunoglobulin production, while the down-regulated Epi_inf_ and IIC_inf_ genes were found to be predominantly associated with pathways such as retinol and butanoate metabolic pathways indicative of epithelial cell dysfunction in active UC (Fig. 1f, Supplementary Table 2c). A subset of the Epi_inf_ and IIC_inf_ DEGs, found in the intersection of Fig.1d and associated with these pathways, is depicted in Fig. 1g. Notable for the upregulation of immune cell related transcripts are *OSM, IL1B, CXCL8, CCL24, IL17A CD4, SAA1, FOXP3, CD14*, *MRC1, MNDA, LYZ, TREM1, CSF3R*, *IGHG and JCHAIN,* while epithelial cell-expressed genes including *PRAP1*, *AQP7*, *GUCA2A, DEFB1, RARB, SYNPO* and innate immune system-related nociceptor gene *TRPV1* were significantly down-regulated ^30–34^ (Fig. 1g, Supplementary Table 2d). Altogether, these data pointed to the influx of non-homeostatic inflammatory cells in the EC of inflamed UC, associated with the replacement of homeostatic epithelial programs with powerful pro-inflammatory programs.

**Fig. 1:**
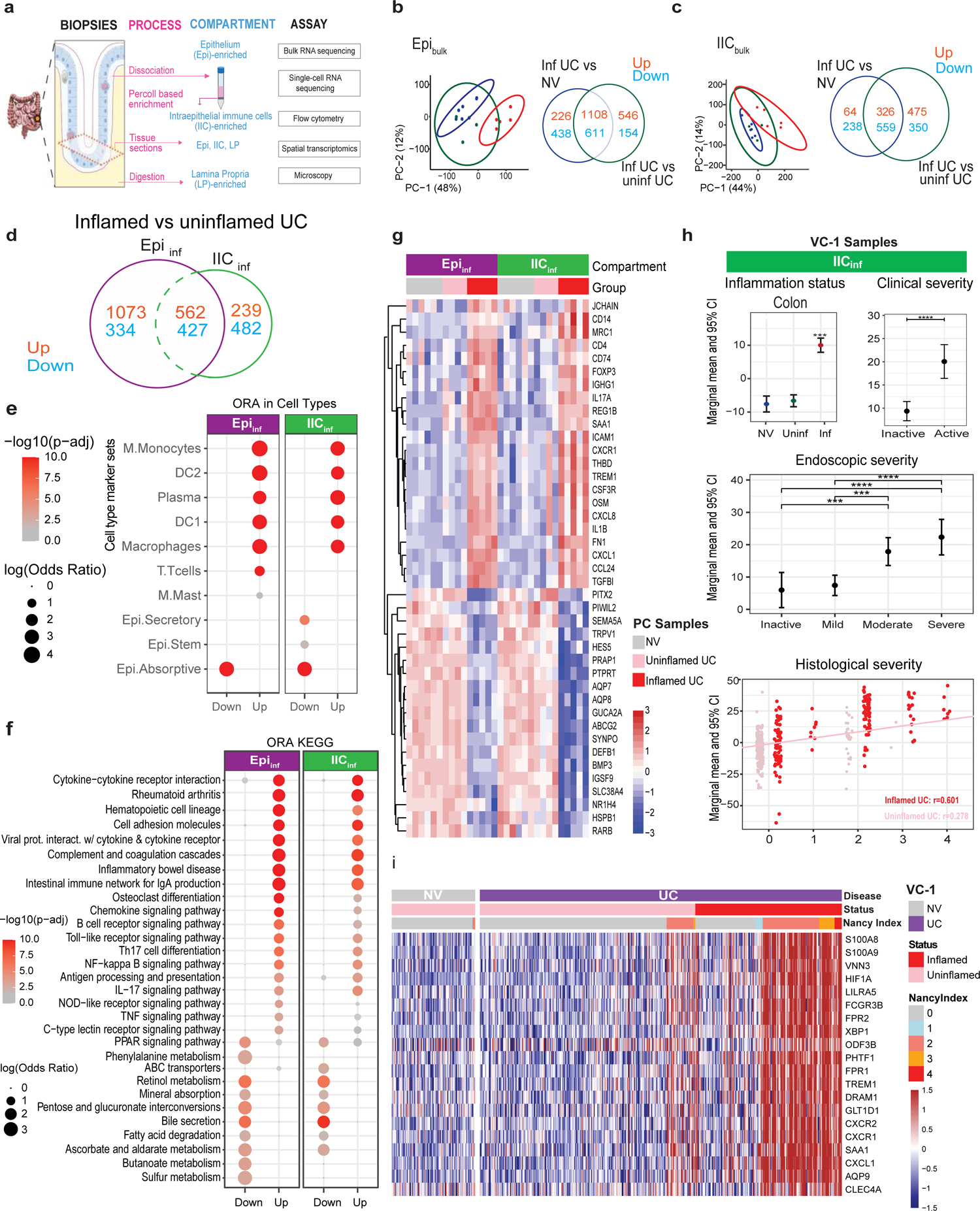
Intraepithelial immune compartment is extensively remodeled in active UC. **a**, Experimental design outlining the methodologies and assays used. **b,** *Left*; Principal component analysis (PCA) plot of the epithelial compartment (Epi_bulk_) expression profiles from the healthy (blue) (n=6), uninflamed (green) (n=4), and inflamed (red) (n=5) biopsies. *Right;* Venn diagram showing the differentially expressed genes (DEGs) from inflamed UC samples compared to NV samples (blue) and the DEGs from inflamed UC samples compared to the uninflamed UC samples (green). Up-regulated DEGs are in red while down-regulated DEGs in blue. **c**, *Left*; PCA plot of the intraepithelial immune cells (IIC_bulk_) RNA-seq expression profiles of the samples from the IIC from the healthy (blue) (n=6), uninflamed (green) (n=4), and inflamed (red) (n=5) biopsies. *Right;* Venn diagram showing the DEGs from inflamed UC samples compared to NV samples (blue) and the DEGs from inflamed UC samples compared to the uninflamed UC samples (green). Up-regulated DEGs are in red while down-regulated DEGs are in blue. **d,** Venn diagram showing the intersection of inflamed vs uninflamed UC DEGs from the Epi_bulk_ and IIC_bulk_. **e-f,** Overexpression analysis (ORA) of DEGs between inflamed and uninflamed UC samples interrogated from the cell type signature^23^ for selected significantly (adj-p<0.05) enriched cell types (e) and KEGG^29^ pathways (f). Color intensity represents -log10 transformed p-adjusted and the size represents the log2 transformed Odds Ratio (OR) using Fisher’s exact test. OR from enrichments with <5 IIC_inf_ DEG are not presented. **g,** Heatmap depicting expression profiles of 42 DEGs selected for relevance with disease pathogenesis among the DEGs in the intersection of Fig.1d. **h,** Association of the IIC_inf_ signature with UC disease and severity in the validation cohort-1 (VC-1). Each plot in **h** represents the estimated marginal mean expression and 95% CI for the GSVA-scores of the up-regulated genes in the IIC_inf_ signature across conditions. *Upper left*; Activity of IIC_inf_ signature in endoscopically inflamed colon compared to uninflamed biopsies of IBD patients and non-IBD normal volunteers (NV) from VC-1 cohort using fora function from fgsea R package. *Upper right*; The mean activity of up-regulated DEGs from IIC_inf_ signature was compared between clinically active IBD patients compared to IBD patients with no active disease and was also associated with endoscopic scores. *Lower;* Activity of IIC_inf_ up-regulated genes was associated with histological scores (measured by Nancy score 0-4) of endoscopically inflamed biopsies (red) and uninflamed biopsies (blue) from UC patients using Pearson’s correlation test *p ≤ 0.05; **p ≤ 0.01; ***p ≤ 0.001; ****p ≤ 0.0001 (ic test and BH-adjusted p-value). **i,** Heatmap showing the expression of the top 20 genes (among DEGs in inflamed versus uninflamed or NV comparisons) correlated with histological disease severity (Nancy index 0-4) measured from endoscopically inflamed gut biopsies of UC patients in VC-1 cohort.

Having seen a significant overlap of Epi_inf_ and IIC_inf_ gene signatures, we studied the association of IIC_inf_ signature in an independent Validation Cohort (VC-1) consisting of 406 patients with UC and 243 NV with available bulk RNA-seq data on whole biopsies (Extended Data Fig. 1, Methods). Using Gene Set Variation Analysis (GSVA), we noted significantly higher expression of the up-regulated, primary cohort (PC)-derived IIC_inf_ genes in inflamed intestinal biopsies from VC-1 compared to NV and uninflamed UC biopsies (Fig. 1h, Extended Data Fig. 2a). Furthermore, the expression of up-regulated IIC_inf_ genes strongly associated with higher clinical and endoscopic disease severity and was positively correlated with histologic severity in inflamed biopsies from VC-1 (r=0.601, p=1.05*10^-16^ Fig. 1h, Extended Data Fig. 2b-d). The top genes that correlated with histologic severity (Nancy score^35^) included proinflammatory genes *SAA1*, *S100A8*, *S100A9*; myeloid cell-associated genes *TREM1, LILRA5*; neutrophil chemotactic genes *FPR1, FPR2, CXCL1, CXCR1, CXCR2, FCGR3B/CD16*; hypoxia related gene *HIF1A*; autophagy gene *DRAM1* and unfolded protein response gene *XBP1* (Fig. 1i)^36–39^.The findings from our EC-bulk transcriptomic data revealed a complex, myeloid cell-dominated milieu in UC-associated inflammation that significantly correlated with parameters of clinical, endoscopic, and histologic severity.

**Fig. 2:**
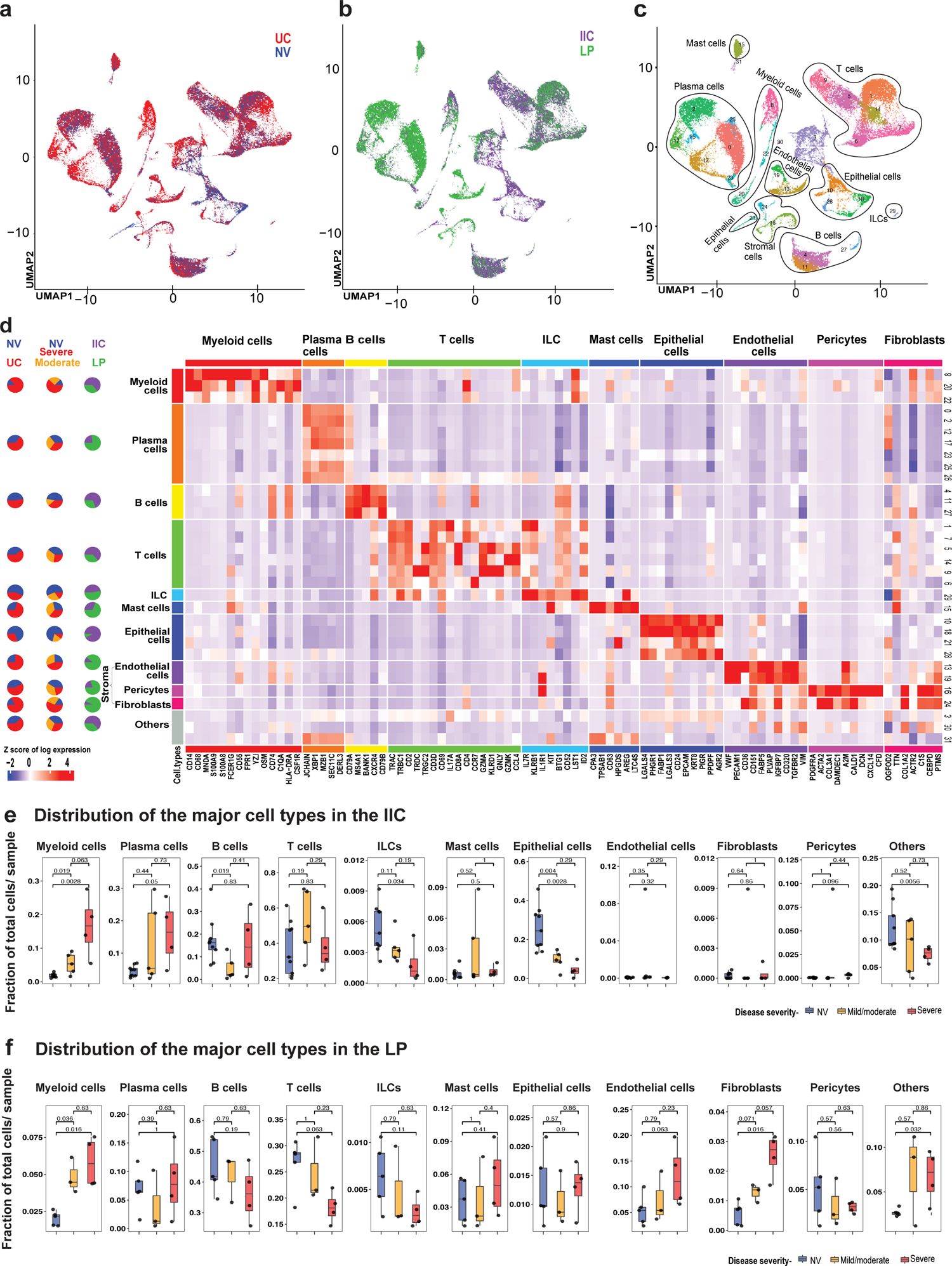
High-resolution mapping of immune cells reveals distinct cellular composition in the colonic EC of ulcerative colitis patients. **a-c**, UMAP of scRNA-seq data colored by **a,** disease status (i.e., UC samples (n=9) and NV samples (n=9)), **b,** intraepithelial immune cells (IIC) and lamina propria (LP) compartments, **c,** 32 cell-type annotated clusters identified by unsupervised clustering **d,** Heatmap showing the average z-score normalized log expression of canonical cell type markers across different clusters. Pie charts on the left indicate overall composition for each cell type, in terms of disease status, severity and tissue compartments. **e-f,** Boxplots of per-sample cell type fractions in IIC (e) and LP (f), stratified by disease status and severity. NV (blue), mild/moderate UC (orange) and severe UC samples (red). P-values from Wilcoxon signed-rank test are reported.

### High-resolution mapping of immune cells reveals distinct cellular composition in inflamed and uninflamed colonic epithelial compartment of UC patients

To further define UC-associated cellular and transcriptomic changes within the IIC (percoll-enriched EC), scRNA-seq was performed on 9 NV and 9 UC patients with active inflammation, stratified as mild-moderate (n=5) or severe (n=4) inflammation. In 5 NV and 7 UC patients, scRNA-seq was also performed on LP-derived mononuclear cells (methods) to capture compartment specific molecular and cellular perturbations between EC and LP from the same patient. We sequenced a total of 108,671 cells, with a mean of 7,698 unique molecular identifiers (UMIs) of 1437 unique genes per cell, on average (Supplementary Table 3a). Unsupervised clustering based on transcriptomic profile of scRNA-seq data resulted in 32 distinct cell clusters spanning across NV and UC (Fig. 2a), as well as IIC and LP (Fig. 2b). Grouping the clusters by expression profile similarities indicated the presence of 10 major cell types (Fig. 2 c). The transcriptomic profiles confirmed the distribution at the single-cell level of UMIs corresponding to well-established immune lineage marker genes of myeloid cells (*CD14*, *CD68*, *MNDA*), plasma cells (*JCHAIN*, *XBP1*, *MZB1*), B cells (*CD79A*, *CD79B*, *MS4A1*, *BANK1*), T cells (*TRAC*, *TRBC1*, *CD3D*), innate lymphoid cells (*IL7R*, *KLRB1*, *IL1R1*, *KIT*), mast cells (*CPA3*, *CD63*, *TPSAB1*), as well as markers of epithelial cells (*EPCAM*, *KRT8*, *CD9*, *FABP1*), endothelial cells (*VWF*, *PECAM1*, *CD36*, *CD151*), pericytes (*ACTA2, COL3A1, ADAMDEC1*) and fibroblasts (*COL1A2, ACTR2, C1S*). We observed higher proportions of myeloid cells, plasma cells, mast cells, T cells and stromal cells (fibroblasts, endothelial cells and pericytes) along with lower proportion of epithelial cells and ILCs in UC samples compared to NV samples. Compartment-wise overview revealed enrichment of epithelial cells, myeloid cells, B cells and T cells in IIC samples compared to LP enriched populations of stromal cells, plasma cells and mast cells in LP samples (Fig. 2d, Supplementary Table. 3b-d). On enumerating the cell type frequencies in IIC and LP compartments stratified by disease severity (Fig. 2e, f), we observed a significant increase in the proportion of myeloid cells and plasma cells with a concomitant decrease in the epithelial cells and ILCs when comparing patients (Wilcoxon signed-rank test p<0.05, Fig.2e) in IIC. Expectedly, within the stromal cells (endothelial cells, fibroblasts and pericytes) comprised a larger fraction in LP compared to IIC and LP-fibroblasts were significantly increased in severe UC compared to normal tissue (Wilcoxon signed-rank test p <0.05, Fig. 2f, Supplementary Fig. 1a).

### Active UC is associated with depletion of homeostatic γδ^+^ T cells and influx of proinflammatory T_H17_ cells into the EC

We detailed T cell-associated changes by selecting T cells *in silico* and re-clustered them using the nearest neighborhood unsupervised clustering algorithm (methods). Ten T cell sub-clusters with variable cell numbers (T0-T9, 749-6091 cells) were detected (Fig. 3a). The scRNA-seq dataset showed an enrichment of T cell specific genes in T cell subclusters such as T0, T4, T6, T7 and T8 (CD4, CD3D, *TRAC, TRBC1*), T1 and T3 (*CD8A, CD8B*, *CD3D, TRAC, TRBC2*); T2 and T9 (*CD8A, CD8B, TRDC, TRGC2*) and T5 (*TRAC, TRBC1*) (Fig. 3b). IELs (IIC related T cells hitherto referred to as intraepithelial lymphocytes/IELs) T1 (p=0.0056) and T2 (p=0.0056) and LP associated T0 (p=0.032) were significantly decreased whereas T4 and T8 were significantly increased in severe UC samples in both IIC (T4,p=0.0028; T8, p=0.0028) and LP (T4, p=0.016; T8,p=0.016; p-value from Wilcoxon signed-rank test < 0.05, Fig. 3b,c, Extended Data Fig. 3a).

**Fig. 3:**
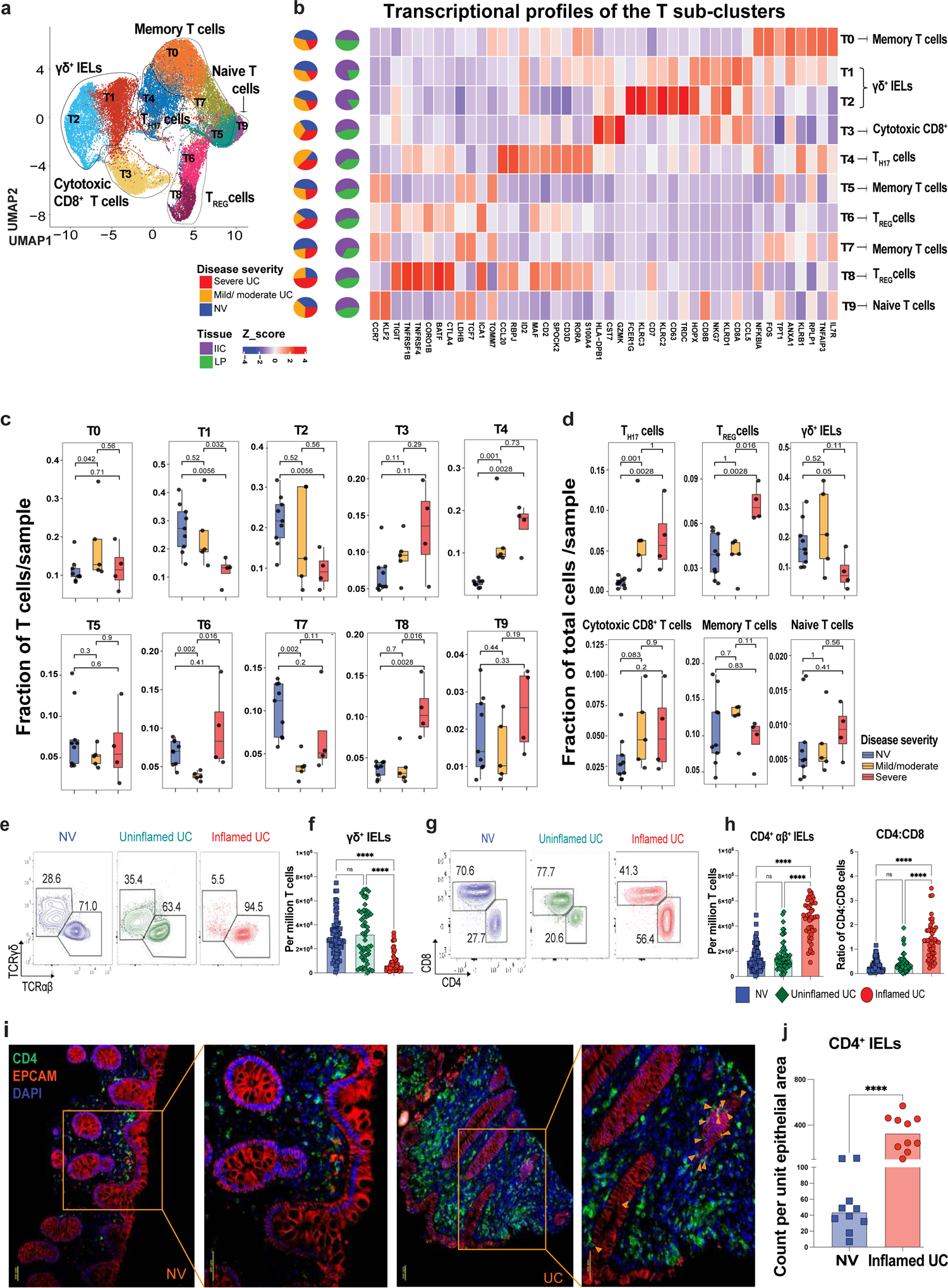
Active UC is associated with depletion of homeostatic γδ^+^ IELs and influx of proinflammatory T_H17_ cells into the EC. **a,** UMAP of T cells from NV (n=9) and active UC (n=9) colonic biopsies colored by 10 sub-clusters identified via unsupervised clustering. **b,** Heatmap showing average z scores of log normalized expression of the T cell sub-clusters defining key genes. The sub-cluster composition in terms of intestinal compartments (IIC and LP), and disease severity (NV, mild/moderate UC and severe UC) is depicted in the pie charts on the left side. **c,** Boxplots showing the distribution of per-sample relative fractions of T cell sub-clusters in the IIC stratified by disease severity. Fractions are computed with respect to all T cells in each sample. P-values from Wilcoxon signed-rank test are reported. **d,** Boxplots of per-sample fractions of T_H17_, T_REG_, γδ^+^ IELs, cytotoxic CD8^+^ T, memory and naïve T cells stratified by disease severity in IIC. Fractions are computed with respect to the total number all cells in each sample. P-values from Wilcoxon signed-rank test are reported. **e-h,** Representative flow cytometry plots to identify γδ^+^ (e) and CD4^+^αβ^+^ IELs (g) in NV, uninflamed UC and inflamed UC samples. Bar plots depicting the frequencies of γδ^+^ IELs (f), the frequencies of CD4^+^ αβ^+^ IELs (h, *left*) and CD4:CD8 ratio (h, *right*) in the inflamed samples compared to the uninflamed UC and NV samples. Cells were gated on live singlet CD45^+^CD3^+^ and on live singlet CD45^+^CD3^+^TCRαβ^+^ respectively. P-values from one-way ANOVA test are reported. ****p<0.0001. **i,** Immunofluorescence (IF) staining for CD4^+^ T cells (green), EPCAM (red) and DAPI (blue) in formalin-fixed paraffin embedded colonic sections derived from NV (*left, 10X and 20X magnification*) and patients with UC (*right, 10X and 20X magnification*). **j,** IF-based quantification of CD4+ T cells within the epithelial compartment with p-values computed using Mann Whitney test. ****p<0.0001.

Based on canonical T cell markers, the sub-clusters were regrouped into cell types and annotated as T_H17_ (T4), cytotoxic CD8^+^ (T3), naïve (T9), Regulatory (T_REG_, T6, T8), γδ^+^ (T1, T2) and memory (T0, T5, T7) T cells (Fig. 3d, Supplementary Table 4). Both IIC- and LP-associated T_H17_ cells were significantly increased in severe UC compared to NV (IIC, p=0.0028; LP, p=0.032) whereas T_REG_ cells were significantly increased only in IIC (p=0.0028) and not LP (p=0.29). In contrast, γδ^+^ IELs and LP specific memory T cells were significantly reduced in severe UC compared to NV (IIC, p=0.05; LP, p=0.016; Fig. 3d, Extended Data Fig. 3b, Supplementary Fig. 1b, c). Multiparameter flow cytometry (Supplementary Fig. 2a) confirmed a significant decrease of the frequency of EC-associated γδ^+^ IELs (p<0.0001; Fig. 3e, f). Among αβ^+^ IELs, a significant increase in CD4^+^αβ^+^ IELs (p<0.0001) and CD4:CD8 ratio (p<0.0001) was noted in the EC of patients with active UC (Fig. 3g, h, Extended Data Fig. 3c). Immunofluorescence (IF, Fig. 3 i,j) and immunohistochemical staining (IHC, Extended Data Fig. 3d,e) further confirmed an increase of CD4^+^ IELs (p<0.0001) in active UC patient colonic tissues. Gene Set Enrichment Analysis (GSEA) on gene level fold changes identified several UC-enriched pathways notable for T cell activation (*HLA-DRB1, CD3D*), cytolytic activity (*GZMB*, *PRF1*), MHC class-I antigen presentation (*TAP1*) and pro-inflammatory cytokines (*STAT1, IL2RG*, *JAK3, JAK1, STAT3, PRF1, IL2RG, JAK1*) (Extended Data Fig. 3f) suggesting an elevated secretion and activation status of IELs in the inflamed UC compared to NV samples. Altogether, a substantial remodeling of IEL was noted with a reduction in homeostatic γδ^+^ IELs and a concomitant increase in T_H17_ and T_REG_ subsets in EC.

To understand the pathogenic significance of EC-associated T_H17_ cells, we cultured normal colonic biopsy tissue with T_H17_ cell-associated cytokines^32^, using either IL-17 alone, IL-22 alone or IL-17 in combination with IL-22 (Extended Data Fig. 4a). Through subsequent RNA sequencing, we identified 119 DEGs (FCH>1.3, FDR<0.05, compared to unstimulated) upon IL-17+IL-22 stimulation (Extended Data Fig. 4b, hitherto referred to IL17+IL22 signature, Supplementary Table 5). Minimal DEGs were observed with either cytokine in monoculture. The IL17+IL22 signature genes mapped (using a published dataset^23^) mostly to epithelial cells (52%) with minor contributions from other cell types, including immune (1.2%) and stromal cells (1.2%) (Extended Data Fig. 4c). Up-regulated IL17+IL22 signature could be categorized into antimicrobial peptides (AMPs; *DEFB4A, LCN2*)^40, 41^, chemokines (*CXCL1, CXCL3*), intestinal transporters (*SLC30A, SLC25A37*)^42^ and immune regulators (*NFKBIZ, TNIP3*)^43, 44^ (Extended Data Fig. 4d). We next assessed the relevance of the ex-vivo derived IL17+IL22 signature to IBD-associated inflammation. In colonic biopsies from VC-1 patients, a significant positive correlation between the gene expression levels of IL17 and IL22 with GSVA scores of the IL17+IL22 signature were observed for inflamed biopsies (Extended Data Fig. 4e). GSVA-scores for the IL-17+IL-22 signature were significantly increased in inflamed colonic biopsies (p=3.81*10^-78^) compared to uninflamed biopsies (p=2.45*10^-6^) from VC-1 UC patients (Extended Data Fig. 4f), an association driven mainly by myeloid cell recruiting chemokine transcripts (*CXCL1, CXCL2, CXCL3*) and T_H17_- and T_REG_-recruiting chemokine transcripts (*CCL20, CCL28*) (Extended Data Fig. 4g). Notably, IL17+IL22 signature scored higher than IL17 (d=1.21, p< 2.2*10^-16^, paired t test) and IL22 (d=2.6, p< 2.2*10^-16^) monocultured signatures in inflamed biopsies, suggesting a synergistic effect of those two cytokines in UC pathogenesis. Furthermore, we observed consistent transcriptional changes between the ex vivo derived IL-17+IL-22 signature and inflammation-associated EC genes shared between the Epi_inf_ and IIC_inf_ signatures (Fig. 1g, Extended Data Fig. 4h) with 19 genes overlapping with the IL-17+IL-22 up-(OR=18.3, p=6.3*10^-16^, FCH>2, Fisher’s exact test) and 6 down-regulated genes (OR=20.5, p=0.008, FCH>2, Fisher’s exact test). Altogether, these data point to a T_H17_ cell-driven inflammatory reprogramming of the EC and upregulation of myeloid/immune cell-recruiting chemokines.

**Fig. 4:**
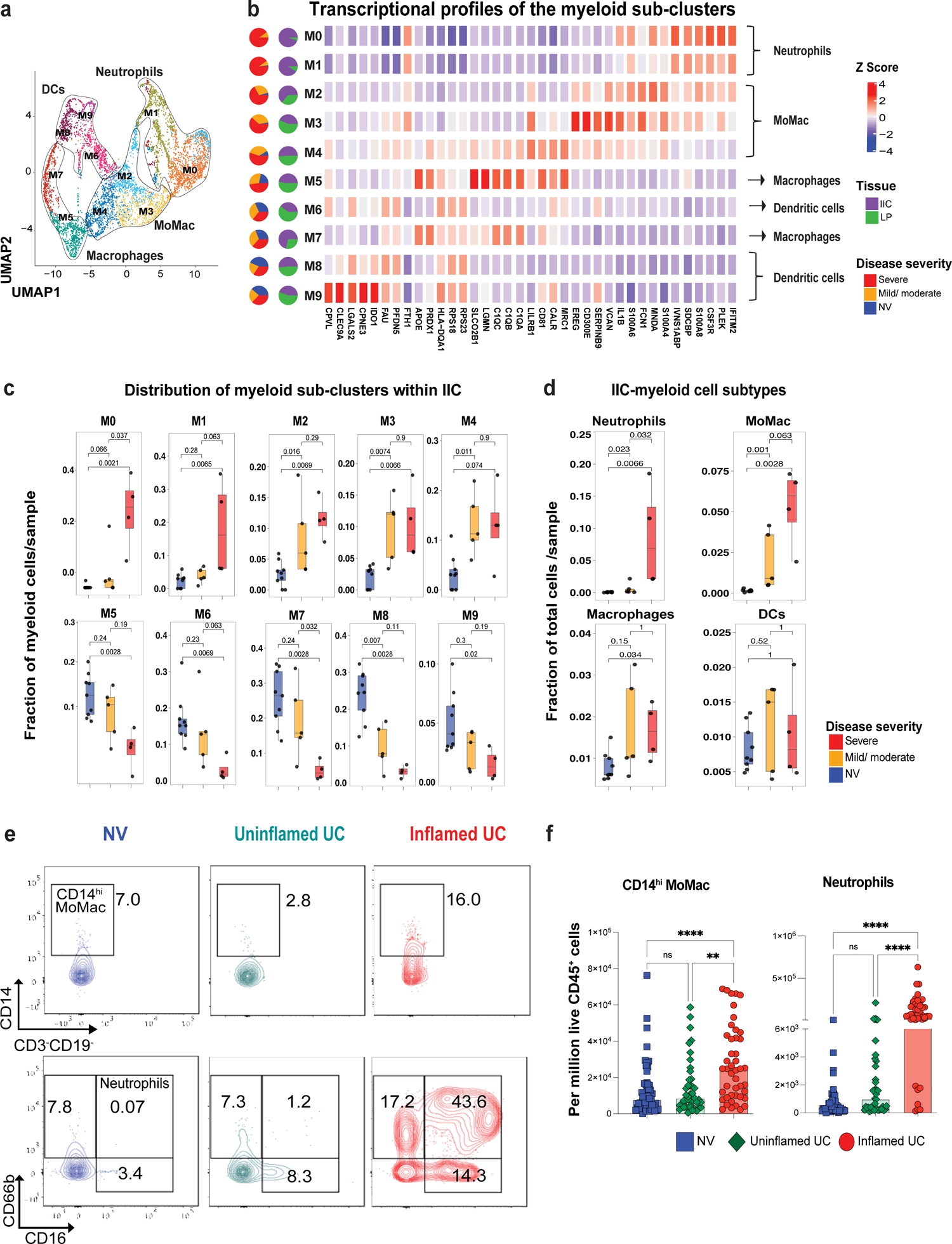
Active UC is associated with a major influx of pro-inflammatory myeloid cells into the EC. **a**, UMAP of myeloid cells from NV (n=9) and active UC (n=9) colonic biopsies colored by 10 sub-clusters (M0-M9). **b,** Heatmap showing the average z-scores of log transformed expression for myeloid sub-clusters defining genes. The sub-cluster composition in terms of intestinal compartments (IIC and LP), and disease severity (NV, mild/moderate UC and severe UC) is depicted in the pie charts on the left side. **c,** Boxplots showing the distribution of per-sample relative fractions of myeloid sub-clusters (M0-M9) in IIC stratified by disease severity. Fractions are computed with respect to all myeloid cells in each sample. P-values from Wilcoxon signed-rank test are reported. **d,** Boxplots showing the distribution of per-sample fractions of myeloid cell subtypes (i.e., neutrophils, MoMac, macrophages and DC) in the IIC stratified by disease severity. Fractions are computed with respect to the total number of all cells within each sample. P-values from Wilcoxon signed-rank test are reported. **e,** Representative flow cytometry plots identifying CD14^hi^ MoMac, CD16^hi^ MoMac, eosinophils and neutrophils in NV, uninflamed UC and inflamed UC samples. **f,** Bar plots show an increased frequency of CD14^hi^ MoMac (*left*) and neutrophils (*right*) in the epithelial compartment from inflamed UC samples compared to uninflamed UC and NV samples.

### Active UC is associated with a major influx of pro-inflammatory myeloid cells into the EC

Having observed the potential for T_H17_ cell-derived cytokines to recruit myeloid cells, we further elaborated on the myeloid cells in the scRNA-seq data and re-clustered them using the nearest neighborhood unsupervised clustering algorithm (Fig. 4a, methods) to derive 10 sub-clusters with variable cell numbers (M0-M9, 142-1099 cells, Supplementary Table 6). The transcriptomic profiles of the myeloid sub-clusters were distinct, and their distribution varied within the two intestinal compartments-IIC and LP. IIC-associated M0, M1, M2 and M3 were significantly higher in severe UC compared to NV (p = 0.0021; p= 0.0065; p= 0.0069; p= 0.0066); while M5, M6, M7 and M8 were significantly lower in severe UC compared to NV (p = 0.0028; p=0.0069; p= 0.0028; p=0.0028). The same differences between severe UC and NV were noted in the LP except for sub-clusters M1, M5 and M7 (Fig. 4b,c; Extended Data Fig. 5a, Wilcoxon signed-rank test p< 0.05). Based on canonical markers, myeloid sub-clusters were coalesced and annotated into 4 cell types: neutrophils (*CSF3R*), monocytes-macrophages (MoMac) (*EREG, MNDA*), macrophages (*MRC1, C1QA/B/C*) and dendritic cells (DCs, *LGALS2*) (Supplementary Table 6). Neutrophils and MoMac were significantly increased in the severe UC compared to NV both in IIC (p=0.0066; p= 0.0028) and LP (p=0.018; p=0.016) while macrophages were significantly increased only in the IIC (p=0.034) in the severe UC compared to NV (Fig. 4d, Extended Data Fig. 5b, Supplementary Fig. 1d,e). Using multi-parameter flow cytometry (Supplementary Fig. 2b), we observed a significant increase in the frequency of neutrophils (CD45^+^CD3^-^CD19^-^ CD16^hi^CD66b^+^cells; p<0.0001) and CD14^hi^ MoMac (live CD45^+^CD3^-^CD19^-^CD14^hi^; p<0.0001) in the EC of inflamed UC compared to the uninflamed UC and NV (Fig. 4e,f, Supplementary Table 6). Though neutrophils showed an analogous increase in the inflamed LP, their increase was more pronounced within the EC (Extended Data Fig. 5c).

**Fig. 5:**
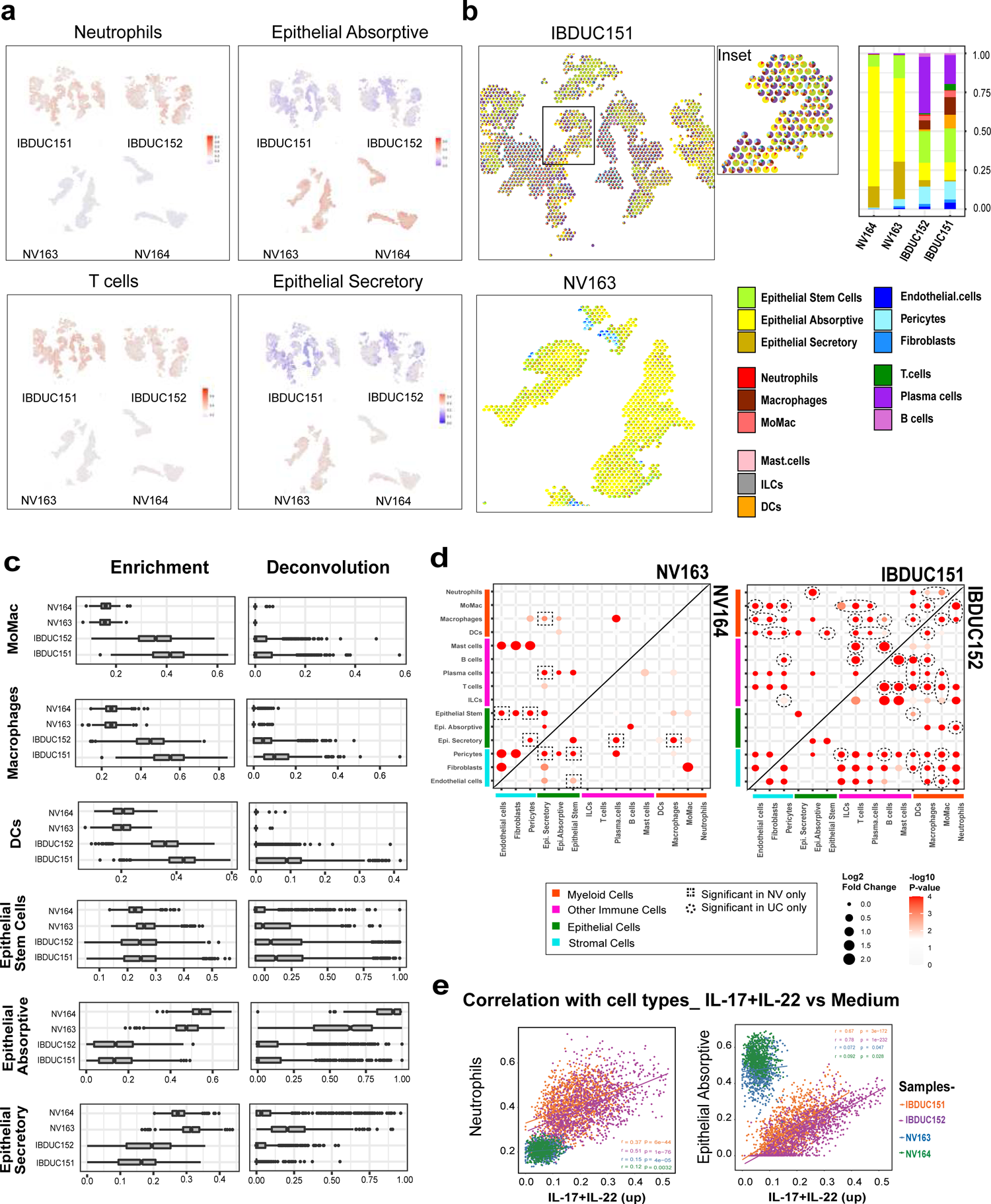
Spatial transcriptomics (ST) reveals depletion of epithelial subtypes and enhanced cell-cell interactions within colonic tissues of patients with UC. **a**, Spatial distribution of enrichment scores of neutrophils, T cells, epithelial absorptive and epithelial secretory in the four ST samples. **b,** Cell type fractions obtained via deconvolution. Spatial distribution of cell types, represented as cell type fraction pie charts at each spatial location in representative inflamed and healthy colonic biopsies. The spot diameter is 55μm and center-to-center distance between the spots is 100μm. Bar plot (*right*) showing the average fraction across all spatial spots inferred via deconvolution analysis for each sample. **c,** Per-sample distribution across spatial spots of enrichment scores and cell-type fractions inferred via deconvolution analysis for different cell types. **d,** Cell-cell co-localization analysis for NV samples (left panel) and UC samples (right). For significantly enriched co-localization pairs, the relative effect magnitude is represented by the bubble size (log2 fold change) while the significance by the bubble color (-log10 p-value). Co-localized cell type pairs detected in both NV samples and not in UC samples are highlighted with a square, while those detected in both UC samples only by a circle. **e,** Scatterplots showing enrichment scores from ST data of neutrophils and epithelial absorptive (y-axis) versus IL-17+ IL-22 gene signatures (x-axis) across spatial spots. For each scatterplot, Pearson’s correlation and corresponding correlation test p-values are shown. These statistics were separately derived for each sample.

To gain insights on the biological functions of EC-specific myeloid cell DEGs in UC (Extended Data Fig. 5d), we performed pathway enrichment analysis (GSEA) to find biological pathways dysregulated in UC samples compared to NV within the myeloid compartment (methods). This pathway analysis revealed an elevated granulocyte adhesion and diapedesis (*CXCL8, ICAM1, SELL, ITGAM*), monocyte-associated surface molecules (*CD44, ICAM1, SELL, ITGAM*), endogenous TLR signaling (*S100A8, S100A9, TLR2, CD14, VCAN*), interferon alpha/beta (*IFITM2, IFITM3, IRF1*) and interferon gamma signaling (*STAT1, SOCS3, IFNGR2, PTPN1*), TRAF3-dependent IRF activation (*TRIM25, IRF7, EP300*) and NF-κB signaling (*NFKBIA, TNFAIP3,TNFRSF1B*) in UC. In contrast, pathways up-regulated in NV included oxidative phosphorylation (*COX7C, COX41, UQCRB*), electron transport chain (*SLC25A6, SLC25A5*) and polyamine metabolism (*ADI1, SMS, SRM*) (Extended Data Fig. 5d). Altogether, these data demonstrate a major alteration in the composition of myeloid cells, associated with the emergence of neutrophils and MoMac within the epithelium leading to elaboration of potent pro-inflammatory programs along with a concomitant decrease of the homeostatic metabolic processes during active disease.

### Increase in IgG^+^ plasma cells in the EC and LP of UC

Plasma cells (PC) and B cells were selected in silico and sub-clustered into 10 PC sub-clusters and 6 B cell sub-clusters (Extended Data Fig. 6a, Supplementary Table 7, methods). P0, P1, P2, P5, P6, P7 were annotated as IgA^+^ PCs (*IGHA1, IGHA2*) while P3, P4, P8 and P9 were annotated as IgG^+^ PC (*IGHG1, IGHG2, IGHG3, IGHG4*) (Extended Data Fig. 6b). EC associated P6 and LP associated P0 were decreased whereas EC associated P3 and LP associated P3, P4, P8 and P9 were significantly increased in severe UC samples compared to NV (Extended Data Fig. 6b,c, Supplementary Fig. 3a). Consistent with our recent data^28^ IgG^+^ PC were significantly increased in both IIC (p=0.006) and LP (p=0.032) and IgA^+^ PC were significantly decreased in LP (p=0.016) of UC vs NV (Extended Data Fig. 6e, Supplementary Fig. 1f, 3c). Multiparameter flow cytometry (Extended Data Fig. 6g, h, Supplementary Fig. 2c) further confirmed a significant increase in IgG^+^ PC (p=0.0016) within the EC of UC vs NV (Extended Data Fig. 6i-k). Finally, the pathways enrichment analysis within the plasma cell compartment revealed the upregulation of class I antigen presentation (*HLA-C, HLA-A, HLA-B*), noncanonical NFKB pathway (*PSME2, UBC*) and interferon alpha pathways (*IFNAR2, IFNAR1, STAT1*) (Supplementary Fig. 3e) in UC compared to NV; highlighting elevated ER stress and interferon (IFN) imprinting on plasma cells in patients with UC samples.

**Fig. 6:**
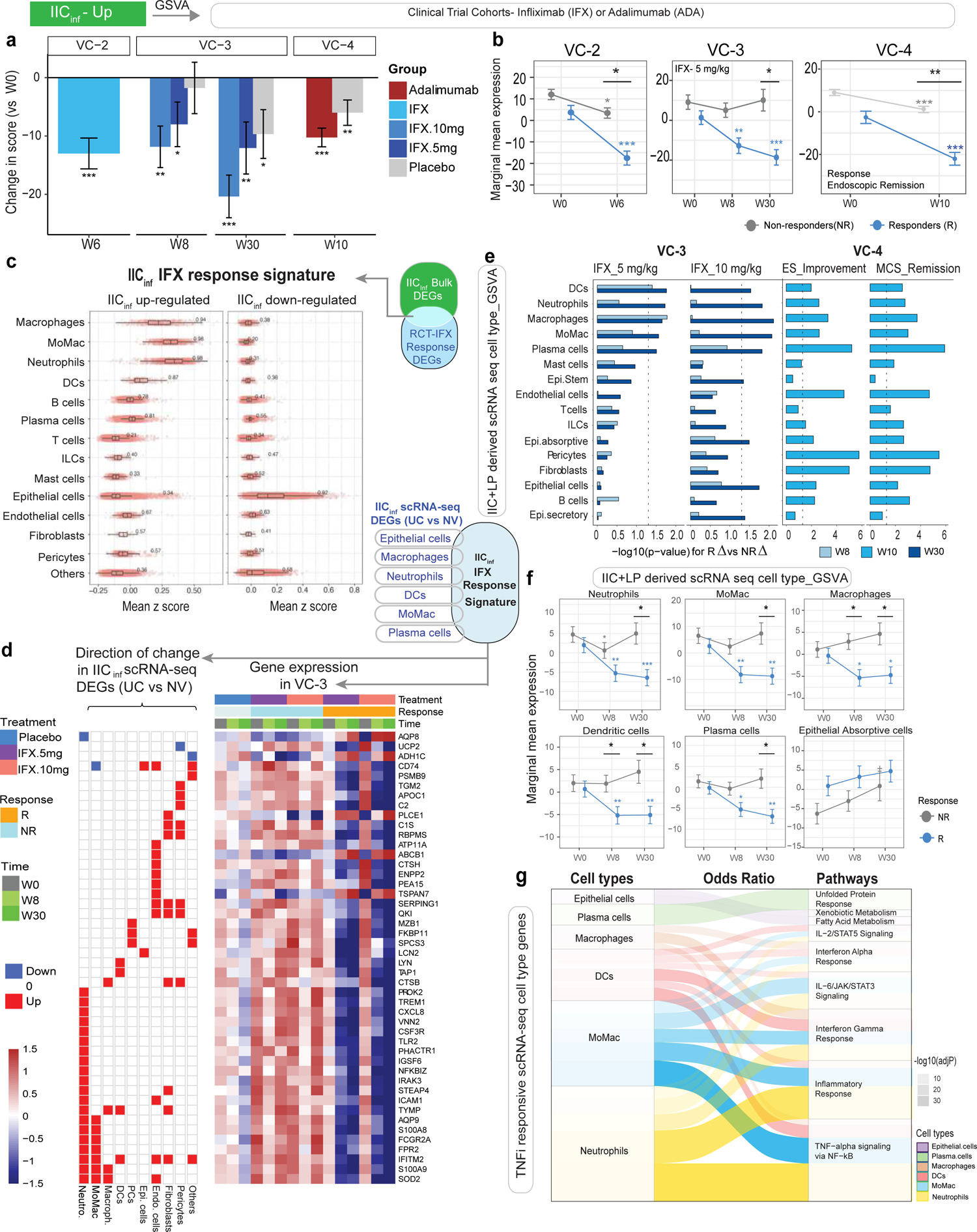
Persistence of myeloid cell and plasma cell signatures in the EC associates with anti-TNF treatment non-response in three distinct validation cohorts. **a**, Changes in score (mean ± SEM) for the activity (GSVA scores) of the IIC_inf_ signatures at baseline and after 6 weeks (in VC-2, left) or 8 and 30 weeks (in VC-3, middle) of anti-TNF therapy (Infliximab (IFX) in VC-2 and at week 10 (in validation cohort 4, VC-4, right) after adalimumab (ADA) therapy. **b,** IFX-induced changes in IIC_inf_ signatures responders (R) and non-responders (NR), where treatment response was defined by mucosal healing in VC-2, clinical response in VC-3 and by endoscopic remission in VC-4. Changes with treatment were modeled using LMEM and comparisons tested using contrasts (emmeans package in R). p values above the error bars denote significant change from baseline within the group at each time point, while p-values at the top indicate that treatment changes over time are significantly different between R vs NR, * p< .05; ** p< .01; *** p< .001 **c,** Boxplots showing the average z-score of IIC_inf_ up- and down-regulated response signatures (methods) based on the transcriptomic profile of each cell from scRNA-seq derived cell types. For each cell type, we report one-sided AUC reflecting the specific z-score to discriminate between that cell type and the rest of the cell types. **d,** Heatmap representing the expression profile of the IIC_inf_ IFX-response signature that overlaps with scRNA-seq derived cell type signatures in VC-3. On the left side of the heatmap [IIC_inf_ (UC vs NV) scRNA-seq], the status of the gene in the differential expression analysis comparing UC vs NV profiles for each cell type is indicated (up-regulated (red) or down-regulated (blue). **e,** Differential anti-TNF effect between R and NR for the activity (GSVA scores) of cell-type scRNA-seq derived signatures in VC-3 and VC-4 cohorts. Bar plots indicate -log10(p-values) for the comparison of the changes (from W0) in scores between R and NR at respective time points (W8, W10, W30). The dashed line indicates p-value=0.05, indicating a significant change in R vs NR at W30 with respect to W0. **f,** Estimated marginal mean for the GSVA scores of scRNA-seq derived cell types that were significantly different between R and NR at week 30 from the start of IFX therapy at dose 5 mg/kg. **g,** Alluvial plots showing the pathway ORA for the leading-edge genes (LEGs) of each cell type (only those associated with IFX response in f are shown) using mSigDB Hallmark^80^ database. Color density denotes adjusted p-values while the band width denotes odds ratio of the pathway enrichment.

Among B cells, we annotated germinal center (GC) B cells (B5, *AICDA*, *MME*, *CD38* and *CD27*), naïve B cells (B2 and B3; *IGHD, FCER2*) and memory B cells (B0 and B1; *TNFRSF13B*/*TACI*) (Extended Data Fig. 6a,b,d,f; Supplementary Fig. 1g, 3b,d) with comparable distribution between UC and NV. Consistent with the scRNA-seq data, we did not detect significant changes in total, naïve and switched memory B cell frequency between UC and NV (Extended Data Fig. 6l-p, Supplementary Fig. 2d) by flow cytometry, although and in consistence with our recent report^28^, EC-associated B cells were enriched in IFN alpha signaling pathway (*STAT1, IRF9*) (Supplementary Fig. 3f). Enhanced type-1 interferon signaling in B cells and plasma cells is not only reported to be associated with autoimmune conditions but also has detrimental proinflammatory effects on non-autoimmune diseases^45,46^. Altogether, these data show a remodeling of EC-associated PC, analogous to LP-derived PC, with an increase in IgG^+^ PC in UC.

### Spatial transcriptomics reveals depletion of epithelial subtypes and enhanced cell-cell interactions within colonic tissues of patients with UC

To study the inflammation in a spatially coherent context, we used spatial transcriptomics (ST) to map the regional distribution of cell types in UC (n=2) and NV (n=2) (Supplementary Fig. 4a). The 4 samples contained between 11660 and 15846 genes, with the mean of 4438 unique genes per spot in the two UC samples, and 2873 in NV. We performed GSEA^47^ to characterize the relative spatial distributions of different cell-types using cell-type specific signatures from scRNA-seq data, together with three epithelial (*LGR5^+^* epithelial stem, *BEST4^+^ OTOP2^+^* absorptive and *BEST2^+^ WFDC2^+^* secretory cells) signatures from the literature^23^. Enrichment scores indicated an infiltration of the immune cells such as neutrophils and T cells in active UC samples and a depletion of epithelial cells (Fig. 5a, Supplementary Fig. 4a). To estimate cell type fractions across spatial spots, we performed the deconvolution of ST data via spatialDWLS^48^ (Fig. 5b) leveraging our scRNA-seq data as reference (Methods). Consistent with the enrichment analysis, deconvolution analysis revealed that a significant fraction of UC samples was comprised of different types of immune cells whereas NV mostly consisted of epithelial cells (Fig. 5b). Both enrichment analyses and cell-type deconvolution revealed increased proportions of immune cells including the four myeloid cell types, T cells, B cells, plasma cells, epithelial stem cells and the three stromal cell types in UC with a concomitant decrease in mature epithelial absorptive and secretory cell types (Wilcoxon signed-rank test, all p < 10^-16^). While epithelial stem cells showed a prominent presence in both UC and NV, these cells comprise a significantly higher portion of epithelium in UC (64% and 57% in UC, 15% and 8% in NV, Fig. 5c, Supplementary Fig. 4b-c, 5a-b, Supplementary Table 8).

To investigate cell-cell interactions, we performed co-localization analysis of different cell types in ST data for each patient (methods) and observed co-localization between myeloid cells and T cells as well as co-localization between neutrophils and epithelial absorptive cells solely in UC (Fig. 5d, Supplementary Table 8). Given the significant proximity of neutrophils and epithelial absorptive cells in UC samples (Fig. 5d), we investigated if neutrophils and epithelial cells were associated with up-regulated gene signature from IL-17+IL-22 treated biopsies. Our association analysis based on ST data confirmed higher positive correlations with the activity of the IL-17+IL-22 signature in UC (neutrophils: r=0.37, r=0.51, p<10^-43^; epithelial absorptive cells: r=0.67, r=0.78, p<10^-170^) compared to NV (neutrophils: r=0.15, r=0.12; epithelial absorptive cells: r=0.07, r=0.09, p<10^-2^, Fig. 5e). These results further confirm that myeloid cells could be stimulated by T_H17_ associated signals.

Quantitative IHC revealed a significant increase in CD14^+^ MoMac in UC vs NV, both infiltrating the colonic crypts (p=0.02) and in a peri-crypt distribution (p=0.0001; Extended Data Fig. 7a, b, Supplementary Fig. 6). Further, IF staining showed a significant increase in CD68^+^ macrophages in UC vs NV (p=0.01; Extended Data Fig. 7c,d, Supplementary Fig. 7). Finally, a significant increase in EC-associated MPO^+^ neutrophils was noted in inflamed UC vs NV (p=0.008; Extended Data Fig. 7e,f, Supplementary Fig. 8). The microscopy findings thus affirmed the epithelial cell-myeloid cell proximity observed using ST. Overall, consistent with our sequencing data, ST-derived data reveal a major loss of mature enterocytes and increased T cell-myeloid cell-epithelial cell proximity in UC, with T_H17_-cell driven imprinting of UC tissues.

### Persistence of myeloid cell and plasma cell signatures in the EC associates with anti-TNF treatment non-response in three distinct validation cohorts

We first tested the clinical relevance of DEGs associated with inflammation in the EC (IIC_inf_) (Fig. 1c) using three published data sets of UC patients treated with TNFi (Extended Data Fig.1, cohort details in methods). The activity of up-regulated IIC_inf_ genes was significantly reduced for both, 5 and 10 mg/kg doses of Infliximab/IFX- at week 8 (Δ_IFX-10_= −11.8(−25.4%), Δ_IFX-5_= −8 (−19.9%)) and week 30 (Δ_IFX-10_= −20.4 (−43.7%), Δ_IFX-5_= −12.1 (−30%)), in VC-3 (Fig. 6a, p<0.05) while placebo patients had no significant change over 8 weeks. Similar effects were observed in VC-2 with IFX ( Δ_IFX_= −13 (−28%), p<10^-5^) and VC-4 with Adalimumab/ ADA (p<10^-3^), Extended Data Fig. 8a)

Next, we determined if expression of IIC_inf_ up-regulated genes differed between responders (R) and non-responders (NR) to TNFi. The IIC_inf_ score after 6 weeks of IFX treatment (p<0.05) was significantly lower in patients that respond to IFX treatment (defined as mucosal healing) as compared to NR in VC-2 (R: Δ_IFX_= −21 (−51.7%), NR: Δ_IFX_= −8 (−17.5%)) (Fig. 6b). Similarly, in VC-3, TNFi R (defined as clinical response) demonstrated a significantly greater decrease (p<0.05) in the IIC_inf_ score (R Δ_IFX-5_=-20 (−54.1%), Δ_IFX-10_=-26 (−56.9%)) as compared to NR (Δ_IFX-5_=-1 (2.2%), Δ_IFX-10_=-2 (−3.6%). Finally, in VC-4, IIC_inf_ up-regulated genes were significantly reduced in endoscopic R to TNFi therapy (Δ_ADA_ R= −27.6(−45.3%), p=2.25*10^-13^; NR=-5.14 (−7.8%), p=0.005) (Fig. 6b, Extended Data Fig. 8b). Overall, these observations support that inflammation associated genes expressed in the IIC compartment are potentially targeted by TNFi and are important for disease remission.

Having observed a significant decrease in IIC_inf_ scores with TNFi therapy, we sought to determine the IIC_inf_ genes that reversed with TNFi response using the VC-3 RCT dataset. From the IIC_inf_ bulk signature, we identified 255 genes (159 up-regulated and 96 down-regulated, *IIC_inf_ IFX response signature*, detailed in methods, Supplementary Fig. 9, Supplementary Table 9). Pathway enrichment analyses on the IIC_inf_ IFX-response signature (Supplementary Table 10) revealed proinflammatory pathways such as IL-17, TNF and NF-kB signaling pathways together with Th1, Th2 and Th17 differentiation pathways as IIC specific mechanisms attenuated with TNFi therapy in patients responding to the treatment. To delineate which cell types (in the epithelium compartment) express these treatment response genes we projected the IIC_inf_ IFX response signature onto our scRNA-seq data and noted higher expression of genes down-regulated with TNFi in myeloid cells including neutrophils, MoMac, macrophages and DCs, while genes up-regulated with TNFi were expressed in epithelial cells (Fig. 6c). To identify the likely TNFi responsive genes within cell types, we next intersected the IIC_inf_ IFX response signature with our scRNA-seq IIC-DEGs (UC vs NV). Genes up-regulated in active UC samples that were down-regulated in TNFi responders (VC-3 cohort) primarily included IIC-associated myeloid cells (CXCL*8, CSF3R, FPR2, FCGR2A, SOD2, TLR2, TREM1* and *STEAP4)* while the down-regulated genes in active UC samples that were up-regulated such as *ABCB1, ADH1C* in epithelial cells were up-regulated with treatment (Fig. 6d).

To further explore the importance of the IIC in TNFi response, we projected the cell type defining signature sets (Fig. 2b) onto the clinical trial associated datasets (VC-2, VC-3 and VC-4, Fig. 6e, Extended Data Fig. 9a-e, Supplementary Fig. 10-13). In this way, we explored the contribution of both IIC and LP-resident cells together in relation to TNFi response. Overall, we observed consistent decreased expression of genes enriched in myeloid cells (neutrophils, MoMac, macrophages, DCs), plasma cells and increased expression for genes marking epithelial cells (secretory, absorptive and stem cells) (Fig. 6e). Fig. 6f highlights the key cell types associated with treatment response in VC-3, where we found that both R and NR showed significant difference in genes associated with neutrophils, MoMac, macrophages, DCs, plasma cells at 5mg/kg dose of IFX. Furthermore, both VC-3 and VC-4 suggested that early W8-W10 reduction in myeloid cell- and plasma cell-associated genes was associated with treatment response. In VC-4, additional genes associated with mast cells, stromal cells, T cells, epithelial cells and B cells were also observed significantly changed at W10 compared to W0 (Fig. 6e) with TNFi. Conversely, increased expression of epithelial cell-associated genes with TNFi therapy was associated with therapeutic response (Extended Data Fig. 8e, f, Extended Data Fig. 9b, d,e, Supplementary Fig. 11,12,13).

By correlating expression of the cell type defining signature genes with anti-TNF response in VC-3 (Pearson’s correlation for genes that have correlation>0.5 and FDR<0.05), we identified leading-edge genes (LEGs) considered as ‘driving’ anti-TNF response for each of the cell types. Pathway enrichment analysis of these LEGs revealed TNF-α signaling, interferon-γ response, and IL-6-JAK-STAT signaling as well as IL-17 signaling and Th1, Th2 and Th17 differentiation and neutrophil extracellular trap formation as pathways associated with anti-TNF therapeutic response in neutrophil-, MoMac- and macrophages. LEGs in plasma cells associated with TNFi response included unfolded protein response and protein export while LEGs in epithelial cell genes were enriched in fatty acid and xenobiotic metabolism along with cell adhesion and tight junction related pathways (Fig. 6g, Supplementary Fig. 14, Supplementary Table 10). Using both bulk sequencing and scRNA-seq approaches, we narrowed down to an overlapping set of cells and pathways to be associated with therapeutic non-response. Since these pathways remain elevated in TNFi NR, these could be potentially good targets for primary NR to therapy.

## Discussion

EC-based investigations are highly relevant to understanding the pathogenesis of UC. However, EC-focused high-dimensional studies in UC are sparse^23,25^. Additionally, EC-specific immune details get overlooked with whole tissue-based, and LP-based analyses of intestinal biopsies^22,24^. Here, using two distinct cohorts of patients, and integrating bulk RNA-seq-, scRNA-seq-, spatial transcriptomic-, flow cytometry- and microscopy-derived data, we have identified novel cellular and transcriptional signatures specific to the EC in patients with UC. We show that the molecular signature associated with colonic inflammation (IIC_inf_) highly correlates with UC severity (clinical, endoscopic and histologic). By identifying high resolution intestinal inflammation signatures in our data and intersecting them with 3 published longitudinal patient cohorts, including subjects from a seminal clinical trial of TNFi in UC^49-51^, we have identified key transcriptional and cellular correlates of TNFi failure in UC. Identification of such cellular modules associated with anti-TNF non-response can be harnessed to identify early correlates of treatment non-response (based on the trajectory of response to anti-TNF in UC at early time points such as week 6 (based on VC-3) and week 10 (based on VC-4), as well as to stratify patients as per the likelihood of response. Thus, the present study details high resolution EC dynamics in UC, paves the way for development of novel biomarkers to identify TNFi failure, and provides a rational basis of the design of sequential and combinatorial therapies in UC.

In addition to nutrient absorptive and secretory functions, to facilitate mucosal homeostasis, colonic epithelial cells tether mucosal immune cells such as IELs and mediate innate and adaptive mucosal immunity^52^ through bidirectional communication with immune cells^9,11,53-55^. During the inflamed state in UC, we observed a notable loss of differentiated enterocytes, both absorptive and secretory, while the stem cell-like enterocytes were numerically preserved. Accordingly and associated with the loss of epithelial tethers, physiological EC residents, especially γδ^+^ IELs which are known to enhance barrier function^56^, dampen inflammation^57^ and regulate αβ^+^ IELs^5^^8^ were significantly depleted from the EC in UC as was previously reported in CD^26^. γδ^+^ IEL loss from the EC was accompanied by a major influx of CD4^+^αβ^+^ IELs, especially T_H17_ and T_REG_ cells encoding a number of pathogenic, pro inflammatory programs, associated with tissue damage and worsening of disease activity^57, 59^ that included TNFR2 signaling^60^, TCR activation^61, 62^, IL-2/STAT5 signaling ^63^, IL-7 signaling^64^, IL-9 signaling^65^ and IL-12 signaling ^66^.

In addition to T cell-associated changes, multiple orthogonal approaches demonstrated a dominant presence of myeloid cells (including neutrophils, inflammatory monocytes, inflammatory macrophages and DCs), infiltrating or proximate to the EC during active inflammation. Notably, we could capture previously unreported neutrophils in our scRNA seq dataset and ascribe this to the use of fresh biopsies, gentle EDTA-based dissociation protocols and short processing times in our scRNA-seq methodologies. In addition to infiltrating neutrophils, and inflammatory monocytes, homeostatic myeloid cells, especially cDCs and resident macrophages, key sentinels and drivers of tissue homeostasis at steady state^67^, switched to inflammatory phenotypes (including inflammatory macrophages) and were noted to encode powerful inflammatory programs including interferon alpha/beta signaling, interferon gamma signaling, IRF activation and NF-κB signaling in UC, likely contributing in a major way to disease pathogenesis^67, 68^. Spatial transcriptomic data and microscopic approaches corroborated a significantly increased proximity between myeloid cells and epithelial cells during UC. Furthermore, IL-17/IL-22-stimulated colonic tissues up-regulated key myeloid-recruiting chemokine genes including *CXCL1*, *CXCL2* and *CXCL3*, providing a mechanistic link between the observed T cell and myeloid cell perturbations in UC. Top drivers of histologic activity in UC were also dominated by myeloid-cell associated genes, including *S100A8/S9*, *FPR1* and *FPR2*^69^, *FCGR3B*^70^, *CLEC4A*, *TREM1*^39, 71^*, CXCR1/2*^72^ and *CXCL1*. Other notable genes associated with increased histological activity included epithelium-related *SAA1*^73^, innate immune related *LILRA5,* oxidative stress sensor /responder genes *VNN3*^74^, *HIF1A*^75^, *DRAM1*, *XBP1* and *AQP9.* These data provide potential biomarkers for monitoring of disease activity, in addition to some established biomarkers such as myeloid cell-derived calprotectin ^76, 77^.

A major aspect of our report details cell signatures in UC that are associated with TNFi-nonresponse. This was facilitated by intersecting high-resolution tissue data with three published cohorts, of which two were large, randomized placebo-controlled, clinical trials with a cumulative total of 252 patients^50, 51^. In both clinical trials, we noted unequivocal and a significant increase in the frequency of neutrophils, inflammatory MoMacs, inflammatory macrophages and PC in non-responders to TNFi. Analogously, early, week 8-10 reductions in the frequencies of epithelium-associated myeloid cells and absorptive and secretory enterocytes were associated with TNFi treatment-response. These data, first in UC, have potentially significant implications for patients who do not respond to TNFi and are reminiscent of the GIMATs-associated myeloid cell drivers of TNFi-non response in CD^22, 24^. Findings from our study indicates the prominence of epithelium associated myeloid cells to be associated with lack of early response in UC patients on TNFi^78^. In sight of multitude of treatment options in UC, our findings support the development of myeloid - targeted therapies particularly in TNFi-non responders. Among the clinically approved drugs, JANUS Kinase inhibitors (JAKi) target many of the pathways that persist in TNFi-non responders. Our work proposes a rationale to use JAK inhibitors in combination with or sequentially in the face of TNFi non-response in alignment with a recent meta-analysis report^79^. Thus, by detailing EC-compartment-associated changes in UC and elaborating on cell- and transcriptomic-signatures associated with non-response to TNFi, here we provide rational underpinnings for therapeutic decision-making with major clinical implications.

## Supporting information

Legends for Supplementary materials

## Acknowledgements

We thank the patients who participated in the study. We thank Travis Dawson, Rachel Chen and Dr. Zhihong Chen in Human Immune Monitoring Centre at the Icahn School of Medicine for their single cell transcriptomic assay technical assistance and the Mount Sinai Flow Cytometry Core for support. This work was supported by the following grants: NIH/NIDDK R01 123749 (SM). CA was supported in part by The Leona M. and Harry B. Helmsley Charitable Trust and RC2 DK122532/DK/NIDDK NIH. JCM is supported by NExT “Junior Talent”, ANR JCJC (ANR-20-CE17-0009) and ARC Programmes labellisés (PGA). RCU is supported by an NIH K23 Career Development Award (1K23DK111995-01A1).

## Author contributions

DJ, FP, MSF, CA and SM drafted the manuscript. SM designed the study, supervised experimental data collection and coordinated integration of collaboration between all participating laboratories. DJ led the work, conducted experiments, analyzed the data and coordinated collaboration between all participating laboratories. ZAT, MSF and CA performed bulk RNA seq analyses and TNFi treatment response analyses. AK and FP performed single cell RNA seq analyses. AK, FP and GCY performed spatial transcriptomic analyses. STE, JBM, JAH and AN performed analysis on VC4. JFC, RU, MK and JM contributed patient-related samples and clinical analyses. JCM, AF, TL, AB and RJ performed in vitro stimulation of biopsies with IL-17/IL-22 and the relevant analyses. All other authors contributed to experimental data and analyses, and along with SM, DJ, FP, MSF and CA, they critically reviewed and edited the final version of the manuscript.

## Extended Data Figure Legends

**Extended Data Fig. 1:**
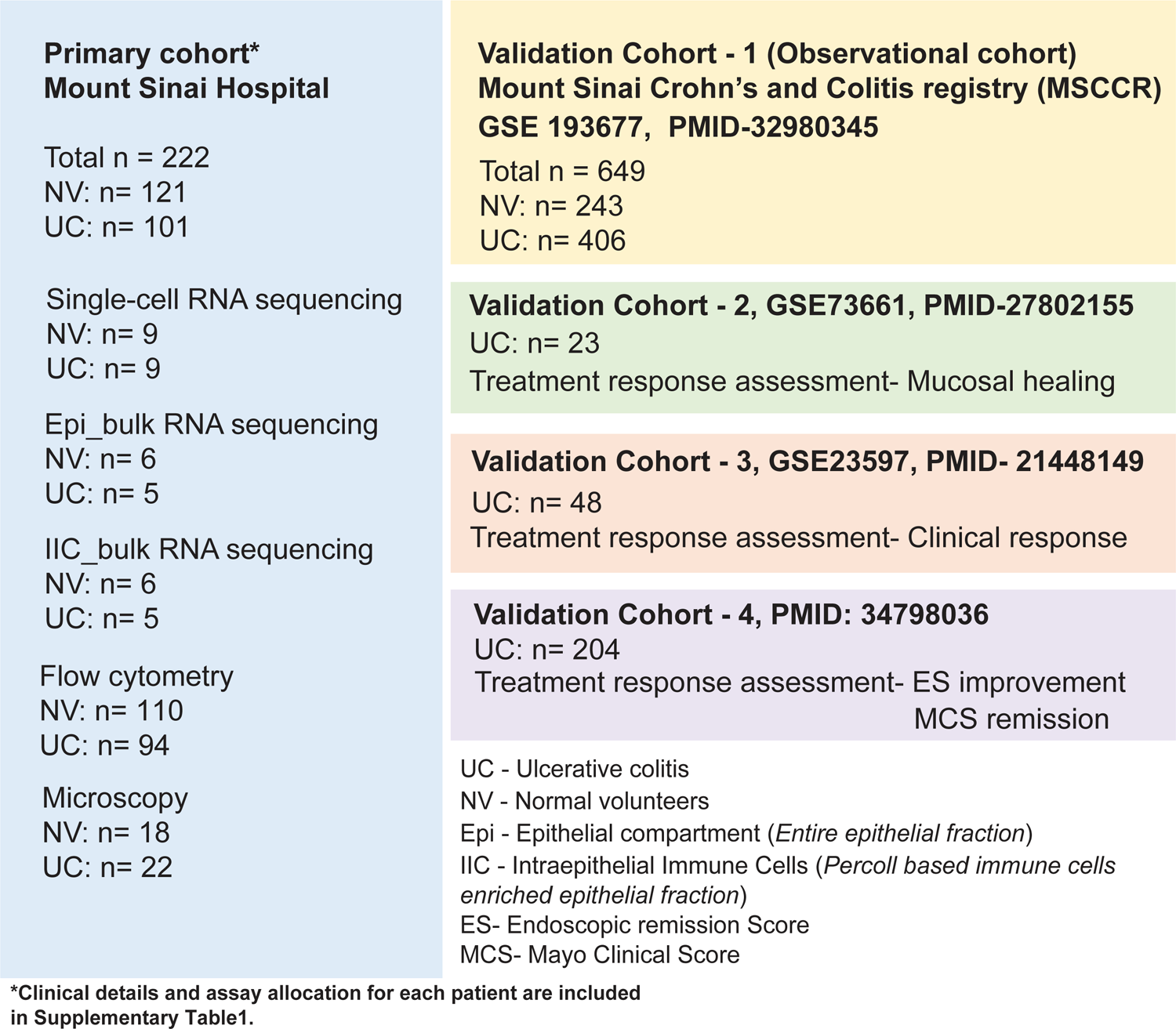
Outline of the patients and samples included in the study.

**Extended Data Fig. 2:**
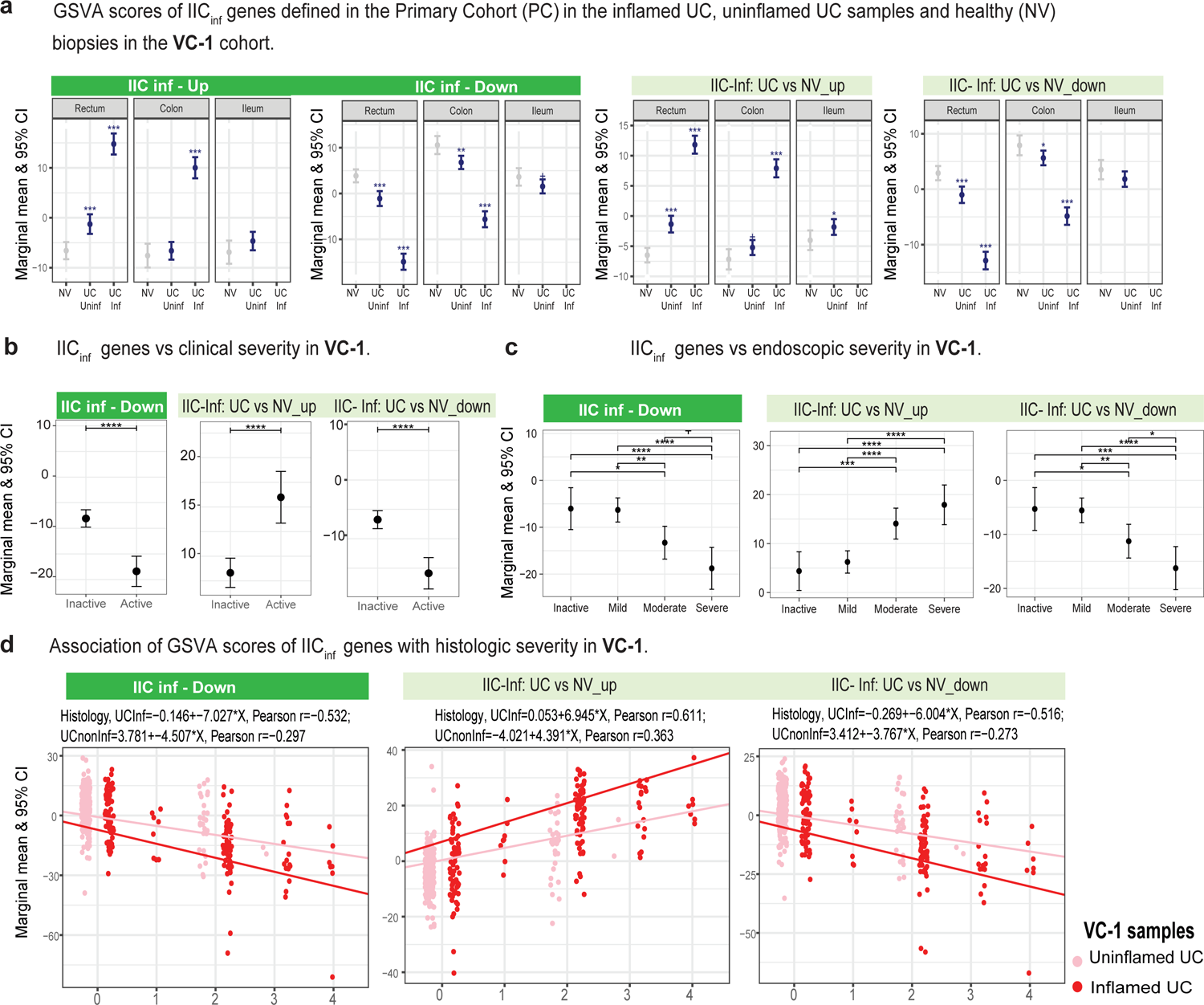
Association of IIC_inf_ GSVA activity scores in biopsies from the Validation Cohort-1/ VC-1 stratified by inflammation status and disease severity. **a,** Expression of IIC_inf_ GSVA activity scores of the inflamed UC samples from the Primary Cohort (PC) in the inflamed UC, uninflamed UC samples and NV (control) samples in the VC-1. **b,** Association of the mean marginal expression for GSVA score of IIC_inf_ genes from PC with clinical severity in VC-1. **c,** Association of mean marginal expression for GSVA score of IIC_inf_ genes with endoscopic severity in VC-1. **d,** Association of mean marginal expression for GSVA score of IIC_inf_ genes with histologic severity in VC-1. Up and down indicate direction associated with inflammation.

**Extended Data Fig. 3:**
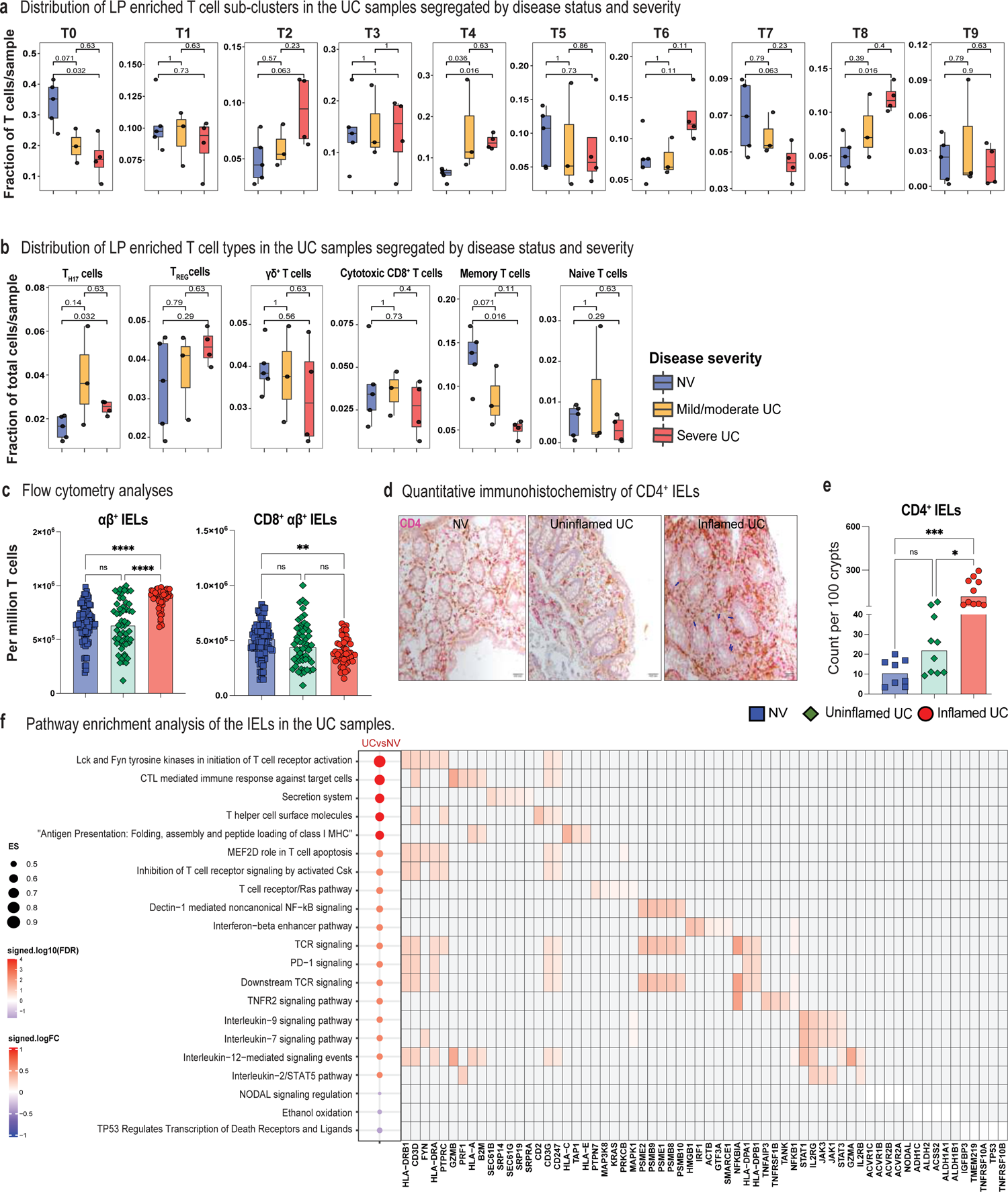
Distribution of T cell types in the LP in patients with active UC and pathway analysis of IELs. **a,** Boxplots showing the distribution of per-sample relative fractions within the T cell compartment of different sub-clusters in LP stratified by disease severity. Fractions are computed with respect to all T cells in each sample. P-values from Wilcoxon signed-rank test are reported. **b,** Boxplots of per-sample fractions of T_H17_, T_REG_, γδ^+^ IELs, cytotoxic CD8^+^ T, memory and naïve T cells stratified by disease severity in LP. Fractions are computed with respect to the total number of all cells in each sample. P-values from Wilcoxon signed-rank test are reported. **c,** Flow cytometry-based quantification of the frequencies of αβ^+^ T cells and CD8^+^ αβ^+^ T cells. **p<0.01,****p<0.0001. **d,** Immunohistochemical (IHC) staining showing the infiltration of CD4^+^ T cells (pink cells) within the EC. Representative sections from an inflamed UC biopsy (black arrows, *right panel*) uninflamed UC biopsy (*middle panel*) and NV biopsy (*left panel*) are shown. **e,** Bar plot comparing intraepithelial CD4^+^ T cell count between NV, uninflamed UC and inflamed UC samples, based on IHC staining. P-values from one-way ANOVA test are reported, ns=not significant, *p<0.05; ***p<0.001. **f,** Heat map shows pathway enrichment analysis from IELs in UC versus NV samples. The bubble plot on the left side of the figure shows the summary statistics from pathway analysis; with the size of the bubble corresponding to the enrichment score (ES) while the bubble color to the signed -log10 adjusted p-value. The heatmap at the right side shows the log fold change between UC and NV for the top five leading edge genes in each pathway.

**Extended Data Fig. 4:**
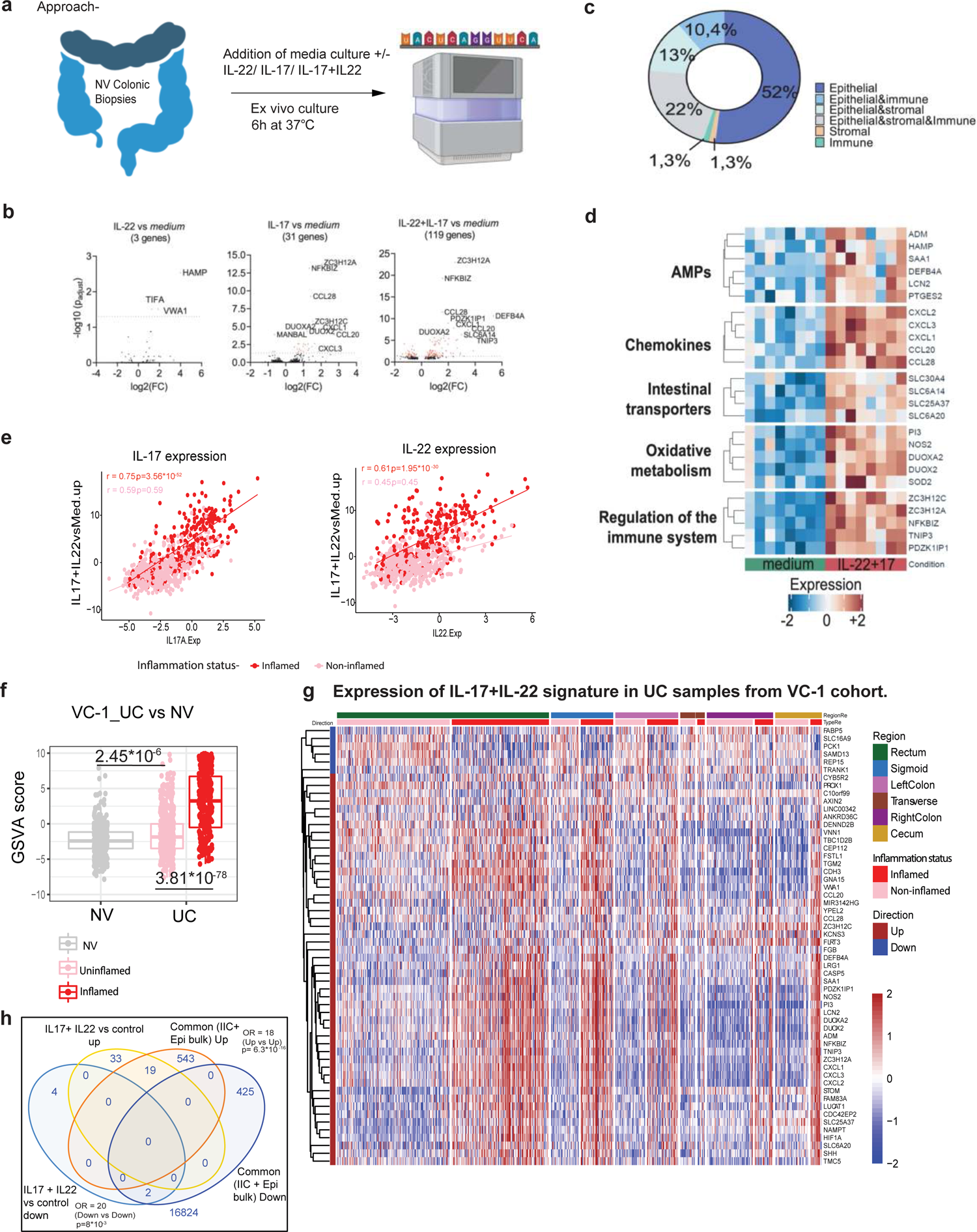
IL-17 and IL-22 stimulation led to proinflammatory transcriptional reprogramming of healthy colonic tissue. **a,** Methodological approach adopted to examine the effect of IL-17 and IL-22 stimulation on colonic intestinal biopsies. **b,** Volcano plots depicting top differentially expressed genes post-IL-22, IL-17 and IL17+ IL-22 stimulation of colonic biopsies derived from RNA sequencing. **c,** Cell types corresponding to the DEGs from RNA sequencing done on colonic biopsies stimulated by lL-17 + IL-22 cytokines. **d,** Heatmap showing biological functions for gene expression levels for up-regulated genes after colonic biopsies were stimulated with medium containing IL-17 and IL-22 compared to medium only. **e,** Correlation plots for IL17 and IL22 gene expression in VC-1 with the GSVA scores of IL17+IL22 signature (up-regulated genes) from the ex-vivo experiment using Pearson’s correlation. **f,** Box plots indicating GSVA scores of up-regulated IL-17+IL-22 signature post stimulation of colonic tissues in VC-1 cohort stratified by inflammation status (NV, uninflamed and inflamed samples from UC patients) used estimated marginal means using lsmeans function from emmeans R package. **g,** Heatmap showing the expression of the IL-17+ IL-22 signature per intestinal segment (large intestine) in VC-1. **h,** Venn diagram showing the intersection of genes that were up-/down-regulated by IL-17+IL-22 stimulation and intersecting DEGs common to Epi_inf_ and IIC_inf_ (as shown in Fig. 1d) using Fisher’s Exact Test.

**Extended Data Fig. 5:**
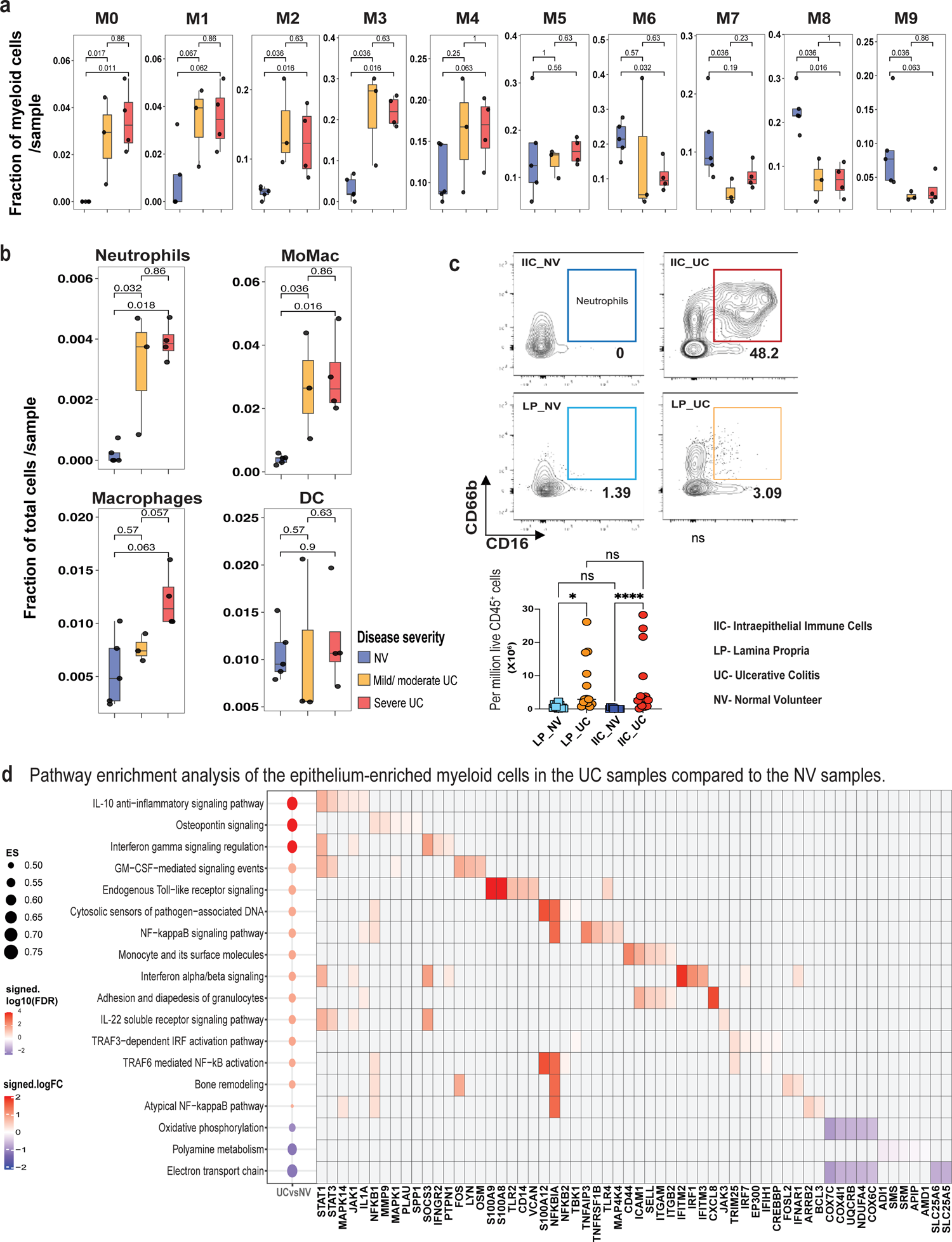
Distribution of myeloid cell types in the LP in patients with active UC and pathway analysis of IIC enriched myeloid cell types. **a,** Boxplots showing the distribution of per-sample relative fractions within the myeloid cell compartment of different sub-clusters in LP stratified by disease severity. Fractions are computed with respect to all myeloid cells in each sample. P-values from Wilcoxon signed-rank test are reported. **b,** Boxplots of per-sample fractions of myeloid cell types (i.e. Neutrophils, MoMac, Macrophages and DCs) stratified by disease severity in LP. Fractions are computed with respect to the total number of all cells in each sample. P-values from Wilcoxon signed-rank test are reported. **c,** Flow cytometry comparing of EC-associated neutrophils and LP-associated neutrophils in the UC samples. **d,** Pathway enrichment analysis of epithelium-enriched myeloid cells in UC samples compared to NV samples. The bubble plot on the left side of the figure shows the summary statistics from pathway analysis; with the size of the bubble corresponding to the enrichment score (ES) while the bubble color to the signed -log10 adjusted p-value. The heatmap at the right side shows the log fold change between UC and NV for the top five leading edge genes in each pathway.

**Extended Data Fig. 6:**
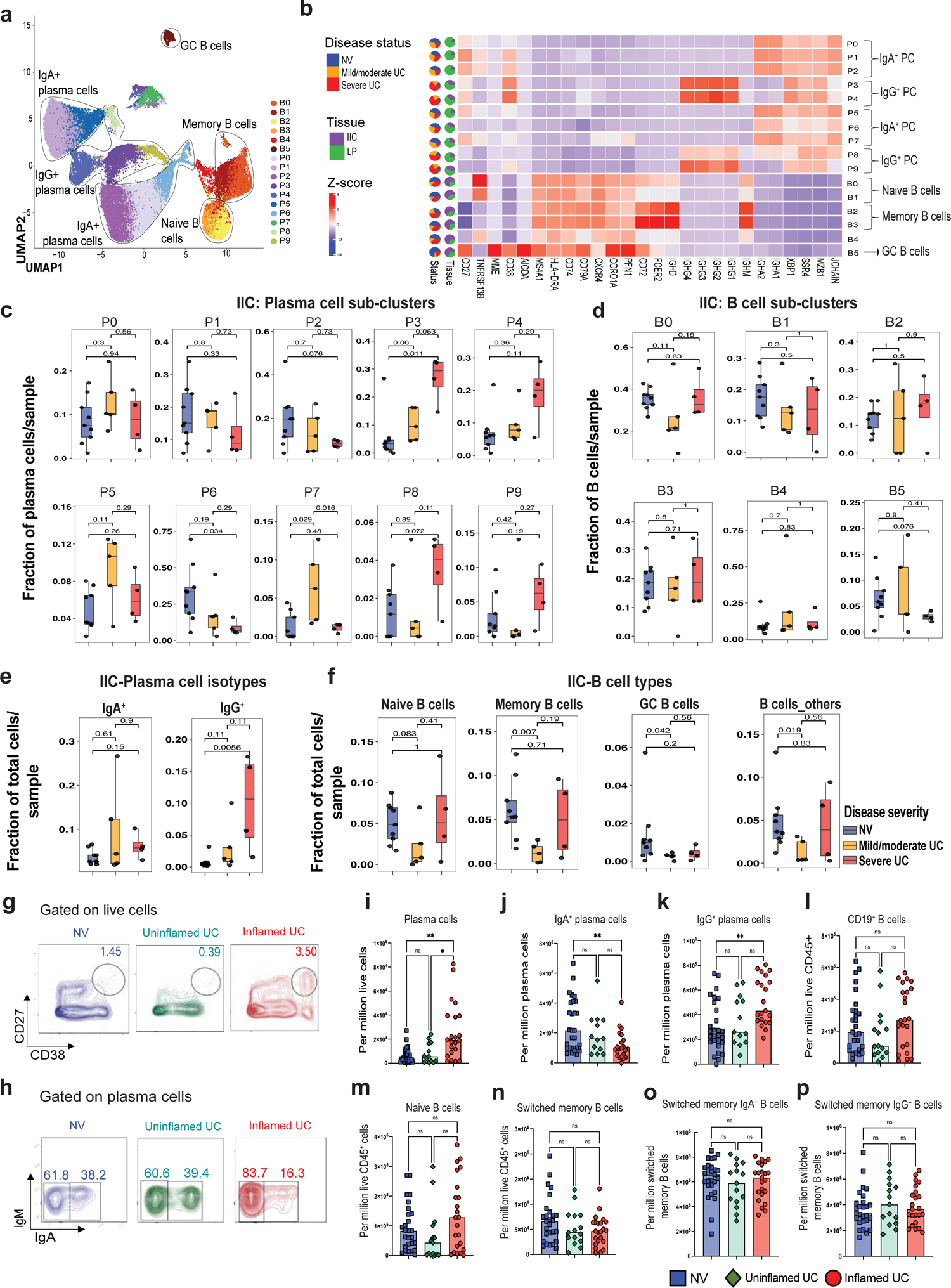
Altered transcriptomic profiles of plasma cells and B cells in active UC patients and NV. **a,** UMAP of plasma cells and B cells from NV (n=9) and active UC (n=9) colonic biopsies by unsupervised clustering to demonstrate 10 plasma cell sub-clusters and 6 B cell sub-clusters**. b,** Heatmap showing the average expression z-scores of plasma cells and B cells defining genes for each sub-cluster. The sub-cluster composition in terms of intestinal compartments (IIC and LP), and disease severity (NV, mild/moderate UC and severe UC) is depicted in the pie charts on the right side of the heatmap. **c-d,** Boxplots showing the distribution of plasma cell sub-clusters (c) and B cell subclusters (d) in the IIC stratified by disease severity. Fractions are computed with respect to all plasma (c) or B (d) cells in each sample. P-values from Wilcoxon signed-rank test are reported. **e-f,** Boxplots showing the distribution of per-sample fractions of plasma cell isotypes, IgA^+^ and IgG^+^ (e), and naïve, memory, GC and unclassified (other) B cell subtypes (f) in the IIC stratified by disease severity. Fractions are computed with respect to the total number of all cells in each sample. P-values from Wilcoxon signed-rank test are reported. **g-h,** Representative flow cytometry plots to identify plasma cell types in NV, uninflamed UC and inflamed UC samples. **i-p,** Bar plots showing frequencies of plasma cells (i), IgA^+^ plasma cells (j), IgG^+^ plasma cells (k),B cells (l) and B cell types (naïve (m), switched memory (n), switched memory IgA^+^ (o) and switched memory IgG^+^ B cells (p) in healthy and inflamed samples. P-values from Kruskal-Wallis tests are reported. **p<0.01, *p<0.05.

**Extended Data Fig. 7:**
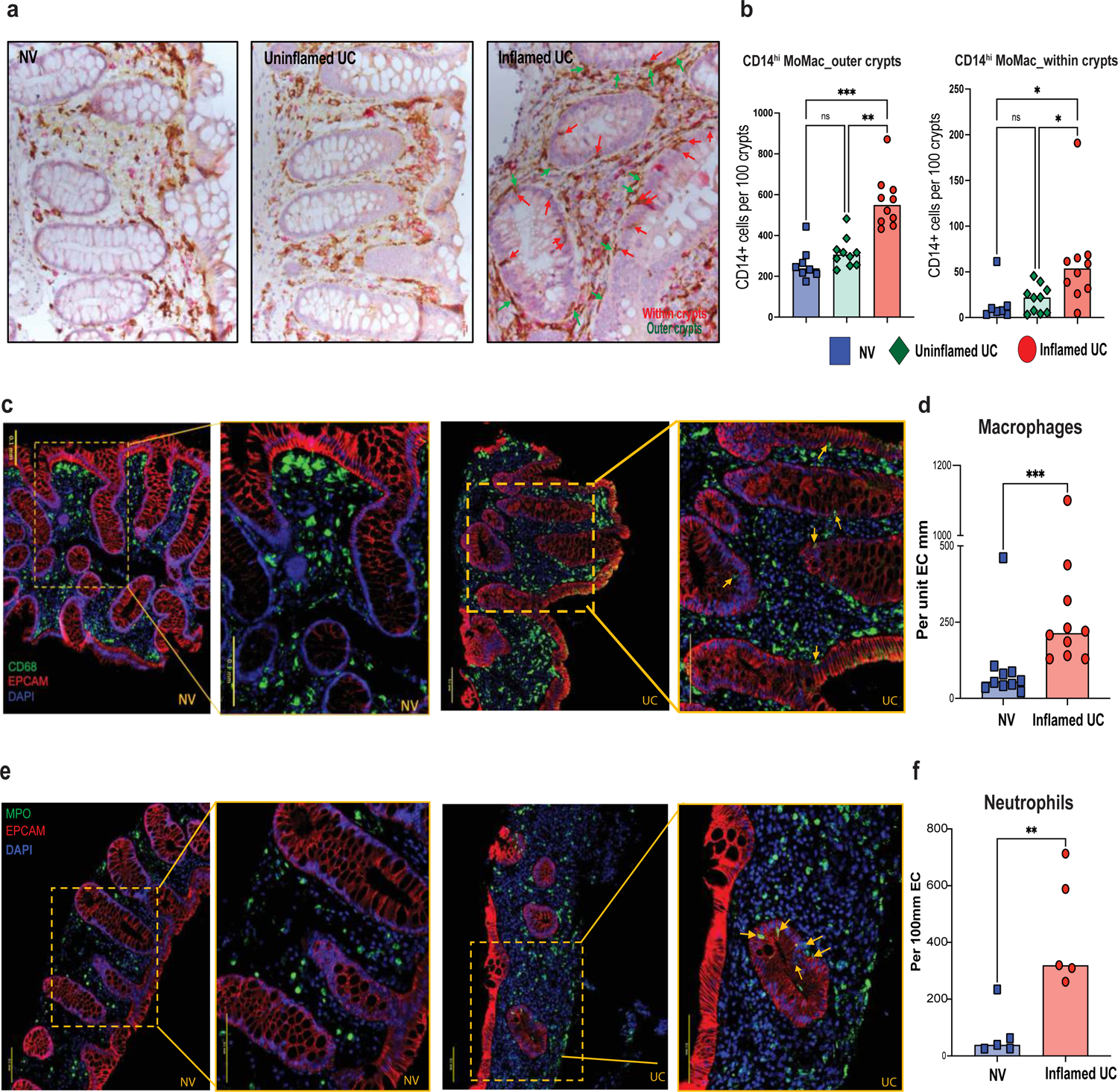
Neutrophils, monocytes and macrophages penetrate the colonic epithelium in patients with active UC. **a,** IHC staining showing the infiltration of CD14^+^ myeloid cells (in brown) within the EC of active UC patients (*right*) at the inflamed sites in comparison to the uninflamed sites (*middle*) and NV samples (*left*). Red arrows indicate the CD14^+^ cells entering the crypts and green arrows indicate the CD14^+^ cells surrounding the crypts. **b,** Bar plots comparing CD14^+^ cells surrounding the intestinal crypts (*left*) and CD14^+^ cells entering the crypts (*right*) between inflamed UC-, uninflamed UC- and NV-derived colonic tissues. P-values from one-way ANOVA test are reported. ***p<0.001, **p<0.01, *p<0.05, ns-p>0.05. **c,** IF staining depicting the expression of CD68 (green), EPCAM (red), DAPI (blue) in NV (*left, 10X and 20X magnification*) and UC (*right, 10X and 20X magnification*) samples. **d,** Bar plots comparing epithelium associated macrophages between inflamed UC and NV-derived tissues. P-value from Mann-Whitney test is reported. ***p<0.001. **e,** IF staining depicting the expression of MPO (green), EPCAM (red) and DAPI (blue) in NV (*left, 10X and 20X magnification*) and UC (*right, 10X and 20X magnification*) samples. **f,** Bar plot comparing epithelium-associated neutrophils between inflamed UC and NV-derived tissues. P-value from Mann-Whitney test is reported. **p<0.01.

**Extended Data Fig. 8:**
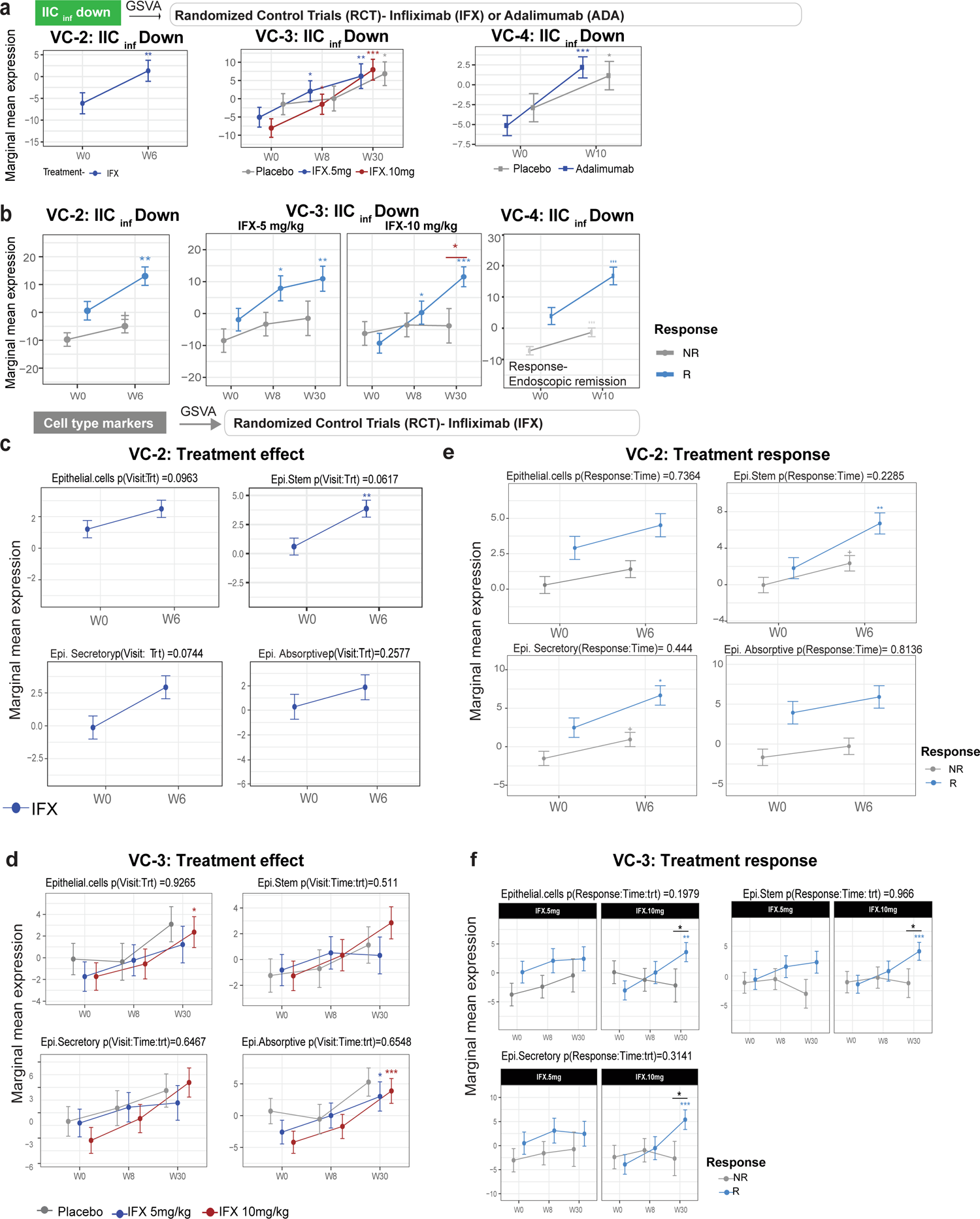
Down-regulated IIC_inf_ signature associates with non-response to anti-TNF therapy. **a,** Estimated marginal means (mean ± SEM) for the activity (GSVA scores) of the down-regulated IIC_inf_ signature at baseline and after 6 weeks (in VC-2) or 8 and 30 weeks (in VC-3) and 10 weeks (in VC-4) of anti-TNF therapy (Infliximab (IFX) in VC-2 and IFX 5 mg/kg and 10 mg/kg in VC-3); ADA in VC-4 and placebo. **b,** IFX-induced changes in down-regulated IIC_inf_ signature in responders (R) and non-responders (NR), where treatment response was defined by mucosal healing in VC-2, clinical response in VC-3 and endoscopic response in VC-4. P-values above the error bars denote significant change from baseline within the group at each time point, while p-values at the top indicate that treatment changes over time are significantly different between treatment groups (a-b) or R vs NR (c-d), * p< 0.05; ** p< 0.01; *** p< 0.001. **c-d,** Estimated marginal means (mean ± SEM) for the activity (GSVA scores) of the down-regulated Epi_inf_ subtype signature at baseline and after 6 weeks (**c**, in VC-2) or 8 and 30 weeks (**d**, in VC-3) of anti-TNF therapy (Infliximab (IFX) in VC-2 and IFX 5 mg/kg and 10 mg/kg in VC-3) and placebo. **e-f,** Effect of anti-TNF therapy on the activity (GSVA scores) of the epithelial cell subtype signatures in VC-2 (**e**) and VC-3 (**f**) in R and NR.

**Extended Data Fig. 9:**
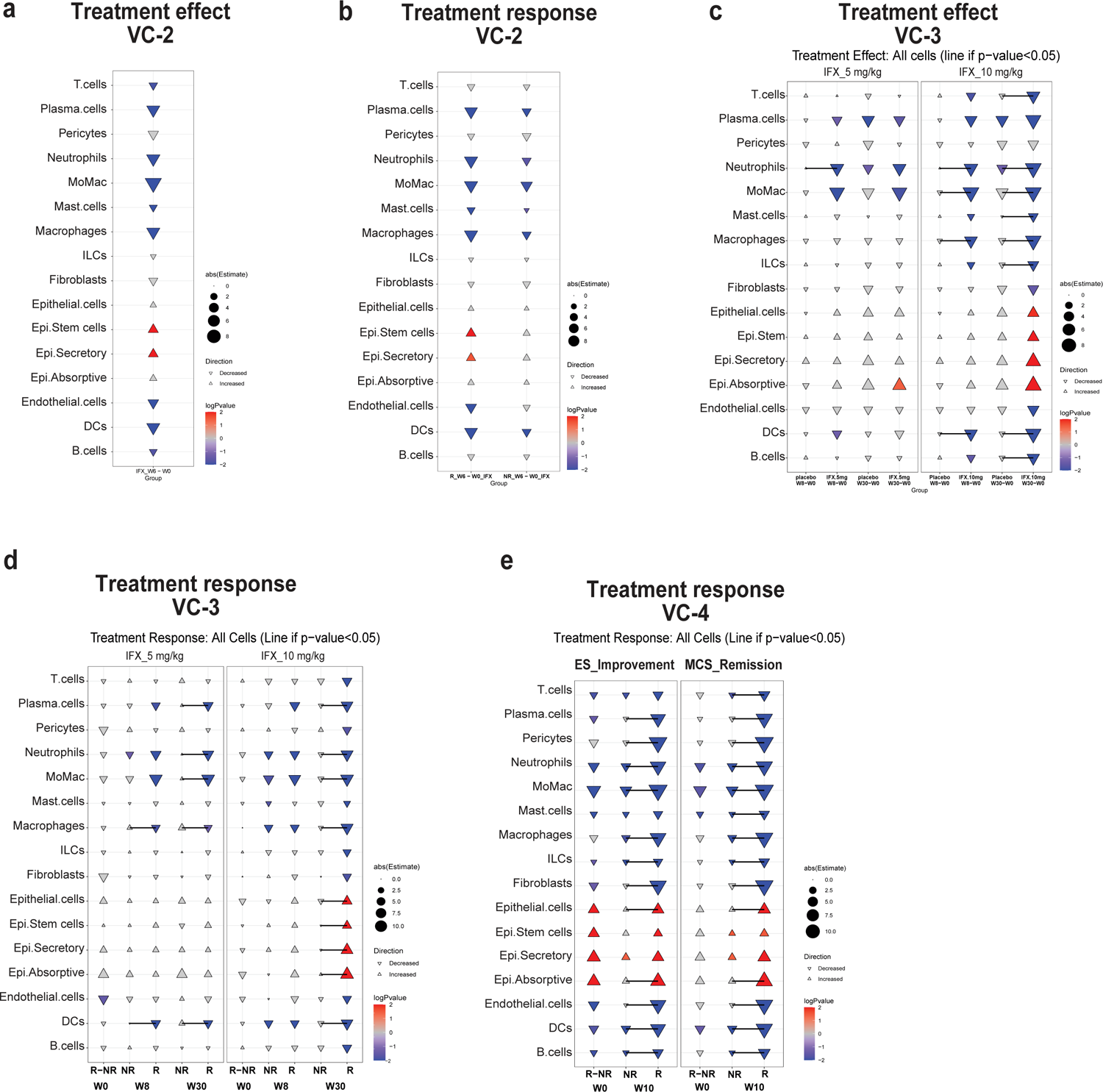
Effect of anti-TNF therapy on cell types in VC-2, VC-3 and VC-4 cohorts in patients on anti-TNF. **a,c**, Effect of anti-TNF therapy on the activity of cell-type signatures derived from scRNA-seq between active treatment and placebo in VC-2 **(a**) and VC-3 (**c**). **b,d,e,** Effect of anti-TNF therapy on the activity of cell-type signatures derived from scRNA-seq between responders (R) and non-responders (NR) in VC-2 (**b**), VC-3 (**d**) and VC-4 (**e**). Gene activity is assessed by GSVA scores from scRNA-seq derived cell type signatures. Connecting line shows p-value<0.05 between R and NR at respective timepoints.

## Data availability statement

All requests for raw and analyzed data and materials will be promptly reviewed by corresponding author and the study team. We will provide source data files for all the figures. Raw sequencing reads of scRNA-seq samples as well as UMI tables are available on the Gene Expression Omnibus (GEO) under accession number **GSE231817**) that may be accessible with the following token **cbefiwcgvvsxrwf**.

## Code Availability Statement

The R code developed for clustering and analyses in this study will be made available.

## Materials and Methods

### Primary cohort

Subjects with UC and NV (Primary cohort) were recruited, from the Inflammatory Bowel Disease (IBD) Center, the Gastroenterology Department, and the Endoscopy Unit at Mount Sinai Hospital. The protocol was approved by the Institutional Review Board #17-01304 and informed consent was obtained from all participants. Patients who were enrolled in the study were asked to provide research biopsies during the standard-of-care colonoscopy. All samples were fully deidentified before analyses. The public datasets derived from the patients will be made available after the acceptance of the paper. Clinical characteristics of patients with research biopsies in the Primary Cohort are displayed in Supplementary Table 1.

### Validation cohorts

#### Validation Cohort-1 (VC-1)^1^

GSE193677 series include RNAseq-based gene expression profiles from gut biopsies taken at endoscopy visit for 401 MSCCR UC patients and 243 non-IBD controls. Multiple biopsies from rectum and colonic regions were available for each patient, and they were defined as inflamed or non-inflamed based on endoscopy. Besides clinical and endoscopic disease severity measures, histological scores were also available for a subset of UC patients (n=178). Gene expression matrices from biopsy were generated from the count matrices using the voom transformation as previously described^1,2^. Expression profiles adjusted for technical variables (batch, rRNA rate, exonic rate and RIN score) using all the MSCCR cohort and after subsetting to our population of interest, used to generate signature scores.

#### Validation Cohort-2 (VC-2)^3^

GSE73661 series includes colonic gene expression (Affymetrix, HGU-1.0 ST) profiles from colonic biopsies from 23 UC patients before and 4-6 weeks after first infliximab (IFX) treatment. Response to therapy was defined as endoscopic mucosal healing (Mayo endoscopic score 0-1).

#### Validation Cohort-3 (VC-3)^4^

GSE23597 series includes mRNA expression of 113 mucosal colonic biopsies taken from 48 moderate-to-severe UC patients (6-12 Mayo score) at baseline and 8 and 30 weeks after treatment with 5 mg/kg (n=15) or 10 mg/kg (n=17) of infliximab or placebo (n=13). Patients were enrolled in the ACT1 randomized clinical trial (RCT, NCT00036439). Patients were considered responsive to therapy after 30 weeks if the total Mayo score decreased by at least 3 points from the baseline with an accompanying decrease in the sub-score for rectal bleeding of at least 1 point or an absolute sub-score for rectal bleeding of 0 or 1.

#### Validation Cohort-4 (VC-4)^5^

This cohort includes bulk RNAseq gene expression profiles of colonic biopsies collected from patients enrolled in the Hibiscus I trial (ClinicalTrials.gov: NCT02163759), a multicenter phase 3, randomized double-blind placebo-controlled and active-controlled study of etrolizumab, adalimumab and placebo in adult patients with moderately to severely acute ulcerative colitis (6-12 Mayo score, with endoscopic subscore >=2, and rectal bleeding and stool frequency subscores >=1). This dataset includes samples collected from patients in the placebo (n = 70) and adalimumab (n = 134) arms before treatment and at week 10 after treatment. MCS remission is defined as a Mayo clinic score of 2 or lower, a stool frequency and hybrid endoscopy subscore of 1 or lower and a rectal bleeding subscore of 0. Endoscopic improvement is defined as an endoscopic subscore of 1 or lower.

### Preparation of single cell suspension from pinch biopsies

Eight to 12 biopsies from inflamed and/or noninflamed parts of the colon of each patient were collected in RPMI on ice. Samples were processed within 2 h. Biopsies were first transferred in the dissociation medium consisting of 10 ml HBSS (free of calcium and magnesium) containing EDTA (0.5M, pH 8, Invitrogen) and HEPES (1M, Lonza) and incubated for 40 min at 37°C. The biopsies were then vortexed, washed with HBSS. After 40 min, the supernatant containing cells was removed, filtered through a cell strainer, spun down and resuspended. This cell suspension was referred to as the ‘epithelial cell’ fraction. To obtain LP cells, the remaining tissue was transferred into digestion media (10 ml of RPMI containing 0.005 mg of collagenase IV (Sigma-Aldrich) and DNAse I (Sigma-Aldrich). After an incubation of 40 minutes at 37°C in a rotating incubator (180 rpm), single cell suspension was made by mechanical dissociation, followed by subsequent filtering through a 100-µm and a 40-µm cell strainer and washed with RPMI twice. The digestion supernatant was then centrifuged, and the pellet was resuspended. This cell suspension was referred to as ‘LP cell fraction’. Both, the epithelial cell and LP fractions were resuspended into 500ul of RPMI (Gibco) containing 10% FBS. The single cell suspensions were then used for different assays as detailed.

### RNA extraction for IIC_bulk_ and Epi_bulk_ RNA sequencing

RLT plus buffer was added to the cell pellet obtained post EDTA dissociation (Epi_bulk_) and post-percoll (IIC_bulk_) and vortexed to obtain the lysate. The lysate was transferred to a gDNA eliminator spin column and centrifuged. Flow through was collected while the column was discarded. 1 volume of 70% ethanol was added to the flow-through and mixed well by pipetting. The sample was transferred to a RNeasy MinElute spin column and centrifuged. The flow-through was discarded and 700 µl Buffer RW1 was added to the RNeasy MinElute spin column, centrifuged. 500 µl Buffer RPE was added to the RNeasy MinElute spin column and centrifuged. Flow through was discarded and 500 µl of 80% ethanol added to the RNeasy MinElute spin column and centrifuged. The collection tube was discarded, and a dry spin was run for 5min to dry the membrane. RNeasy MinElute spin column was placed in a new 1.5 ml collection tube. RNase - free water was added, and the collection tube was centrifuged for 1 min at full speed to elute the RNA.

#### Pre-processing

Reads from FASTQ format files were mapped against Homo sapiens reference genome hg38 using STAR aligner. FastQC and RseQC were used to check quality of raw read and aligned reads. The mapping results were analyzed by featureCounts of the Subread package to quantify transcript reads. Features with fewer than 4 normalized counts across all samples were filtered out. The counts were normalized by using *voom^6,7^*, which estimates the mean variance relationship and generates a precision weight for individual normalized observations. Voom expression profiles were adjusted for RIN score using a linear model^8-10^ prior to statistical analysis.

### 3’ DGE RNA-sequencing

3′DGE RNA-sequencing was performed according to Cacchiarelli’s protocol. Briefly, libraries were prepared from 10 ng of total RNA obtained from *ex-vivo* stimulated biopsies. The mRNA poly(A) tail was tagged with universal adapters, well-specific barcodes and unique molecular identifiers (UMIs) during template-switching reverse transcriptase. Barcoded cDNA from multiple samples were pooled, amplified and tagmented using a transposon-fragmentation approach, which enriches for 3′ ends of cDNA. Libraries of 350–800 base pairs (bp) were sequenced with an Illumina HiSeq® 2500 using a TruSeq Rapid SBS Kit. Read pairs used for the analysis fulfilled the following QCs: all 16 bases of the first read must have quality scores of at least 10 and the first 6 bases must exactly align to a designed well-specific barcode. The second reads were aligned to RefSeq human mRNA sequences (GRCh37) using Burrows-Wheeler Aligner version 0.7.15.

### DGE RNAseq analysis for IICb_ulk_ and Epi_bulk_ RNA sequencing

DGE profiles were generated by counting for each sample the number of UMIs associated with each RefSeq genes. The raw count matrices were normalized with a variance-stabilizing transformation (vst) using the R package *DESeq2*. Differential gene expression analysis across culture conditions was performed with the *DESeq2* R package (version 1.30) and visualized using volcano plots. Genes were considered differentially expressed (DE) if adjusted p-values < 0.05. Each DE gene in the comparison of IL-17+IL-22 vs. medium conditions were centred and scaled across patients and the IL-17/IL-22 signature score was defined as the sum of their z-scores. R squared and p-values of the signature score to scaled gene correlations were computed with linear regression.

### Sample preparation for single-cell RNA sequencing

After single cell suspensions were prepared from the epithelial cell fraction (as described), the cell pellets were resuspended in 40% Percoll (BD Pharmigen) complemented with RPMI, 2% FCS, and separated by density gradient centrifugation in a discontinuous Percoll gradient (70%/ 40%) at 2000g for 20 min at 20°C. Cells were isolated from the 70-40% interphase were referred to as intraepithelial immune cells (IIC) in the study.

### 10x Processing

Single cell suspensions were counted using a Nexcelom Cellometer Auto2000 and each loaded on one lane of the 10x Genomics NextGem 3’v3.1 assay as per the manufacturer’s protocol with a targeted cell recovery of 8,000 cells. Gene expression libraries were prepared as per 10x Genomics demonstrated protocol^11^. Libraries were quantified via Agilent 2100 hsDNA Bioanalyzer and KAPA library quantification kit (Roche Cat# 0796014001) and sequenced at a targeted depth of 25,000 reads per cell. Libraries were pooled in equimolar concentration and sequenced on the Illumina NovaSeq 100 cycle kit with run parameters set to 28×8×0×60 (R1xi7xi5xR2).

### Single-cell RNA data processing and analysis

#### Filtering, Normalization, Batch correction

Cells were filtered based on the total number of UMI counts and mitochondria genes fractions. Only cells with total UMI counts greater than 1,000 and mitochondria gene fraction less than 10% were considered for the analysis. The analysis of single-cell RNA data including data normalization and batch effect correction was performed via the Seurat package^12, 13^. First, each cell was normalized independently using function *NormalizeData* which divides feature counts for each cell by the total counts for that cell. Data was then natural log transformed. Next, we found variable features using function FindVariableFeatures^12, 13^. This function first fits a line to the relationship of log(variance) and log(mean) using local polynomial regression; then, standardizes the feature values using the observed mean and expected variance (given by the fitted line). Feature variance was then calculated based on the standardized values. Finally, we selected features which were repeatedly variable across datasets for integration via function *SelectIntegrationFeatures*^12, 13^. We then identified anchors using the *FindIntegrationAnchors* function, which takes a list of Seurat objects as input, and use these anchors to integrate different datasets together via the *IntegrateData* function^12, 13^. Dimensionality reduction to identify anchors was performed using reciprocal principal component analysis. For visualization, the dimensionality of each dataset was further reduced using Uniform Manifold Approximation and Projection (UMAP) implemented with Seurat functions *RunUMAP*. The PCs used to calculate the embedding were as the same as those used for clustering.

#### Clustering

Clustering was performed via function *Find.Cluster* available in the Seurat package, with resolution parameter r=0.5. This function identifies clusters of cells using a shared nearest neighbor algorithm. Once that unsupervised clustering was performed, we identified cluster-specific markers using function *FindMarkers* available in the *Seurat* package^12, 13^. Markers were ranked based on area under the receiver operating characteristic curve (AUC). Based on the cluster-specific markers and leveraging existing signatures from the literature^14^, clusters were annotated into 10 cell-types^12^.

#### Cell-type signatures

We considered the lineage cell-type categories to obtain gene-signature from the single-cell RNA data. This was performed via function *FindMarkers* available in the Seurat package. Only markers with an AUC greater than 0.7 were considered as cell-type specific markers. Mitochondrial and ribosomal protein genes were removed in the final list of genes since they were found expressed in multiple cell-types^12^.

#### Re-clustering of lymphoid and myeloid compartments

T-cells, B-cells, Plasma and Myeloid cells were re-clustered as smaller individual subsets using the same nearest neighborhood clustering algorithm^12^. The resolution parameters used in re-clustering were 0.5 for T cells, Plasma and Myeloid cells and 0.3 for B cells. After identifying markers for each of the subclusters we could assign most of them to specific subtypes of each cell type: Neutrophils, Macrophages, Monocyte-Macrophages and DCs for Myeloid; Memory, Naïve, Cytotoxic, T_REG_, Gamma-Delta (γδ) and T_H17_ for T cells; Memory, Naïve, and Germinal Center (GC) for B cells; and IgA and IgG isotypes for Plasma cells.

#### Pathway enrichment analysis

For each cell-type compartment (i.e., T, Myeloid and Plasma cells), the log fold-change was computed between the average gene expression in cells belonging to UC and that of cells belonging to NV samples. Enrichment scores and adjusted p values (FDR) were derived via gene set enrichment analysis^15^ for UC vs NV and NV vs UC log fold-change separately Specifically, genes were ordered in decreasing order of fold-change and a Kolmogorov-Smirnov statistics was utilized to find pathways significantly enriched at the top of the list. A permutation-based strategy was utilized to derive p-values and FDR for different pathways^15^. Molecular pathway gene sets used in this analysis were from KEGG^16^, Reactome^17^ and BioPlanet^18^ databases.

### Spatial transcriptomic sample collection, data processing and analysis

#### Sample preparation and preprocessing

Biopsy specimen from inflamed and/or noninflamed parts of the colon of each patient were collected into OCT medium in cryomolds and snap frozen over dry ice during the colonoscopy procedure. Additional OCT medium was added to cover the cryomolds. Once frozen, samples were transported on dry ice and stored at −80 until further use. For the Visium Gene Expression library preparation, each frozen section was mounted on Tissue Optimization Slide and permeabilized for increasing times to determine optimal permeabilization incubation following the manufacturer’s instructions (CG000238 Rev E, 10X Genomics). The sections were fixed in pre-chilled methanol for 30 min, H&E stained and imaged using Aperio AT2 scanner (Leica Biosystems) 20X magnification. The sections were then permeabilized for 10 min and spatially tagged cDNA libraries were constructed using the 10x Genomics Visium Spatial Gene Expression 3’ Library Construction V1 Kit (CG000239 Rev F, 10X Genomics). 10 μl of amplified cDNA from each sample was used for library construction and has undergone fragmentation, adapter ligation, index PCR, and purification. Each final library was sequenced on an Illumina NextSeq using 300 cycle high output kits with sequencing depth of ∼25,000 reads per spot. Sequencing data was mapped to GRCH38-2020-A reference transcriptome using the Space Ranger software (Space ranger v1.3) to derive a feature spot-barcode expression matrix.

#### Mapping, filtering, normalization

The four Visium data sets (two UC and two NV) were initially processed by 10x Space Ranger. Higher level analysis and visualizations were performed by *Giotto* R package^19^. In each sample the spots with fewer than 500 unique genes were removed. Genes present in fewer than 30 spots were removed. Expression levels were normalized using default *Giotto* procedure.

#### Cell Type Enrichment

Enrichment scores were computed on normalized expression levels in each spatial spot independently using Kolmogorov-Smirnov statistics^15^. Enrichment scores were derived based on signatures from scRNA-seq data. Cell-type specific markers from scRNA-seq data with an AUC greater than 0.70 were considered to derive enrichment scores (Supplemental Table 8). In addition to these cell-types derived based on our scRNA-seq data, we considered different types of epithelial cells including stem cells, absorptive and secretory. Specifically, we used the gene sets-absorptive_all, secretory_all and stem cells from the epithelial subset^14^ (Supplementaryl Table 8).

#### Spatial deconvolution

In addition to enrichment analysis, we performed cell type deconvolution via spatialDWLS^20^. For the deconvolution, we utilized the same gene signatures described in the enrichment analysis. In addition to gene signatures, *spatialDWLS* requires as input the gene expression profile of different cell types. Briefly, we utilized the gene expression profiles of different cell-types from our scRNA-seq data. For different types of epithelial cells, we re-clustered the epithelial subset of our scRNA-seq data the same way as described for lymphoid and myeloid compartments, with resolution parameter 0.7. We obtained 15 epithelial sub-clusters and mapped some of them to epithelial absorptive, epithelial stem cells and epithelial secretory cells based on elevated levels of corresponding gene set signatures from the literature^14^ (absorptive_all, secretory_all and stem cells from the epithelial subset, Supplementary Fig 5b). Epithelial sub-clusters 0, 5 and 10 were annotated as Absorptive, sub-clusters 4,6 and 9 as Secretory and sub-clusters 3 and 12 as Stem cells. The gene expression profiles corresponding to these sub-clusters were utilized to perform the deconvolution of ST (Supplementary Table 8).

#### Cell-cell colocalization

The identification of cell types frequently co-localized together was performed using *cellProximityEnrichmentSpots* function in *Giotto* package^19^. As input, we used the deconvoluted fractions of different cell types in each spot. Spatial network was used to quantify cell type fractions occurrence in neighboring spots. We performed 1000 random permutations of spot locations to estimate the background co-localization distribution. The significance of the co-localization (p-value for each pair of cell types) was computed as the fraction of random permutations co-localized more frequently than the observed value.

### IIC and Epi bulk RNA-seq data analysis

#### Differential Expression Analysis

Principal component analysis (PCA) was used for exploratory analysis. Expression profiles were modeled using linear mixed-effects models in the limma framework. Models included group variables (Inflamed, uninflamed and normal volunteers) in IIC and Epi as fixed effects, and a random intercept for each subject. Hypotheses of interests were tested using contrasts and p-values from the moderated t-test were adjusted for multiple hypotheses using the Benjamini-Hochberg approach, which controls the FDR. Genes were considered differentially expressed, if false discovery rate (FDR) <0.05; fold change (FCH) >2.0. <u>IIC_inf_</u> and <u>Epi_Inf_</u> signatures consist of DEG between inflamed and uninflamed samples of IIC and Epi compartments, respectively.

#### Pathways and cell type Enrichment analysis

Gene sets were tested for over-representatin analysis using clusterProfiler 3.16.1, p-values from the Fisher’s exact test were adjusted using Benjamini-Hochberg (BH) multiple test correction. The collection of genesets included KEGG^16^ pathways, cell-types signatures from our scRNA-seq experiment and a published dataset^14^.

#### Defining IIC_inf_ IFX-response signature

In VC-3, expression profiles were modeled using linear mixed-effect models with the interaction between time and treatment group (5mg /10mg dose of IFX and placebo). Treatment DEG were identified as those with significant changes in expression compared to baseline (FDR<0.05 and FCH>1.5) at any time point in each treatment group. The difference between responder and non-responder to anti-TNF was assessed for treatment DEGs, using a linear model with interaction between the time, treatment dose, and response. The IFX - response signature thus included genes where the IFX-induced changes in the direction of disease resolution were significantly different between responders (R) vs non-responders (NR) at either week 8 or week 30 (FDR<1.5). *IIC_inf_ IFX-response signature* was thus defined as IIC_inf_ genes that were IFX-responsive by the criteria above and included 159 genes up-regulated (96 down-regulated) in inflamed gut and down-regulated (up-regulated) with treatment.

#### Scoring methods

We estimated the overall expression activity of various gene signatures in the VC-1-4 using gene set variation analysis (GSVA). For each gene signature (IIC_Inf,_ scRNA-seq based cell type markers and subclusters), GSVA, a non-parametric and non-supervised method, estimated a sample-wise enrichment score for each gene set on the validation cohorts. Such enrichment scores, representing overall gene set activity, were then used for hypothesis testing with respect to phenotype information as described in the sections below.

#### Association of IIC_Inf_ activity with UC severity in VC-1

Linear mixed-effect model was used with region, disease status and tissue type (inflamed vs non-inflamed) with a random intercept of patient. Association between gut signature scores with clinical and endoscopic severity for MSCCR UC patients were analyzed with a mixed-effect model including clinical activity or disease severity, age, sex and region as a fixed-effects. Clinical disease activity measures were based on Clinician-based Simple Clinical Colitis Activity Index (SCCAI), clinically inactive disease defined as an SCCAI < 5 and active disease as SSCAI >= 5. The Mayo endoscopic scores were used to categorize UC endoscopic severity as: normal/inactive disease (0); mild disease (1); moderate disease (2); or severe disease (3). Pearson correlation between histology scores (Nancy Index) with signature scores was estimated. The expressions of genes in the signatures were scaled as z-scores in MSCCR data before examining the correlation with Nancy index in inflamed UC biopsies.

#### Association with anti-TNF/ADA response in VC-2, VC-3, VC-4

The effect of IFX on the activity (GSVA-scores) of each gene signature (IIC_Inf_, cell-types and subcluster) was compared between active treatment and placebo (using VC-2 and VC-3) and between week 30 responders and non-responders (VC-3). In the treatment effect analysis, changes in GSVA scores across time in different treatment groups (IFX 5mg/kg, IFX 10 mg/kg and placebo) were estimated by the linear mixed-effect model (LMEM) with visit, treatment and its interaction as fixed effects, and random intercept for each subject. Differences in IFX-induced activity between anti-TNF responders and non-responders was evaluated via LMEM with a three-way interaction of visit, treatment dose, and response status as fixed effects, and a random intercept for each subject. Similarly, in the treatment effect analysis for VC-4, the difference between Adalimumab-induced activity between responders and non-responders for each gene set was evaluated through a LMEM with visit, response, and their interaction term as fixed terms and a random intercept for each subject.

### Statistical analysis

Statistical analysis was conducted in R version 4.1.3.

### Flow cytometry

Multi-parametric flow cytometry was performed on intestinal cells from patients with UC and controls, sampling the non-inflamed and inflamed sites. The gating strategy for colonic T cells, myeloid cells, plasma cells and B cells are shown in Supplementary Fig. 2 a-d respectively. Single cell suspensions were incubated with PBS containing the flow antibody cocktail for 30 min at 4°C in dark. Cells were washed twice with PBS and then fixed using 2% formaldehyde for 30 min at 4°C. Finally, cells were harvested in PBS before acquisition. Dead cells and doublets were excluded from all analyses.

### Immunofluorescence staining

5 μm thick sections were cut from formalin-fixed, paraffin-embedded tissue blocks. These sections were dewaxed twice in 100% xylene for 5 min. These were then rehydrated in graded alcohol starting from 100% ethanol, 90% ethanol, 80% ethanol and then 70% ethanol with 20 dips in each of the alcohol. The sections were then washed in 1X PBS twice for 5 min each. Heat-induced epitope retrieval was performed by incubating slides in a pressure cooker for 15 minutes on in target retrieval solution. Once slides cooled to room temperature, they were washed twice in PBS. Nonspecific binding was blocked with 10% goat serum for 1 hour at room temperature. Sections were incubated in primary antibodies diluted in blocking solution overnight at 4°C. Slides were washed in 0.1% Tween-20 plus PBS thrice and then incubated in secondary antibody and DAPI (4′,6-diamidino-2-phenylindole, 1 μg/mL) for 1 hour at room temperature. Sections were washed twice in 0.1% Tween 20 plus PBS and once in PBS and then mounted with Fluoromount-G. Controls included omitting the primary antibody (no primary control) or substituting primary antibodies with nonreactive antibodies of the same isotype (isotype control). Tissue was visualized and imaged using a Nikon Eclipse Ni microscope and digital SLR camera (Nikon, DS-Qi2).

### *Ex vivo* stimulation of colonic biopsies

Colonic biopsies (n=8) for *ex vivo* stimulations were collected in unaffected areas obtained after surgical resection of colorectal tumors (at least 10 cm distant to the tumor). Written informed consents were obtained under institutional review board protocol # DC-2008-402. Biopsies were transported in ice-cold RPMI and processed within a maximum of 2h. Biopsies were controlled for weight and placed into 4-well Petri dishes filled with in 500µL serum-free medium (RPMI 1640, Gibco^TM^) supplemented with BSA (0.01%), 200µg/mL Penicillin/Streptomycin (Gibco ^TM^; #15140-122) and 0.25µg/mL Fungizone (Gibco^TM^; #15290-026). Biopsies were cultured during 6 h at 37°C in a 95% O_2_ / 5% CO_2_ atmosphere on a low-speed rocking platform in the presence of cytokines (IL-22; 100ng/mL [Miltenyi, # 130-096-295], IL-17; 100ng/mL [Miltenyi, #130-093-959], IL-22BP; 200ng/mL [Miltenyi, #8498-BP-025]). Supernatants were collected and stored at −80°C until use.

### DGE RNAseq analysis for ex-vivo stimulation

DGE profiles were generated by counting for each sample the number of UMIs associated with each RefSeq genes. The raw count matrices were normalized with a variance-stabilizing transformation (vst) using the R package *DESeq2*. Differential gene expression analysis across culture conditions was performed with the *DESeq2* R package (version 1.30) and visualized using volcano plots. Genes were considered differentially expressed (DE) if adjusted p-values < 0.05. Each DE gene in the comparison of IL-17+IL-22 vs. medium conditions were centered and scaled across patients and the IL-17/IL-22 signature score was defined as the sum of their z-scores. R squared and p-values of the signature score to scaled gene correlations were computed with linear regression.

