## Supplementary material for "Myeloid cell influx into the colonic epithelium is associated with disease severity and non-response to anti-Tumor Necrosis Factor Therapy in patients with Ulcerative Colitis": Legends for Supplementary materials

### Supplementary figure legends

**Supplementary Fig. 1:** Distribution of per-sample cell type fractions in the two intestinal compartments, intraepithelial immune cells (IIC) and lamina propria (LP).

**a**, Boxplots showing the distribution of per-sample fractions of the major cell types in IIC (upper) and LP (lower).

**b-c**, Boxplots of per-sample fractions of T cell types in IIC (b) and LP (c) stratified by UC/NV status.

**d-e**, Per-sample fractions of myeloid cell types in IIC (d) and LP (e),

**f**, Per-sample fractions of plasma cell types in IIC and LP.

**g**, Boxplot of per-sample fractions of B cell types in IIC and LP. All fractions are computed with respect to the total number of cells for each sample. Boxplots are stratified based on disease status (NV = blue, UC = red). P-values from Wilcoxon signed-rank test are reported.

**Supplementary Fig. 2:** Flow cytometry-based gating strategy for annotating T cells, myeloid cells, plasma cells and B cells.

**a,** Gating strategy for T cell types- αβ T cells, γδ T cells, CD4+ αβ T and CD8+ αβ T cells.

**b,** Gating strategy for myeloid cell types- eosinophils, neutrophils, $CD14^{hi}$ MoMac and $CD16^{hi}$ MoMac.

**c,** Gating strategy for plasma cell types- $IgG^{+}$ and $IgA^{+}$ plasma cells.

**d,** Gating strategy for B cell types- Memory, naïve and switched memory B cells.

**Supplementary Fig. 3:** Plasma cell and B cell distribution in LP and pathway enrichment analysis in IIC-plasma cells and IIC-B cells in UC.

**a-b,** Boxplots showing the distribution of per-sample relative fractions of plasma cell sub-clusters (a, P0-P9) and B cell sub-subclusters (b, B0-B5) in LP stratified by disease severity. Fractions are computed with respect to all plasma cells (a) or B cells (b) in each sample. P-values from Wilcoxon signed-rank test are reported.

**c-d,** Boxplots of per-sample fractions of $IgA^{+}$ and $IgG^{+}$ plasma cells (c) and B cell types- naïve, memory, germinal center B cells (d) in LP, stratified by disease severity. Fractions are computed with respect to the total number all cells in each sample. P-values from Wilcoxon signed-rank test are reported.

**e-f,** Pathway enrichment analysis from differential analysis between UC versus NV in plasma cells (e) and B cells (f) in IIC-enriched samples. The bubble plot on the left side of the figures shows the summary statistics from pathway analysis; with the size of the bubble corresponding to the enrichment score (ES) while the bubble color to the signed -log10 adjusted p-value. The heatmaps at the right side show the log fold change between UC and NV the top five leading edge genes in each pathway.

**Supplementary Fig. 4:** Sample images and distribution of enrichment scores and deconvolution factors in the inflamed UC and NV samples analyzed using spatial transcriptomics.
**a,** Hematoxylin and eosin (H&E) staining images of UC and NV samples used for spatial transcriptomic analysis.
**b-c,** Per-sample distribution across spatial spots of enrichment scores (b) and cell-type fractions inferred via deconvolution analysis (c) for B cells, plasma cells, T cells, neutrophils, ILCs, mast cells, pericytes, fibroblasts and endothelial cells.

**Supplementary Fig. 5:** Enrichment of the epithelial subtypes from a published dataset[14] in the scRNA-seq derived cell types and clusters.
**a,** Violin plots showing the average z score of marker gene sets defining different epithelial subtypes in the scRNA-seq derived cell types.
**b,** Violin plots showing the mean z score of marker gene sets associated with epithelial subtypes in the scRNA-seq derived epithelial cell clusters.

**Supplementary Fig. 6:** CD14 (brown) IHC staining comparing inflamed UC, uninflamed UC and NV tissues. All the images that were used in the quantitative microscopy are shown.

**Supplementary Fig. 7:** IF staining depicting the expression of CD68 (green), EPCAM (red) and DAPI (blue), in inflamed UC, uninflamed UC and NV tissues. All the images that were used in the quantitative microscopy are included.

**Supplementary Fig. 8:** IF staining depicting the expression of MPO (green), EPCAM (red) and DAPI (blue) in inflamed UC, uninflamed UC and NV tissues. All the images that were used in the quantitative microscopy are included.

**Supplementary Fig. 9:** Enrichment of treatment responsive gene signature in the cell types derived from scRNA-seq data.

**a,** $IIC_{inf}$ bulk seq based treatment response signature- Venn diagram showing intersection of genes that were differentially expressed in inflamed UC samples ($IIC_{inf}$) and reversed with anti-TNF treatment response and the corresponding cell types. 159 DEGs that were higher in expression in inflamed UC samples showed reduced expression with anti-TNF response while 96 DEGs that had lower expression in inflamed samples showed increased expression with anti-TNF response.

**Supplementary Fig. 10:** Enrichment of treatment response associated genes in the list of cell type markers derived based on scRNA-seq data. Barplot shows p-values (-log10 scale) from Fisher's exact test showing the association of scRNA-seq cell type signature (PC) with anti-TNF response (VC-3, W30, 10mg IFX). This analysis was performed considering:
**a,** the major cell types in our scRNA-seq data
**b,** the granular list of cell types in our scRNA-seq data
**c,** immune cell types from a published dataset by Smillie et al[14].

**Supplementary Fig. 11:** Cell type association with anti-TNF therapy in VC-2 cohort.
**a,** Effect of the treatment on the cell types in patients on anti-TNF therapy compared to those on placebo.
**b,** Effect of the treatment on the cell types (B cells, myeloid cells, epithelial subtypes, T cells, ILCs, fibroblasts, pericytes and endothelial cells) in patients responding to anti-TNF therapy in VC-2.

**Supplementary Fig. 12**: Cell type association with anti-TNF therapy in VC-3 cohort.
**a,** Effect of the treatment on the cell types in patients on anti-TNF therapy compared to those on placebo.
**b,** Effect of the treatment on the cell types (B cells, epithelial subtypes, T cells, ILCs, fibroblasts, pericytes and endothelial cells) in patients responding to anti-TNF therapy (5 mg/kg and 10 mg/kg IFX) in VC-3. Plots for myeloid cell types, plasma cells and epithelial absorptive cells included for IFX 10mg/kg dose.

**Supplementary Fig. 13**: Cell type association with anti-TNF therapy in VC-4 cohort.
**a,** Effect of the treatment on the cell types in patients on anti-TNF (adalimumab) therapy and on placebo.

**b,** Effect of the treatment on the cell types (B cells, myeloid cells, epithelial subtypes, T cells, fibroblasts, pericytes and endothelial cells) in patients responding to anti-TNF therapy (adalimumab) in VC-4.

**Supplementary Fig. 14:** Pathway enrichment analysis (KEGG[16] 2021)-
**a,** KEGG pathway enrichment analysis from bulk RNA sequencing derived inflammation signature (IIC-specific and IFX-response signature),
**b**, pathway enrichment analysis on the leading-edge genes from the cell types (plasma cells, epithelial cells, neutrophils, MoMac, DC and macrophages) that were significantly different in responders vs non-responders in patients on anti-TNF therapy (VC-3).

**Supplementary Table Legends**

**Supplementary Table 1.** Detailed clinical details for each of the patients included in the primary cohort (PC) for
**a,** ulcerative colitis (UC) patients and
**b,** normal volunteers (NV).
**c,** shows the list of abbreviations used.

**Supplementary Table 2.** Bulk-RNA sequencing description-
**a,** Differential gene expression analysis in epithelial compartment (Epi) between a, Inflamed vs uninflamed UC; b, inflamed vs NV and; c, uninflamed vs NV
**b,** Differential gene expression analysis in intraepithelial immune cells (IIC) between a, Inflamed vs uninflamed UC; b, inflamed vs NV and; c, uninflamed vs NV
**c,** Cell types for DEG (upregulated and downregulated) in Epi-inf and IIC-inf DEG
**d,** Over-representation analysis for the Epi-inf and IIC-inf signatures of the selected pathways

**Supplementary Table 3.** Single-cell RNA sequencing derived cluster description and cell type annotations.
**a,** cell type counts;
**b,** Cluster cell type annotation;
**c,** gene markers for primary clusters;

**d,** Mean gene expression in primary clusters;

**e,** fractions for cell types in each fraction.

**Supplementary Table 4.**

**a,** Single-cell RNA sequencing derived T cell sub-cluster description and T cell type annotations;

**b,** Relative fractiob (relative to T cell compartment) of subclusters within each sample (patient);

**c,** Total sub-cluster cell counts for sample types (UC vs NV), disease severity (NV, moderate or severe) and tissue type (IEL vs LP);

**d,** Cell type assignments for each of the 10 T cell subclusters;

**e,** Fraction of cell types (relative to all cells) within each sample (patient) for the 6 T cell types;

**f,** Mean gene expression in each of the T cell subclusters;

**g,** Gene markers for each T cell subcluster, significance cutoff AUC >= 0.65;

**h,** ssGSEA scores for pathways enriched in UC vs NV in T cells;

**Supplementary Table 5.** Differential gene expression in ex-vivo stimulation RNA sequencing experiment.

**a,** IL-22 vs medium only;

**b,** IL-17 vs medium only and

**c,** IL-17+Il-22 vs medium only

**Supplementary Table 6.** Single-cell RNA sequencing derived myeloid cell sub-cluster description and myeloid cell type annotations.

**a,** Relative fraction (relative to myeloid cell compartment) of subclusters within each sample (patient);

**b,** Total subcluster cell counts for sample types (UC vs NV), disease severity (NV, moderate or severe) and tissue type (IEL vs LP);

**c,** Cell type assignments for each of the 10 myeloid subclusters;

**d,** Fraction of cell types (relative to all cells) within each sample (patient) for the 4 myeloid cell types;

**e,** Mean gene expression in each of the myeloid subclusters;

**f,** Gene markers for each myeloid subcluster, significance cutoff AUC >= 0.65;

**g,** ssGSEA scores for pathways enriched in UC vs NV in myeloid cells;

**Supplementary Table 7.** Single-cell RNA sequencing derived plasma cell and B cell sub-cluster description and annotations.

**a,** Relative fraction (relative to B cell compartment) of B cell subclusters within each sample (patient);

**b,** Relative fraction (relative to Plasma cell compartment) of Plasma cell subclusters within each sample (patient);

**c,** Total subcluster cell counts for sample types (UC vs NV), disease severity (NV, moderate or severe) and tissue type (IIC vs LP);

**d,** Cell type assignments for each of the B and Plasma cell subclusters;

**e,** Fraction of cell types (relative to all cells) within each sample (patient) for the 4 B cell and 2 Plasma cell subtypes;

**f,** Mean gene expression in each of the B and Plasma cell subclusters;

**g,** Gene markers for each B and Plasma cell subcluster, significance cutoff AUC >= 0.65;

**h,** ssGSEA scores for pathways enriched in UC vs NV in B cells;

**i,** ssGSEA scores for pathways enriched in UC vs NV in plasma cells;

**Supplementary Table 8.** Spatial transcriptomics derived spatial spots description, cell type assignments and colocalization analysis.

**b,** Enrichment scores for each cell type signature in each spatial spot in the four samples;

**c,** Cell type deconvolution fractions in each spatial spot in the four samples;

**d,** Cell-cell colocalization permutation analysis output, sample IBDUC151;

**e,** Cell-cell colocalization permutation analysis output, sample IBDUC152;

**f,** Cell-cell colocalization permutation analysis output, sample NV163;

**g,** Cell-cell colocalization permutation analysis output, sample NV164;

**h,** Matrix with gene set and mean cell type expression values used for deconvolution with spatialDWLS;

**i,** Enrichment scores for IL17+IL22.vs.medium.up gene set in each spatial spot;

**j,** List of cell type markers used for deconvolution, derived from scRNAseq + 3 epithelial cell types from published dataset.

**Supplementary Table 9.** RNA sequencing based anti-TNF response signature describing the set of genes that associated with inflammation and reversed with treatment in responders vs non-responders in VC-3.

**Supplementary Table 10.** RNA sequencing based pathway enrichment analysis from IFX-treatment response gene signature using

**a,** mSig-Hallmark database;

**b,** KEGG-2021 database and

**c,** on the leading edge genes from scRNA-seq derived cell types using both mSig-Hallmark and KEGG-2021 databases.
