## Supplementary Figures for "Myeloid cell influx into the colonic epithelium is associated with disease severity and non-response to anti-Tumor Necrosis Factor Therapy in patients with Ulcerative Colitis"

**Supplementary Fig. 1: Distribution of per-sample cell type fractions in the two intestinal compartments**

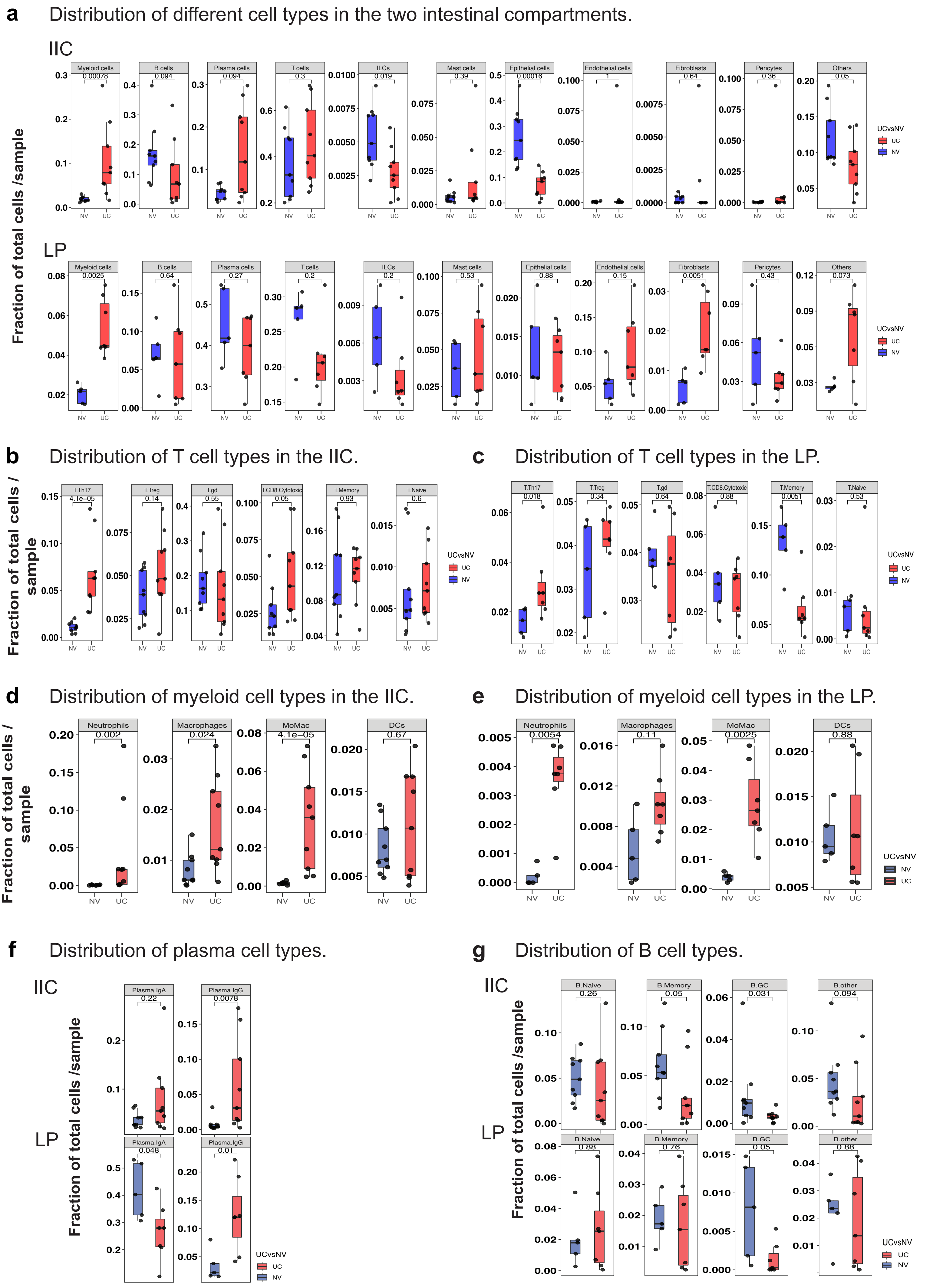

**Supplementary Fig. 2: Flow cytometry based gating strategy for annotating myeloid cells, T cells, plasma cells and B cells.**

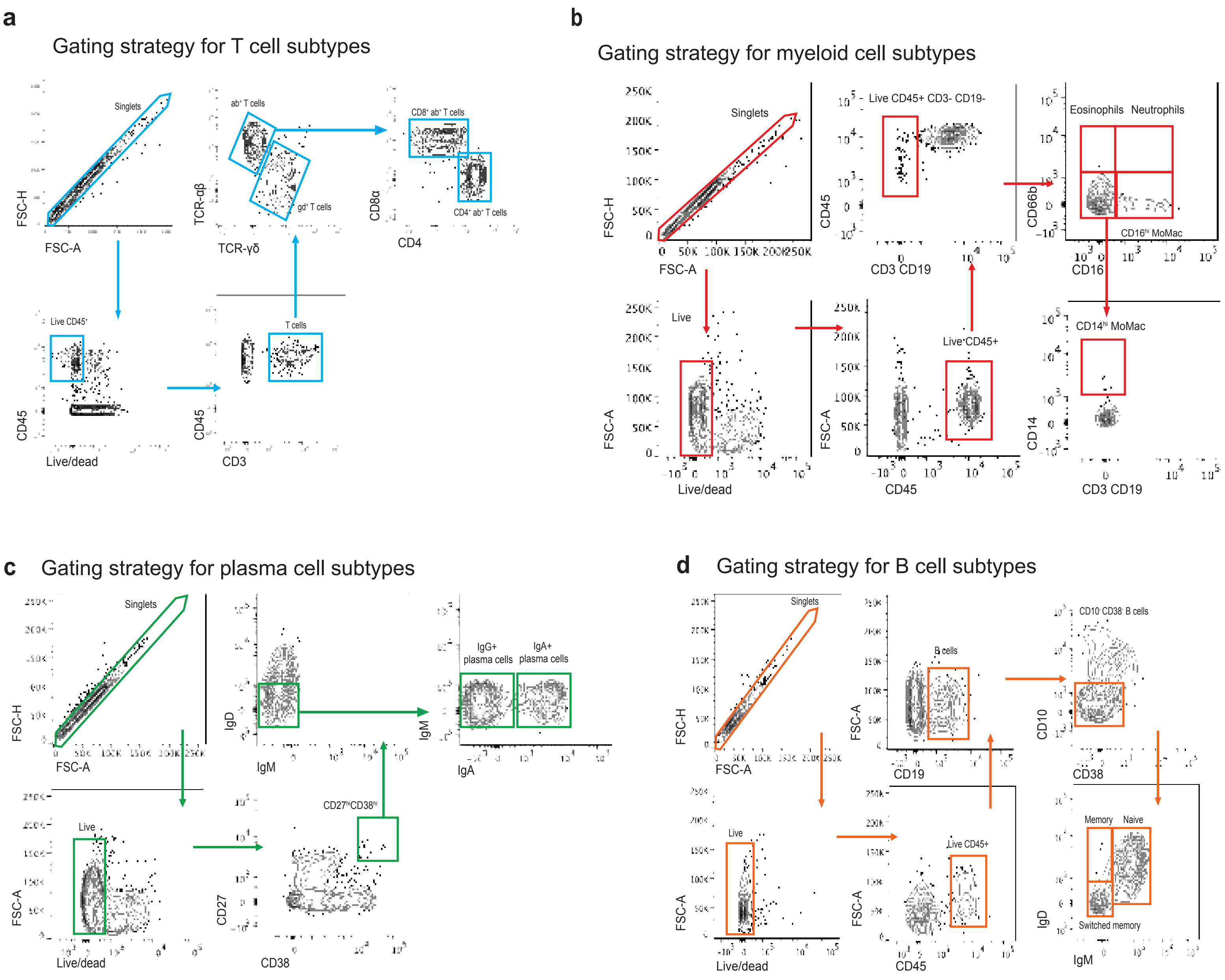

#### Supplementary Fig. 3: Plasma cell and B cell distribution in LP and pathway enrichment analysis in IIC-plasma cells and IIC-B cells in UC.

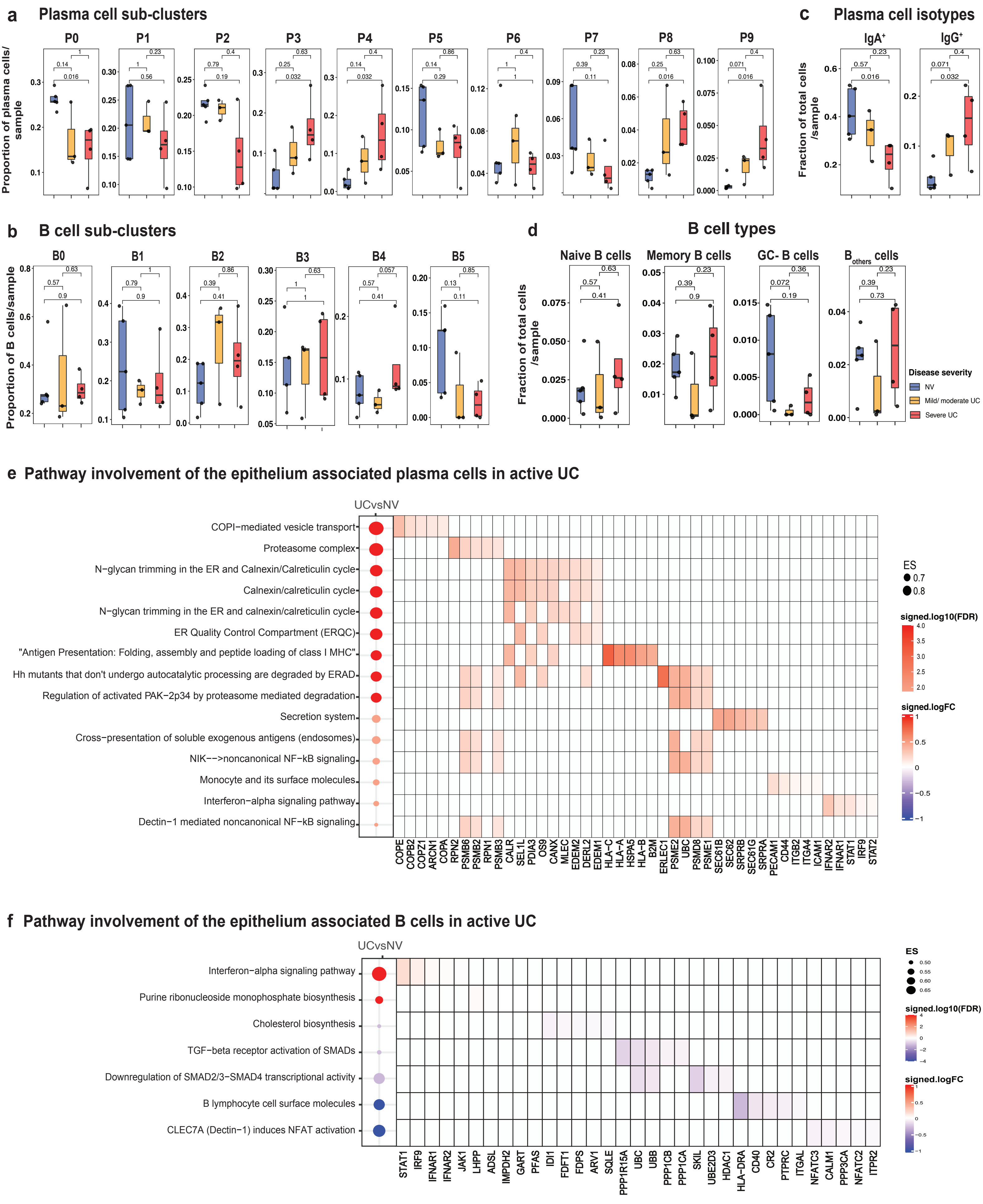

**Supplementary Fig. 4: Sample images and distribution of enrichment scores and deconvolution factors in the active UC and NV samples analysed using spatial trancriptomics.**

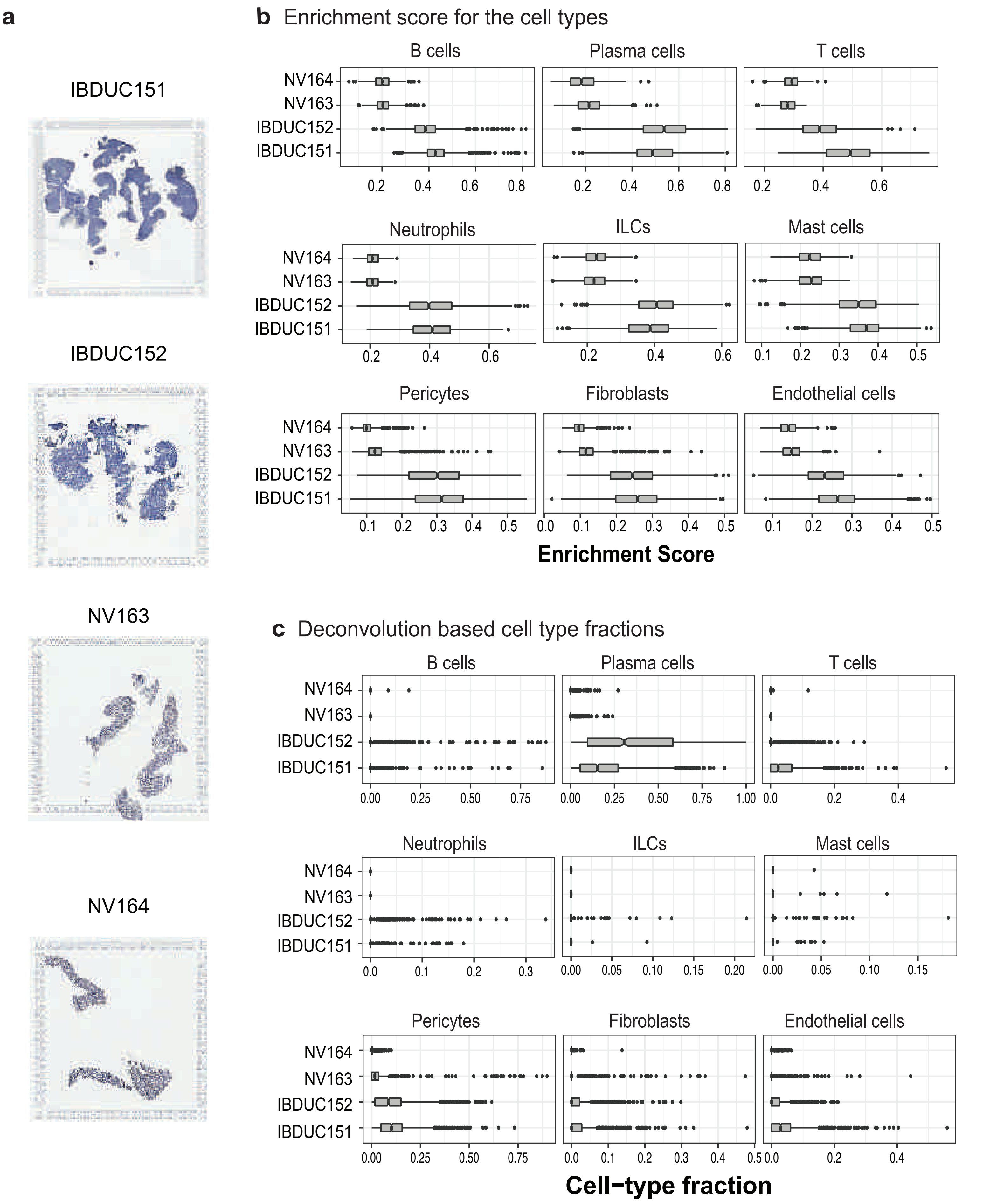

#### Supplementary Fig. 5: Enrichment of the epithelial subtypes from published dataset in the scRNA-seq derived cell types and clusters.

**a** Violin plots showing the enrichment of the epithelial subtypes (Smillie et.al.) within the scRNA-seq derived cell types from PC dataset.

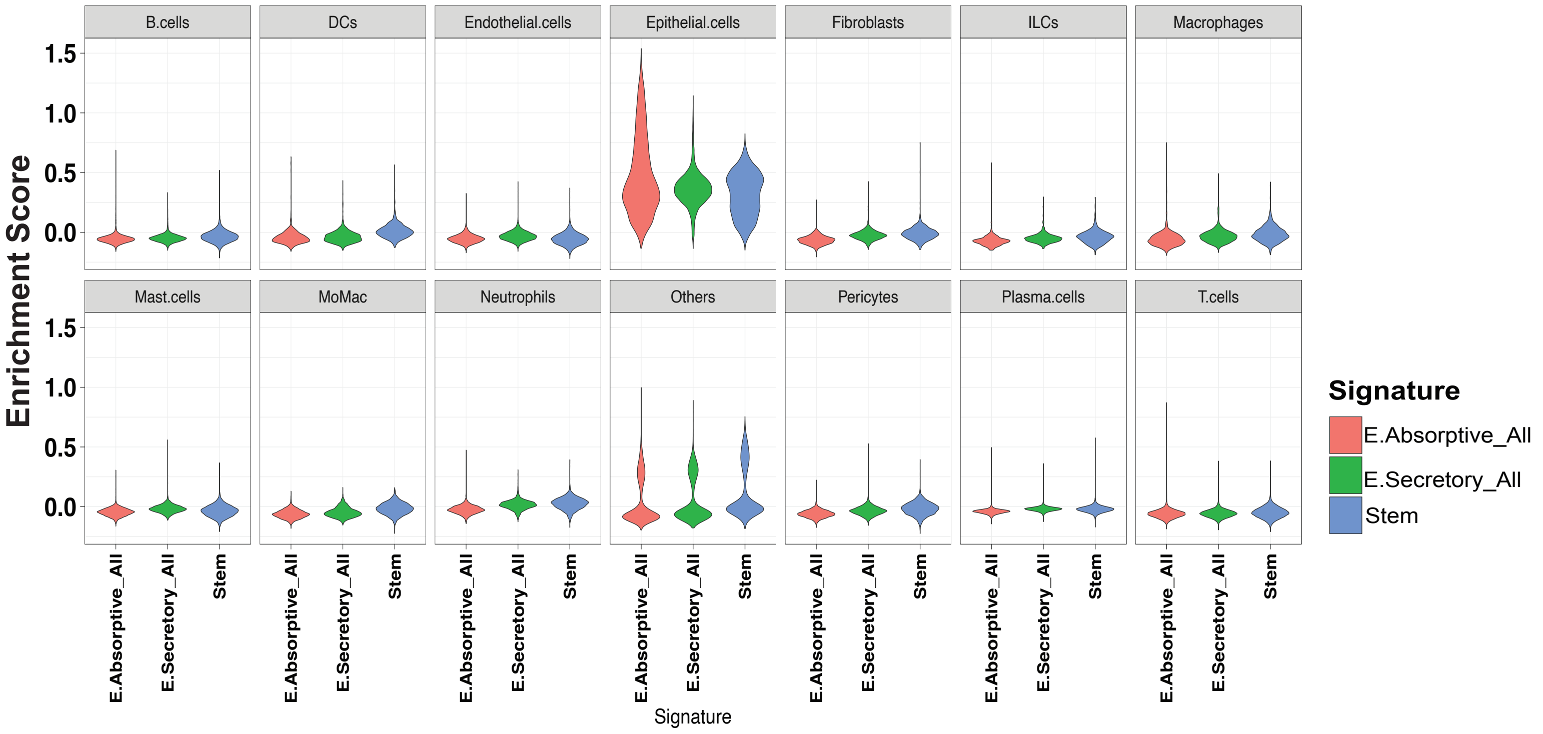


**b** Violin plots showing the enrichment of the epithelial cell subtypes (Smillie et.al.) within the scRNA-seq derived epithelial cell clusters from PC dataset.

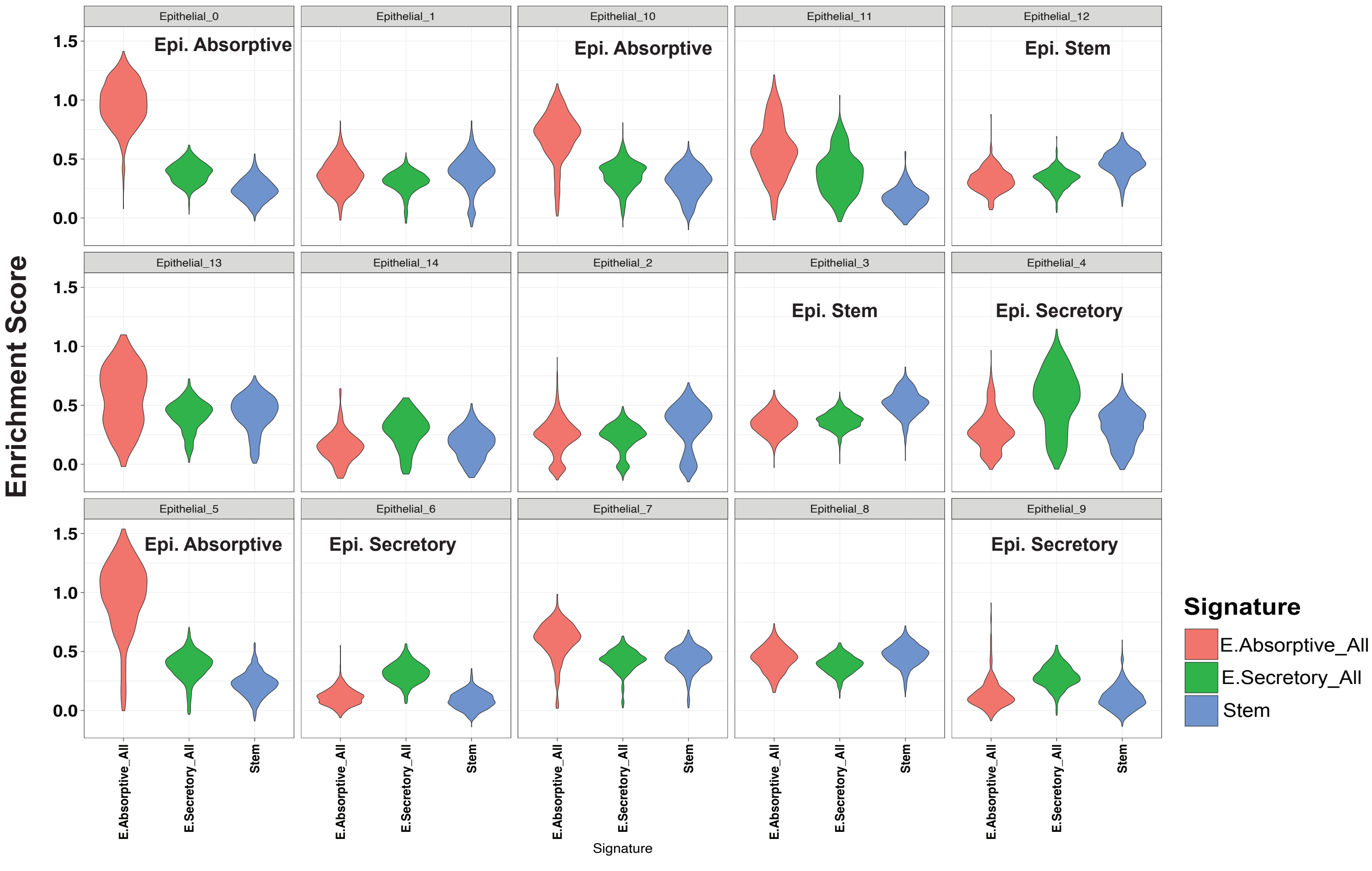

**Supplementary Fig. 6: CD14 (brown) immunohistochemical staining in the inflamed UC colonic sections compared to the uninflamed UC and NV colonic sections.**

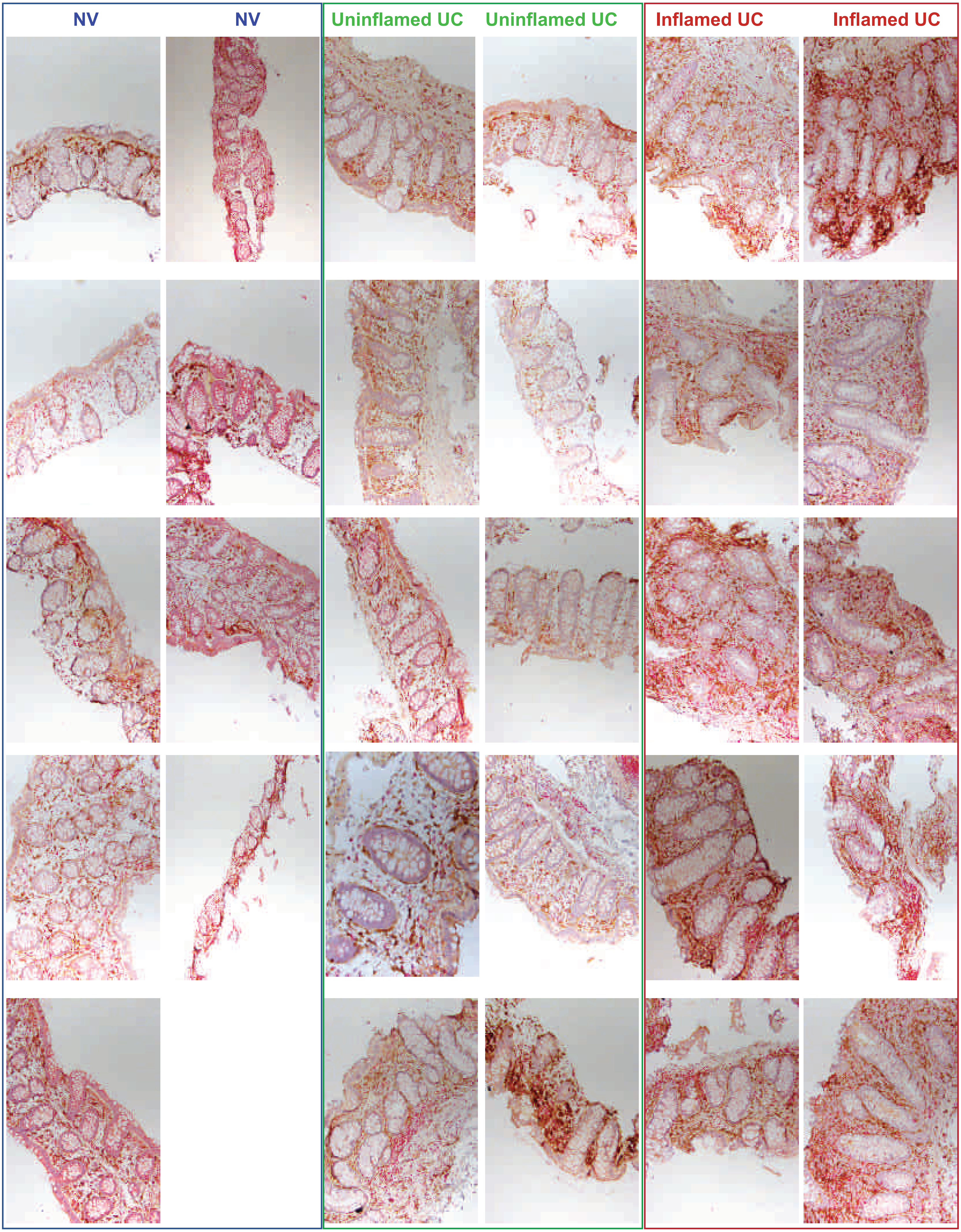

**Supplementary Fig. 7: Immunofluorescence staining of CD68⁺ macrophages in active UC patients and NV.**

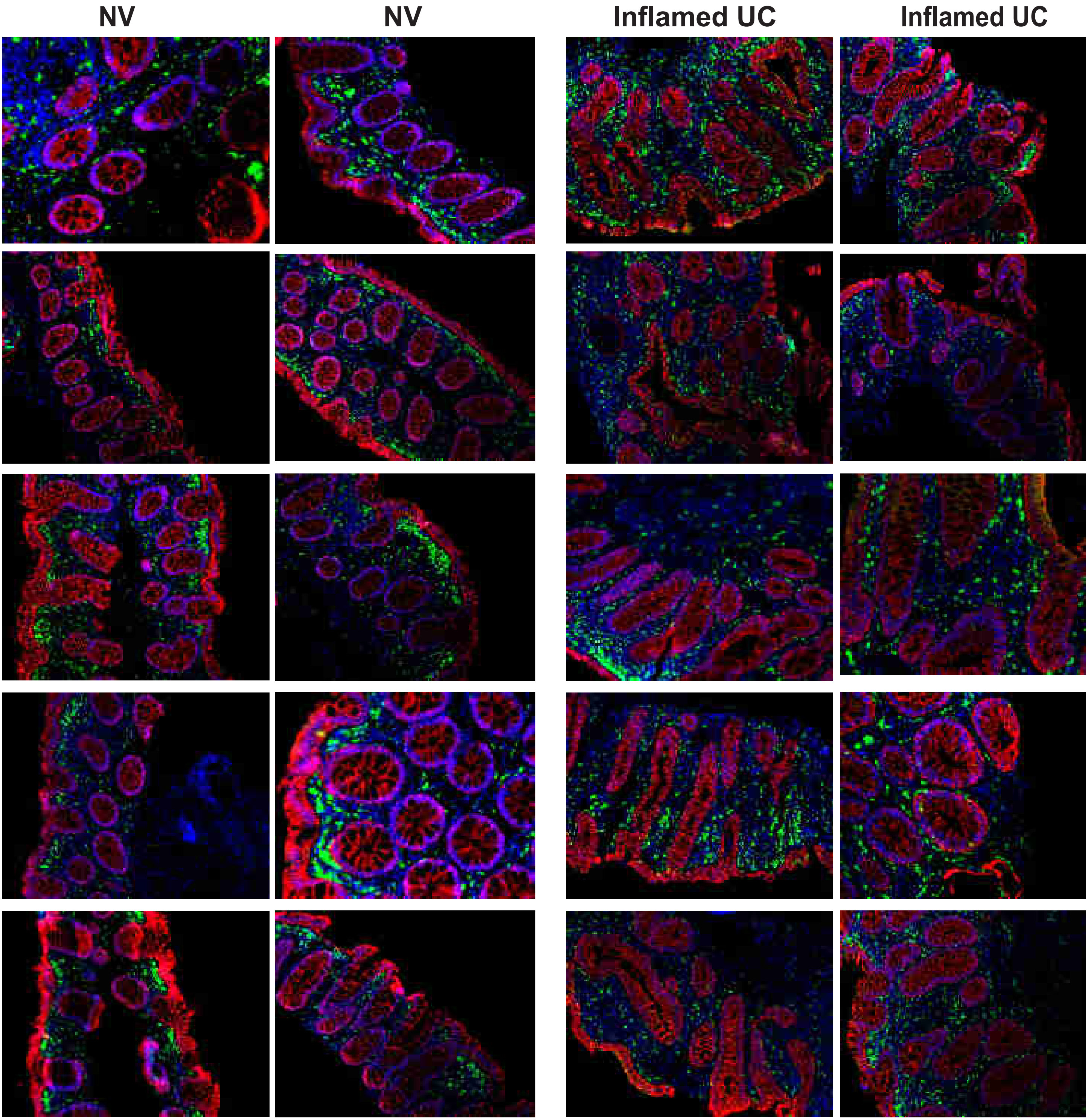

**Supplementary Fig. 8: Immunofluorescence staining of MPO⁺ neutrophils in NV and inflamed UC colonic tissues.**

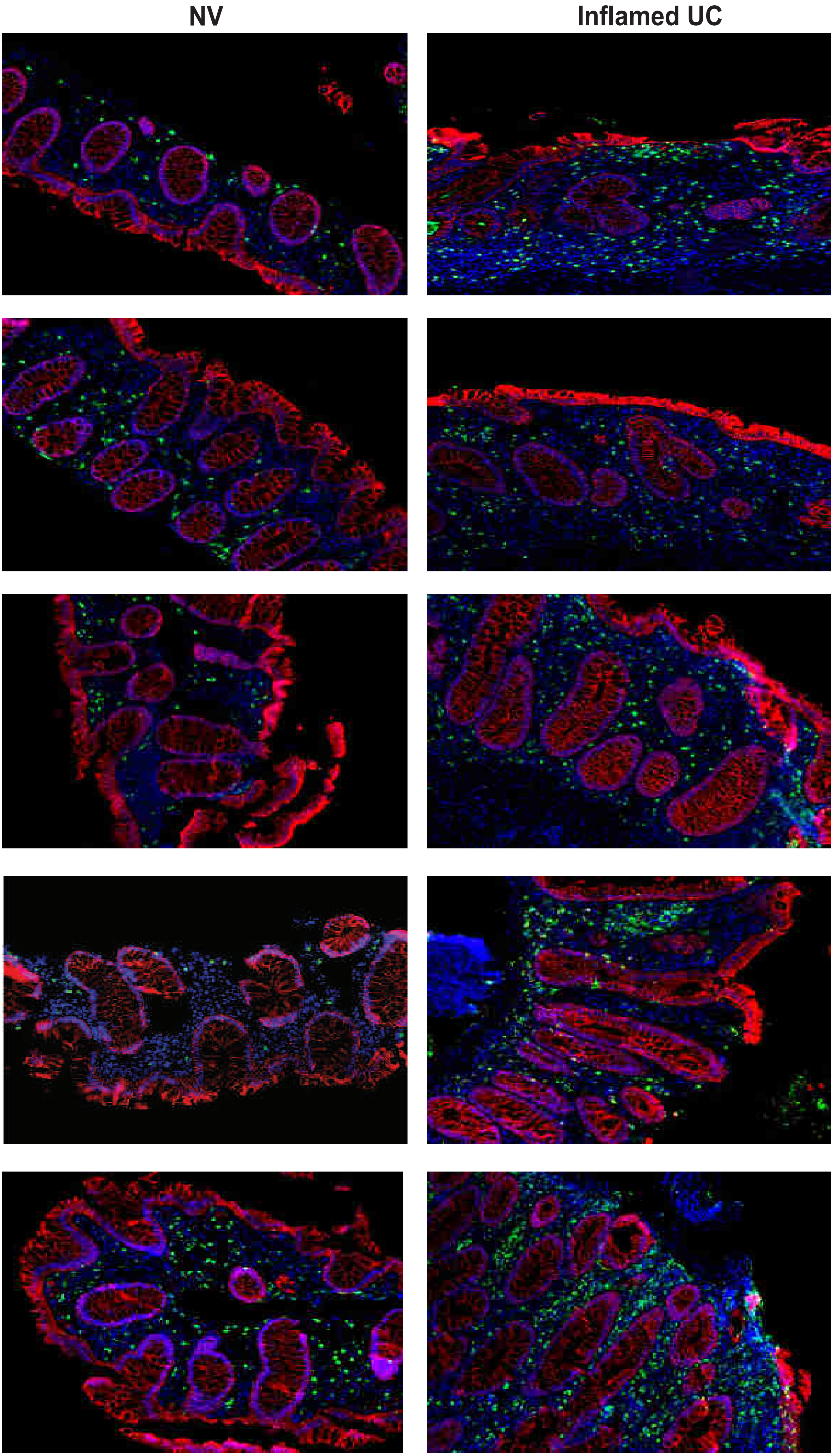

### Supplementary Fig. 9: Enrichment of treatment responsive gene signature on the cell types derived from sc-RNA sequencing data.

#### $IIC_{inf}$ bulk seq based treatment response signature

#### VC-3 cohort

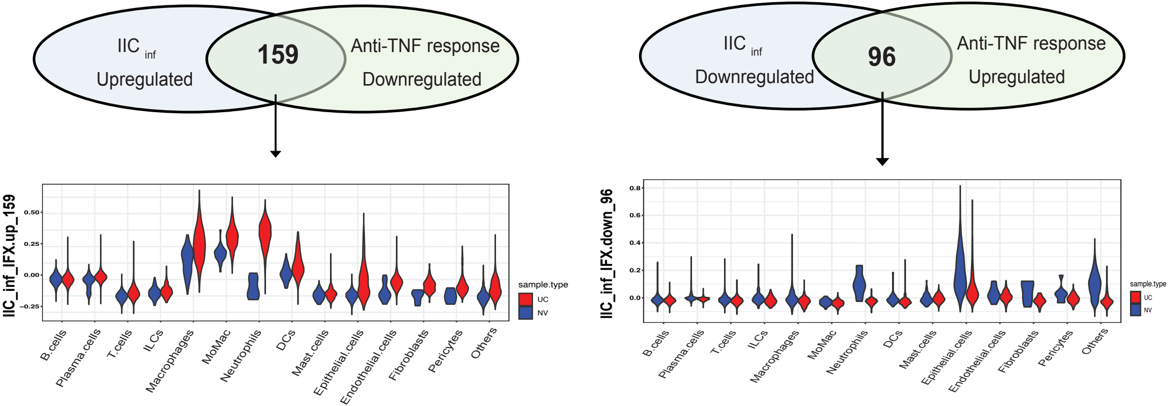

**Supplementary Fig. 10: Enrichment of treatment response associated genes in the scRNA-seq derived cell type markers.**
Fisher's exact test

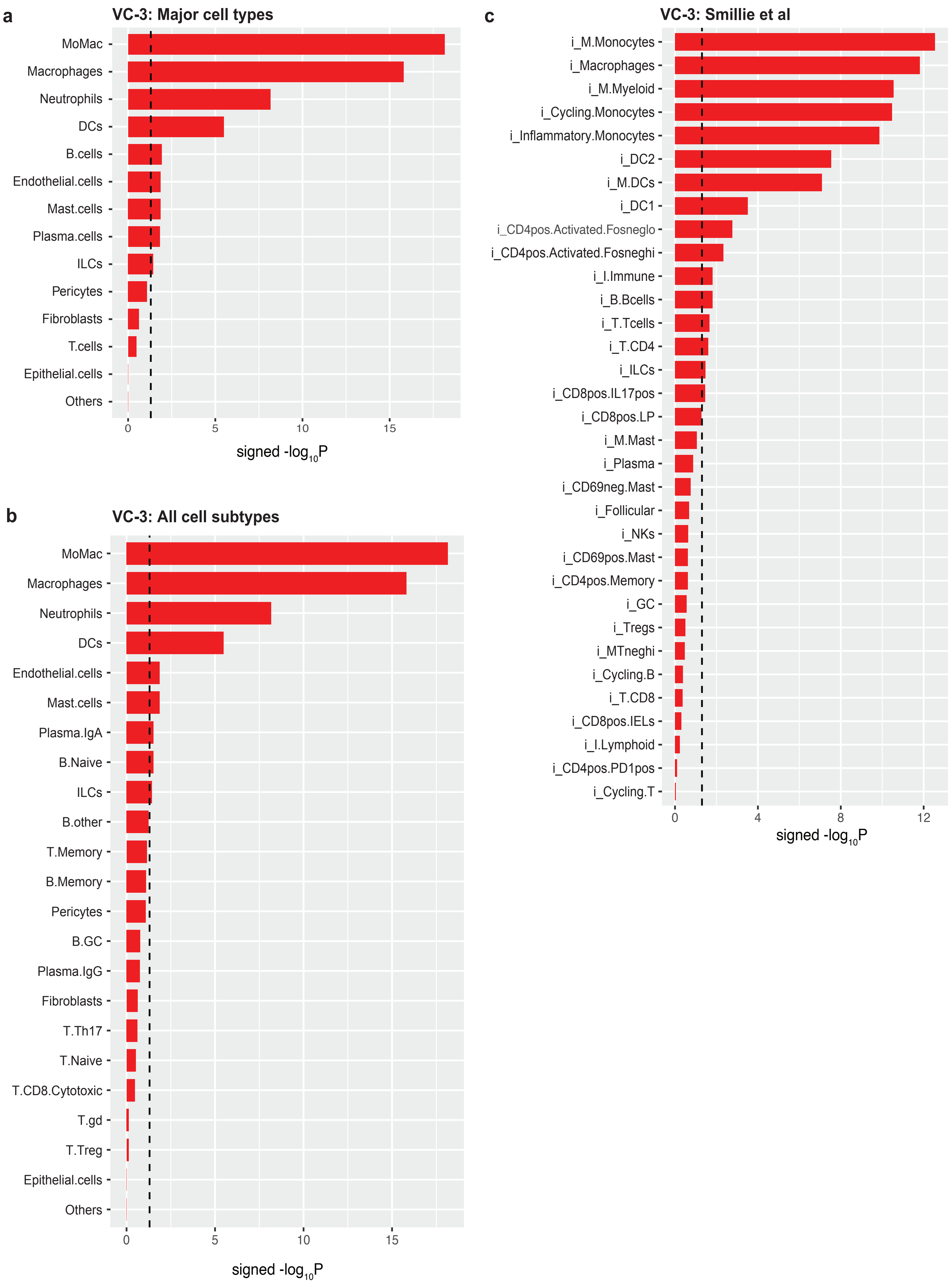

#### Supplementary Fig.11: Cell type association with anti-TNF therapy in VC-2 cohort.

**a.** Effect of anti-TNF (IFX) therapy on activity of cell type signatures in VC-2.

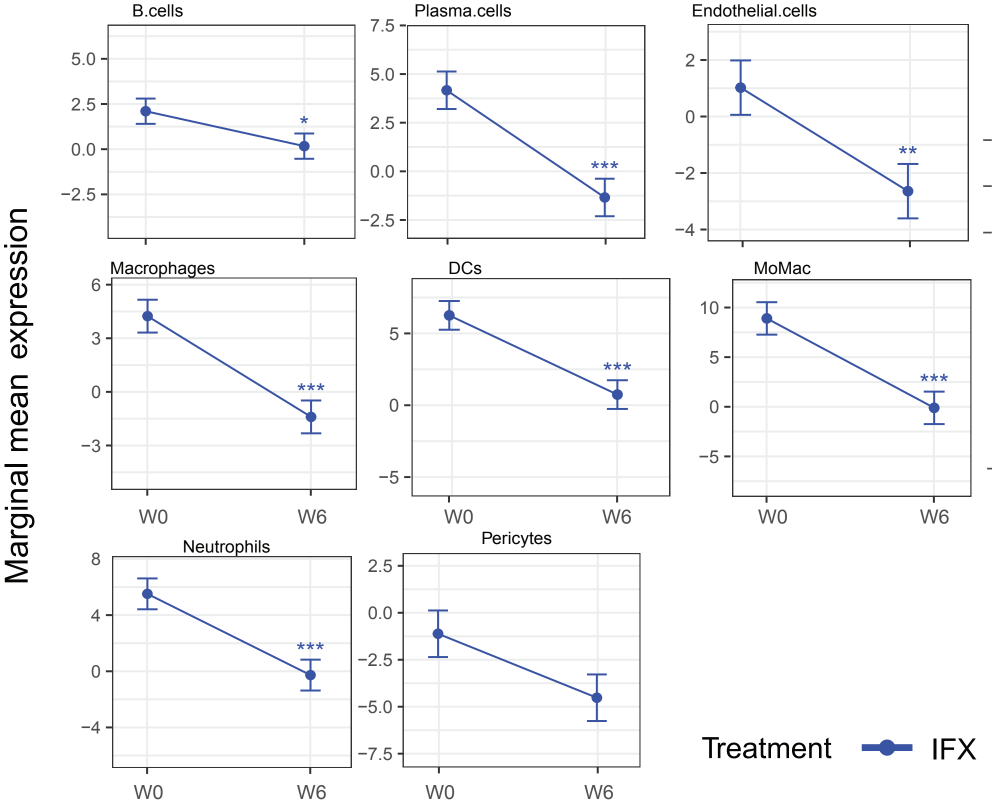


**b.** Effect of response to anti-TNF (IFX) on activity of cell type signatures in VC-2.

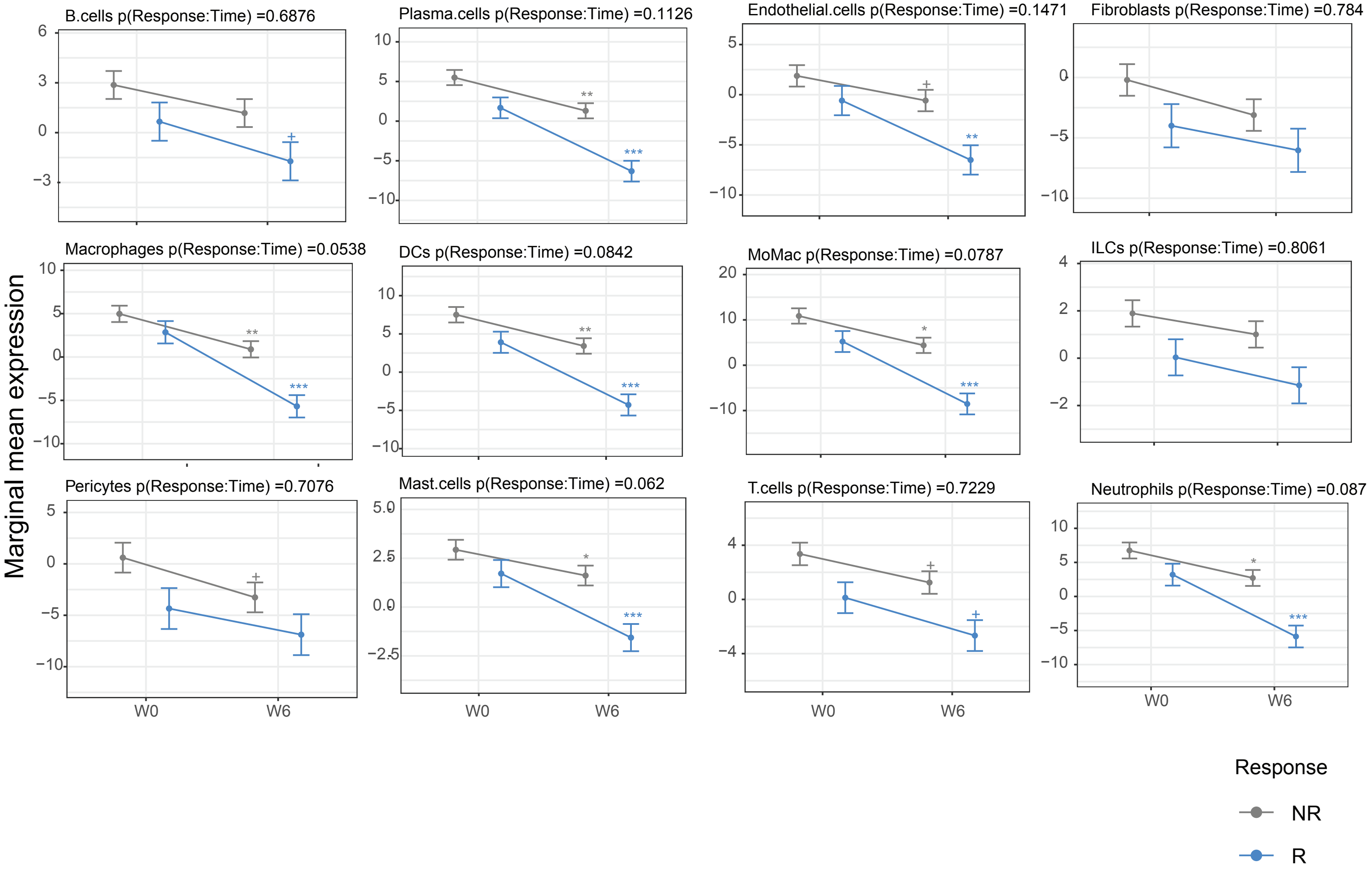

#### Supplementary Fig.12: Cell type association with anti-TNF therapy in VC-3 cohort.

**a.** Effect of anti-TNF (IFX) therapy on activity of cell type signatures in VC-3.

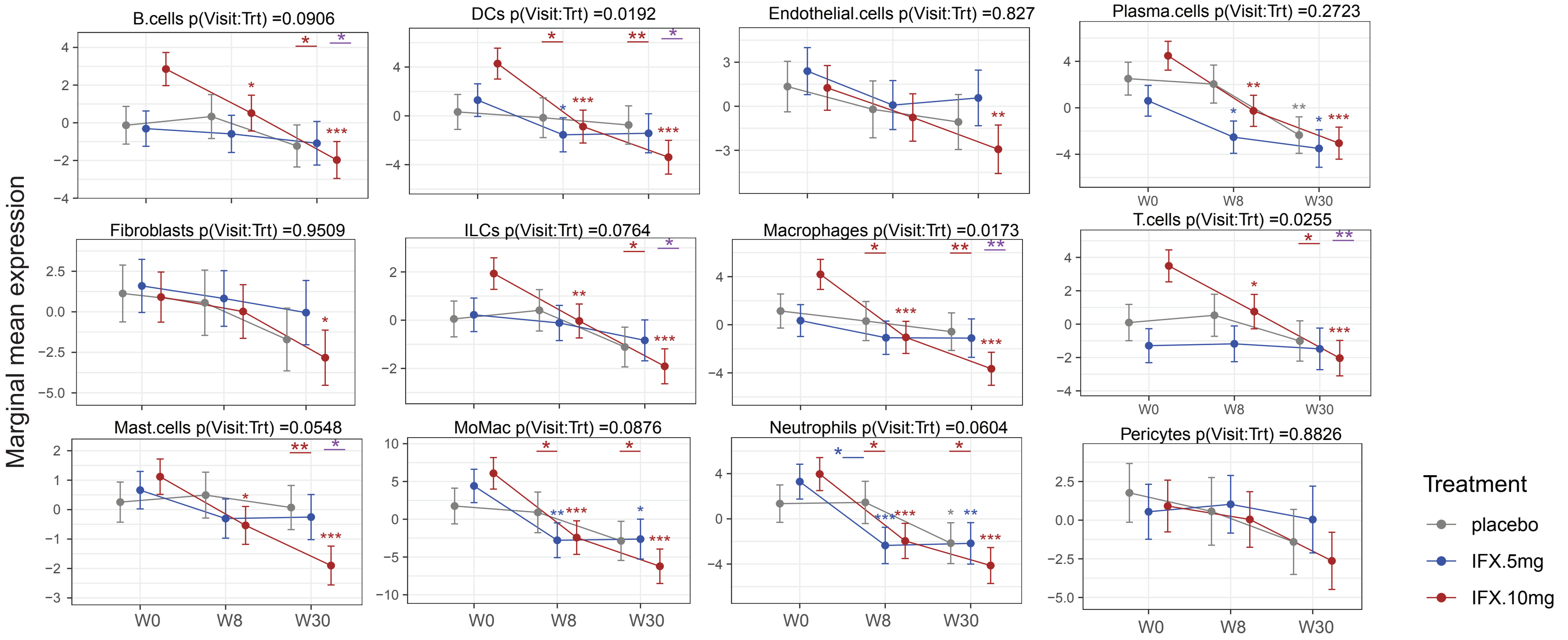


**b.** Effect of response to anti-TNF (IFX) on activity of cell type signatures in VC-3.

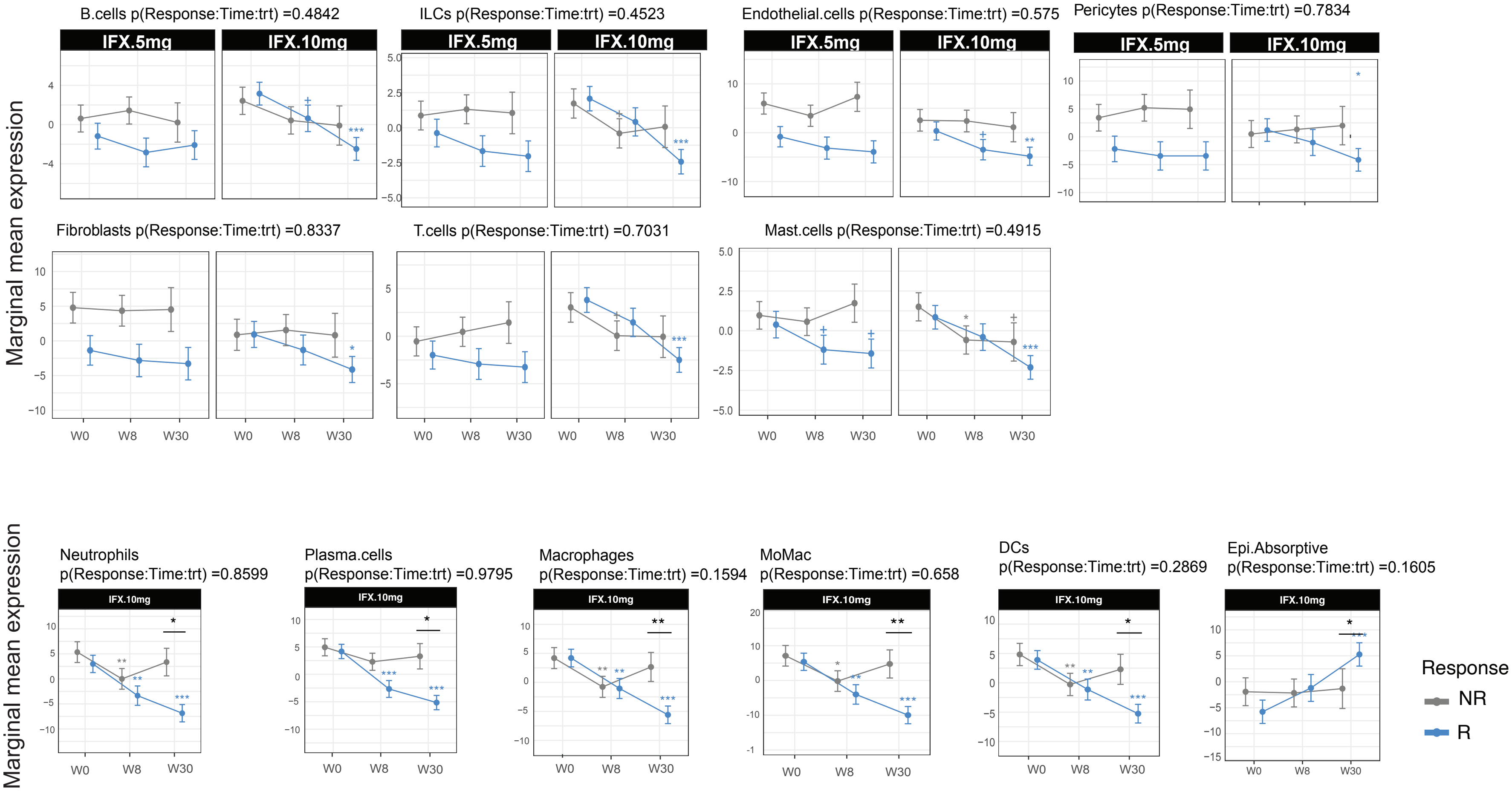

#### Supplementary Fig.13: Cell type association with anti-TNF therapy in VC-4 cohort.

**a.** Effect of anti-TNF (ADA) therapy on activity of cell type signatures in VC-4.

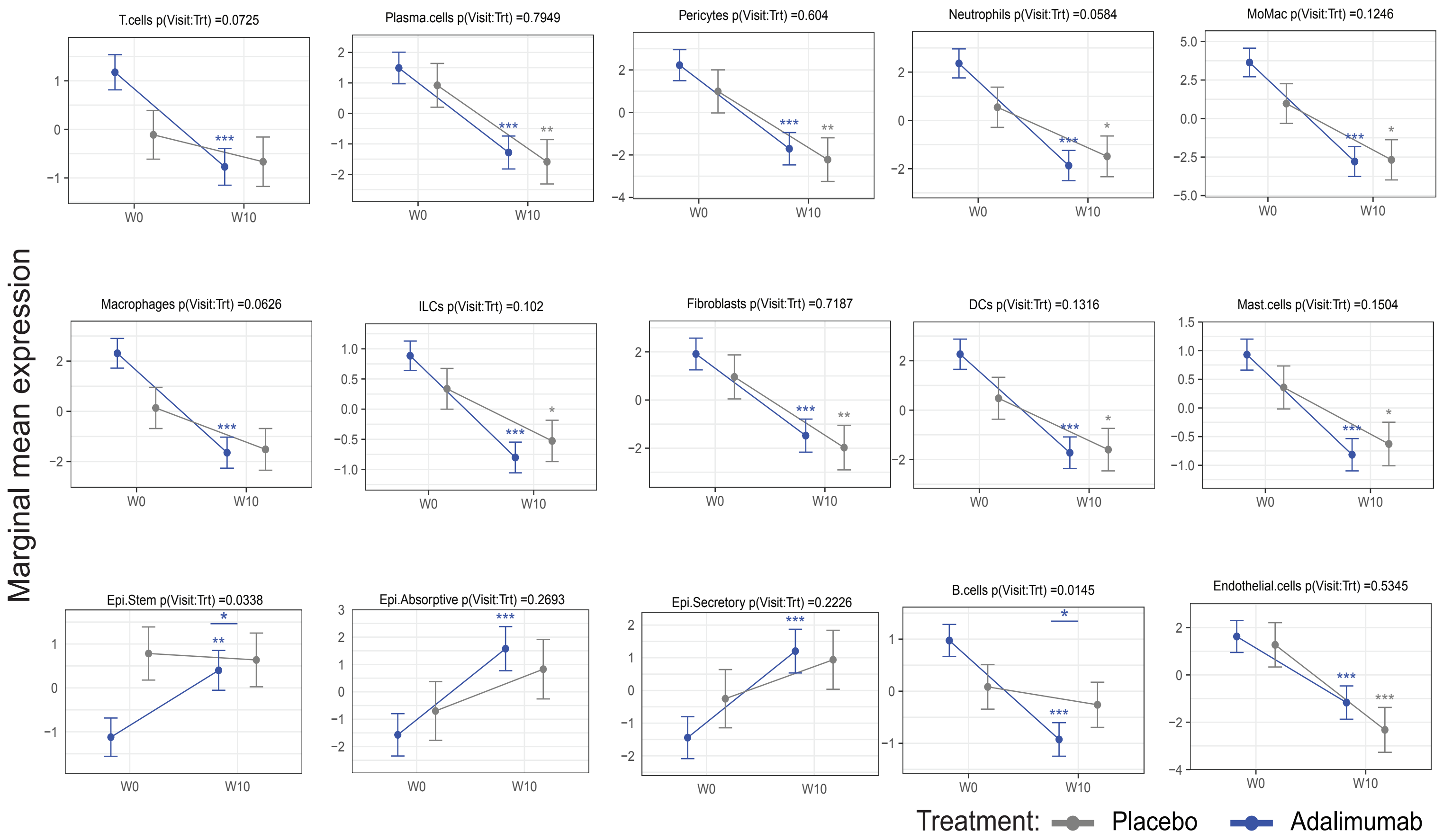


**b.** Effect of response (MCS-remission) to anti-TNF (ADA) on activity of cell type signatures in VC-4.

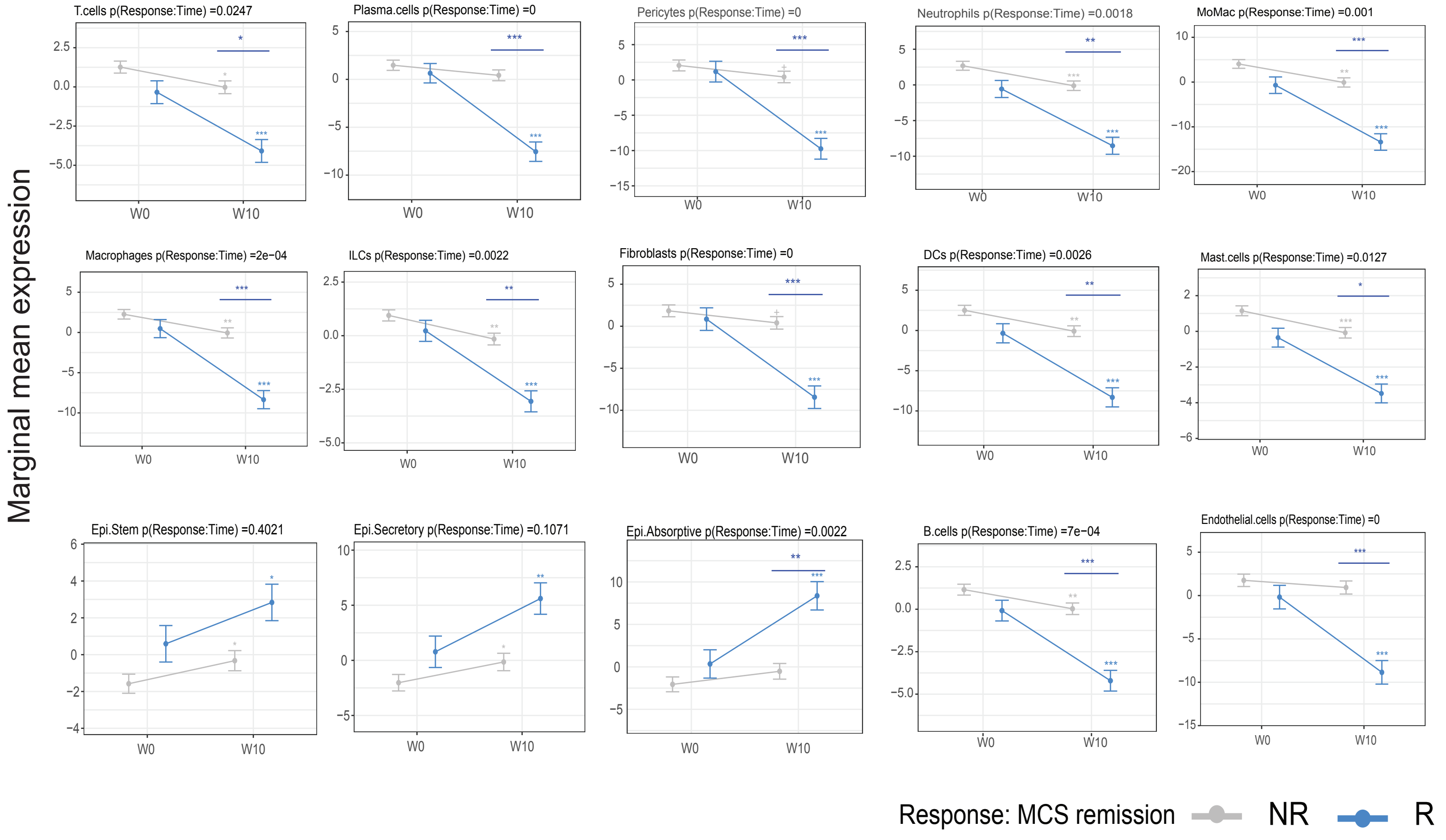

**Supplementary Fig. 14: Pathway enrichment analysis (KEGG database) associated with non-response to anti-TNF therapy.**

**a.** IIC-specific and IFX response signature- Upregulated

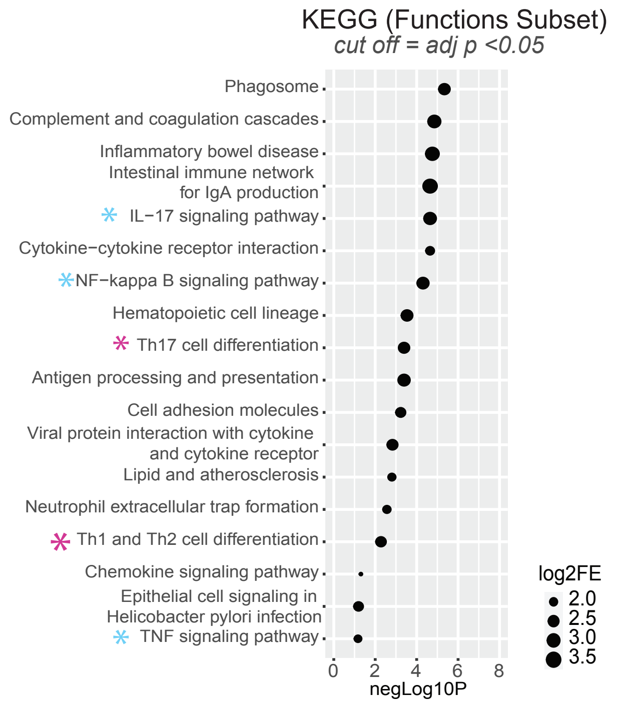


**b** Pathway enrichment analysis (KEGG 2021) on the leading edge genes from the cell types associated with non-response to anti-TNF therapy.

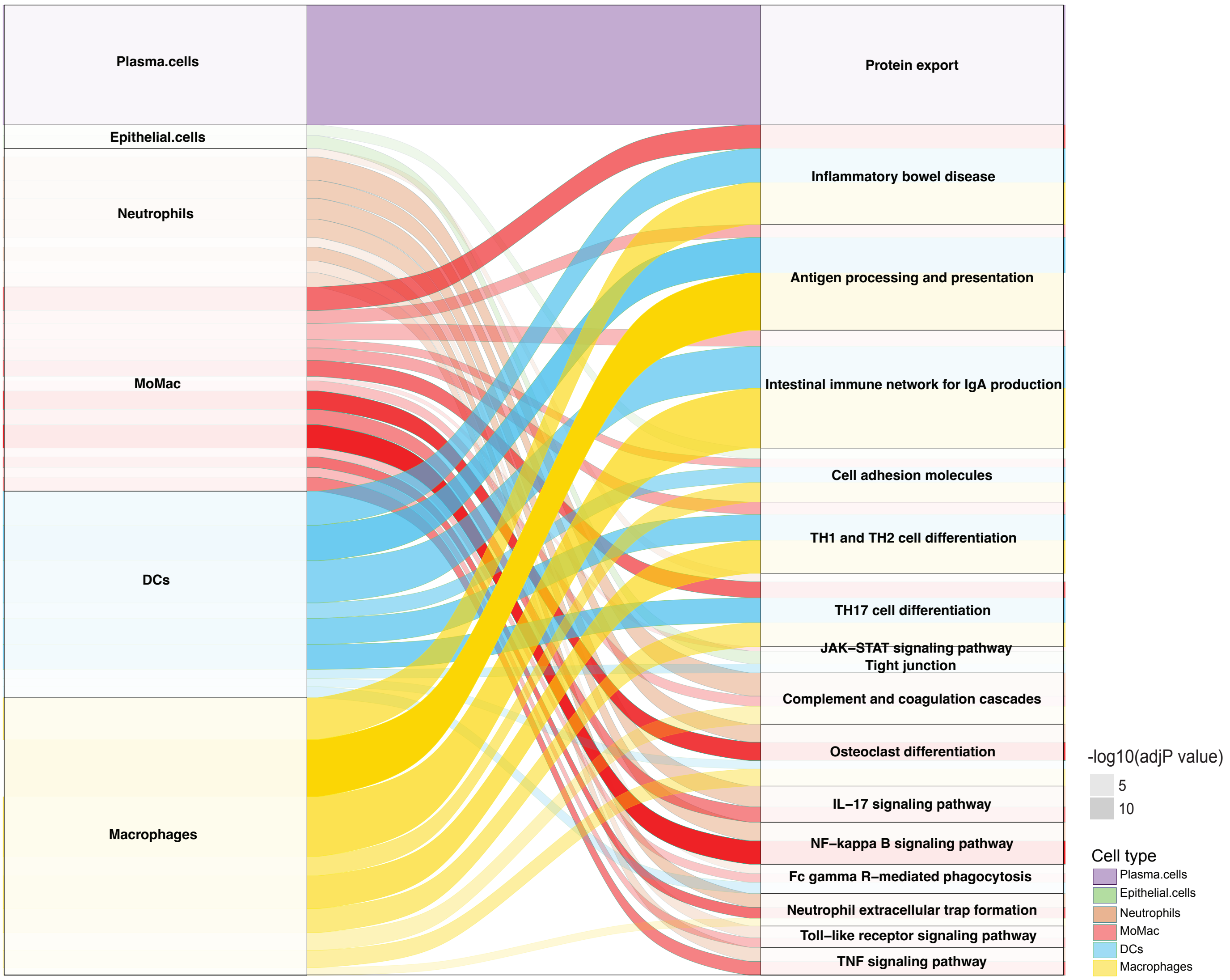
